# First *O*-demethylation activity in Arabidopsis specialized metabolism resolves the missing step in esculetin biosynthesis

**DOI:** 10.64898/2026.07.11.737633

**Authors:** Alicja Dobek, Clément Charles, Izabela Perkowska, Ryosuke Munakata, Jeremy Grosjean, Alain Hehn, Ewa Łojkowska, Anna Ihnatowicz, Alexandre Olry

## Abstract

Coumarins are phenylpropanoid-derived specialized metabolites that contribute to plant defence, shape plant–microbe interactions in the rhizosphere, and promote iron acquisition. In *Arabidopsis thaliana*, a model plant for iron-responsive coumarin metabolism, the enzymatic origin of the catecholic coumarin esculetin has long remained unresolved. Here we identify the first *O*-demethylation reaction in Arabidopsis specialized metabolism and show that 2-oxoglutarate- and Fe(II)-dependent dioxygenases catalyze scopoletin 6-*O*-demethylation to form esculetin. We designate these enzymes scopoletin 6-*O*-demethylases (S6ODs) and validate their activity through biochemical characterization, together with metabolomic profiling and independent loss-of-function mutant lines providing genetic evidence *in planta*. Disruption of S6OD activity remodels coumarin profiles and alters plant performance under limited iron availability, indicating that esculetin biosynthesis contributes to plant responses under these conditions. Our findings resolve the long-sought missing step in esculetin biosynthesis. It establishes *O*-demethylation as a previously unrecognized reaction in Arabidopsis specialized metabolism and suggest that 2OGD-mediated *O*-demethylation is recurrently recruited during evolution of plant metabolism, with implications for metabolic engineering and improvement of iron acquisition traits in crops.

## INTRODUCTION

Coumarins are specialized metabolites produced by numerous plant families (Nan et al., 2025). They function in diverse stress responses, including pathogen defence, modulation of root-associated microbial communities, and adaptation to iron (Fe) limitation (Schmid et al., 2014; Stassen et al., 2021; Zaynab et al., 2024). Although Fe is abundant in soils, its bioavailability is low under alkaline conditions, affecting large agricultural areas (Voges et al., 2019). Non-grass species secrete coumarins as an important strategy to improve Fe mobilization alongside their reduction-based uptake systems (Martín-Barranco et al., 2020). Key catecholic coumarins, including fraxetin, sideretin and esculetin contribute to Fe³⁺ solubilization and reduction in a pH-dependent manner (Schmid et al., 2014; Paffrath et al., 2024). In addition, coumarin homeostasis is regulated by transport processes, including Pleiotropic Drug Resistance 9 (PDR9, ABCG37) and NRT1/PTR FAMILY 7.2 (NPF7.2)-mediated export required for efficient root secretion (Fourcroy et al., 2014; Watanabe et al. 2026), indicating that both biosynthesis and transport are essential for effective Fe mobilization. Beyond their role in Fe acquisition, coumarins have also been implicated in plant responses to other abiotic stresses. For example, exogenous foliar application of esculin improved the yield quality of flax under salt stress (Khattab et al., 2024). Conversely, in cassava roots, accumulation of scopoletin, esculetin and their glycosides contributes to rapid post-harvest deterioration (Buschmann et al., 2000).

Coumarins are also involved in plant responses to biotic stress through their roles in interactions with microorganisms and defense against pathogens. Esculetin inhibits growth of *Phytophthora capsici* (Wang et al., 2021), and esculin, its glycosylated form, accumulates upon infection in multiple plant species (Fini et al., 2012; Witzel et al., 2024; Stringlis et al., 2019). Coumarins also restrict growth and biofilm formation of *Ralstonia solanacearum* and *Pseudomonas aeruginosa* and can shape rhizosphere microbial communities without suppressing beneficial strains (Yang et al., 2016; Zhang et al., 2018; Stringlis et al., 2018; Voges et al., 2019). It was shown that coumarins mediate complex ecological interactions in the rhizosphere, where certain rhizobacteria opportunistically utilize scopoletin and related compounds for colonization (Gu et al. 2026), while beneficial fungal endophytes can exploit host-derived scopoletin to enhance plant Fe nutrition (Van Dijck et al. 2025). It highlights a dual role of coumarins in both microbial competition and cooperation.

The coumarin biosynthetic pathway in *Arabidopsis thaliana* (hereafter Arabidopsis) has been extensively characterized (Kai et al., 2008; Fellenberg et al., 2012; Siwinska et al., 2018; Rajniak et al., 2018; Vanholme et al., 2019), revealing the enzymatic basis for formation of scopoletin, fraxetin and sideretin. However, the step leading to the catecholic coumarin esculetin remained unresolved. Three routes can be envisaged: 1) hydroxylation of caffeoyl-CoA followed by lactonization, 2) hydroxylation of umbelliferone, or 3) *O*-demethylation of scopoletin, the most abundant coumarin in Arabidopsis. The last possibility has been discussed in the literature (Schmid et al., 2014; Schmidt et al., 2014; Clemens and Weber, 2016), but no evidence for such activity *in planta* has been provided so far.

Demethylation occurs throughout the kingdoms of life and can be catalyzed by multiple enzyme classes, including cytochromes P450 (Lv et al., 2017), Rieske non-heme Fe-dependent oxygenases (Lee et al., 2018), 2-oxoglutarate and Fe(II)-dependent dioxygenases (2OGDs) (Hagel and Facchini, 2010), flavin adenine dinucleotide (FAD)-dependent oxidases (Shi et al., 2004), or coordinated multienzyme systems (Rahier et al., 2011). Depending on the chemical context, demethylation may target methyl groups directly attached to carbon or those linked *via* nitrogen (N) or oxygen (O). *O*-demethylation of small molecules is widespread in animals and microbes (Bertilsson et al., 2002; Coller et al., 2012; Harlington et al., 2025; Kasahara et al., 1996; Lalovic et al., 2004; Tripathi et al., 2026). In plants, however, *O*-demethylation of small molecules has been demonstrated only in two species and in both cases was catalyzed by 2OGDs (Berim and Gang, 2013; Hagel and Facchini, 2010). In Arabidopsis, no *O*-demethylation activity has previously been functionally demonstrated in specialized metabolism, and the enzymatic basis of this reaction remained unresolved.

2OGDs are widely distributed and constitute the second largest enzyme family in plants. They catalyze diverse oxidative reactions and participate in multiple metabolic pathways. The plant 2OGD superfamily can be divided into DOXA, DOXB and DOXC classes (Kawai et al., 2014). Most DOXC clades have undergone extensive expansion, vary highly across species and are associated with specialized metabolism. The number of DOXA and DOXB genes is relatively constant among species, whereas DOXC members vary considerably (Kawai et al., 2014). Despite their prevalence, many DOXC enzymes remain functionally uncharacterized. Comparative analyses in lineages such as *Apiaceae* further suggest DOXC diversification, proceeded through repeated gene duplication and functional divergence, reflecting a dynamic evolutionary history of coumarin-associated metabolism (Huang et al. 2024).

Here, we investigate a group of 2OGDs from Arabidopsis belonging to the DOXC52 subclade (Kawai et al., 2014). This clade also includes three characterized genes: thebaine 6-*O*-demethylase (T6ODM) and codeine *O*-demethylase (CODM) from opium poppy, which catalyze successive *O*-demethylation steps in morphine biosynthesis (Hagel and Facchini, 2010), as well norcoclaurine synthase (CjNCS1) in *Coptis japonica* catalyzing the condensation of dopamine and 4- hydroxyphenylacetaldehyde to form norcoclaurin (Minami et al., 2007). Arabidopsis and *Oryza sativa* encode five and thirteen DOXC52 members, respectively, yet do not produce isoquinoline alkaloids, and the biochemical functions of these enzymes remain unclear. The identification of 2OGD-dependent *O*-demethylation in opium poppy alkaloid biosynthesis, and later in flavone metabolism in basil (Berim et al., 2013), raised the question whether similar reactions occur in other plant species and metabolic pathways. Recent *in vitro* biochemical evidence further indicates that 2OGD-mediated *O*-demethylation can arise in phylogenetically distinct enzyme lineages, including the substrate-dependent bifunctional activity reported for FcDOH2 from *Fraxinus chinensis*, which catalyzes both *O*-demethylation and hydroxylation reactions of coumarins *in vitro* (Kong et al. 2026). However, whether this *O*-demethylation activity contributes to coumarin metabolism under physiological conditions in plants remains unknown.

In this work, we functionally characterize members of the Arabidopsis DOXC52 clade, designated AtOD1–AtOD5, and investigate their homologs across the plant kingdom. Using a heterologous *Escherichia coli* expression system, we demonstrate that members of this clade catalyse the *in vitro* conversion of scopoletin to esculetin through 6-*O*-demethylation and confirm their function *in planta*. We designate these enzymes scopoletin 6-*O*-demethylases (S6ODs). Our findings provide the first evidence of *O*-demethylation in specialized metabolism of the widely studied model plant Arabidopsis and resolve the long-standing missing step in esculetin biosynthesis (**Figure 1**). Loss of S6OD activity remodels coumarin profiles and alters plant performance under Fe limitation, linking this reaction to Fe acquisition physiology. Beyond completing our understanding of the Arabidopsis coumarin biosynthetic pathway, the identification of S6ODs highlights *O*-demethylation as a key step in coumarin metabolism *in planta*, with implications for both plant biology and metabolic engineering.

**Figure 1.**
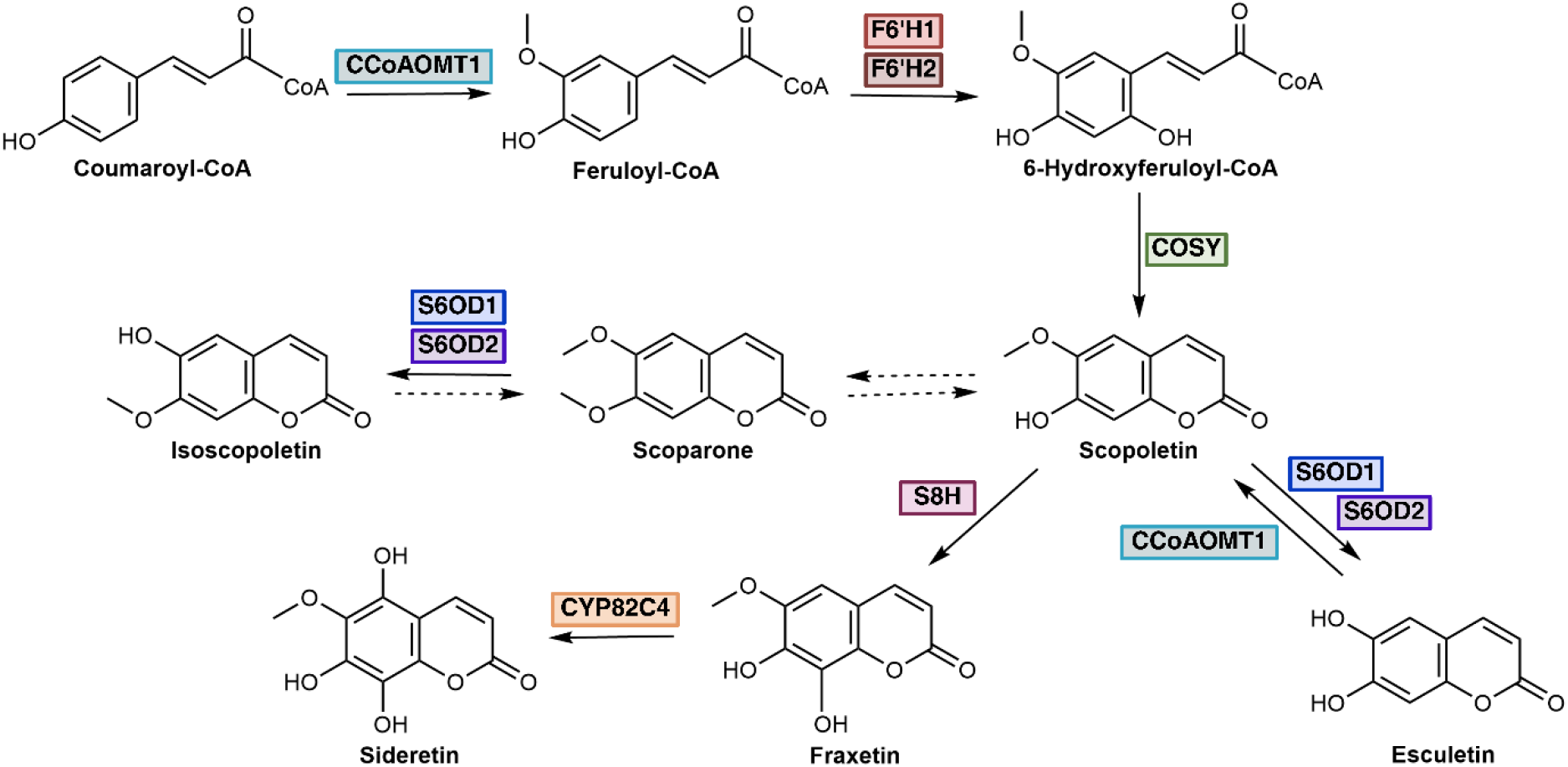
Schematic representation of the coumarin biosynthetic pathway in *Arabidopsis thaliana*. CCoAOMT1 - Caffeoyl-CoA *O*-methyltransferase 1, required for feruloyl-CoA biosynthesis (Fellenberg *et al*., 2012); F6’H1 - Feruloyl CoA ortho-hydroxylase 1 and F6’H2 - Feruloyl CoA ortho-hydroxylase 2, involved in scopoletin biosynthesis (Kai *et al*., 2009); COSY – Coumarin Synthase, catalyzing the biosynthesis of scopoletin through trans–cis isomerization followed by a lactonization (Vanholme et al., 2019); S6OD1 – Scopoletin 6-*O*-demethylase 1, and S6OD2 - Scopoletin 6-*O*-demethylase 2, responsible for esculetin and isoscopoletin production; S8H – Scopoletin 8-hydroxylase and CYP82C4 – Cytochrome p450, family 82, subfamily c, polypeptide 4, which catalyze fraxetin and sideretin biosynthesis, respectively (Siwinska et al., 2018; Rajnjak et al., 2018). Glucosylated forms are not shown for clarity. Sideretin occurs in reduced and oxidized forms.

**Table 1.**
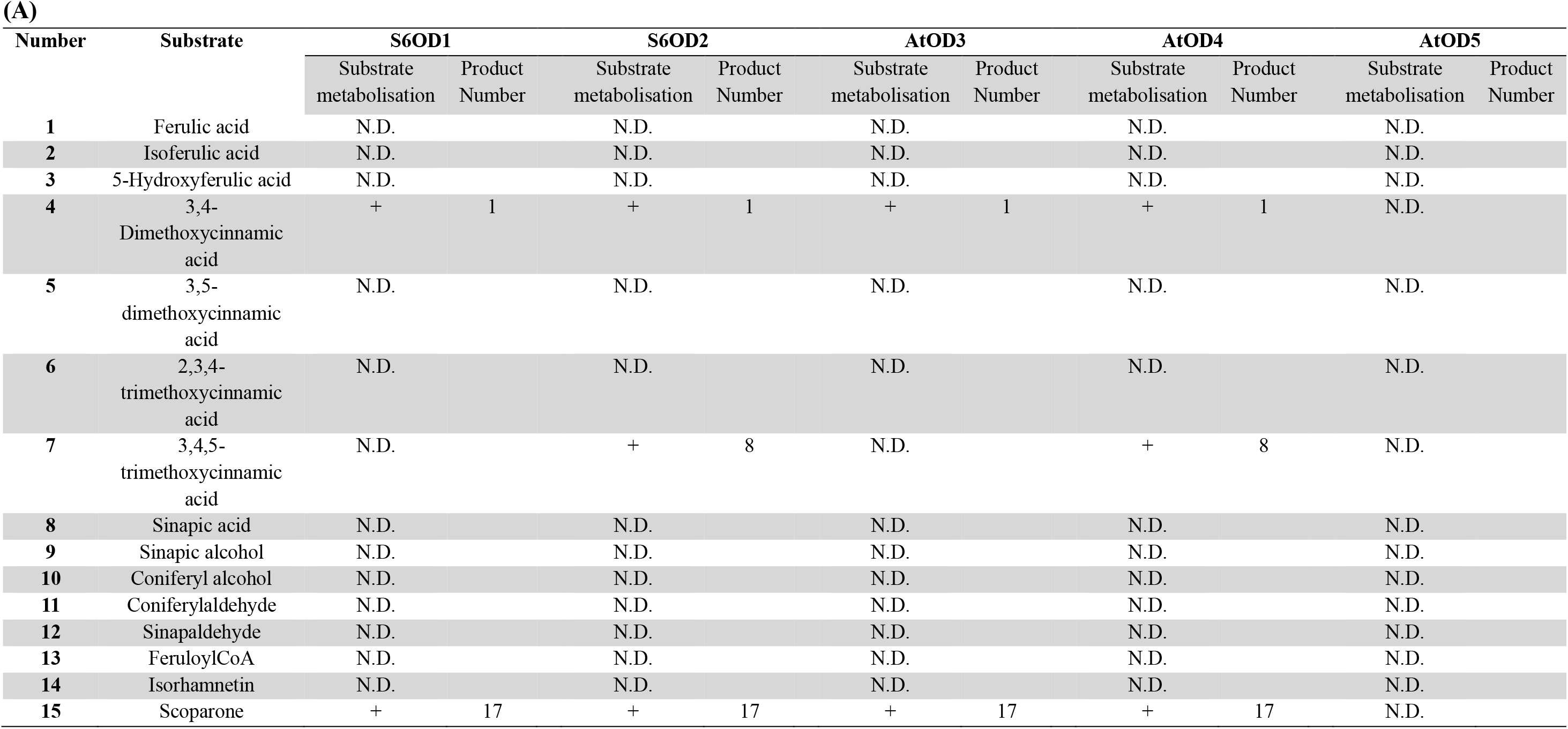
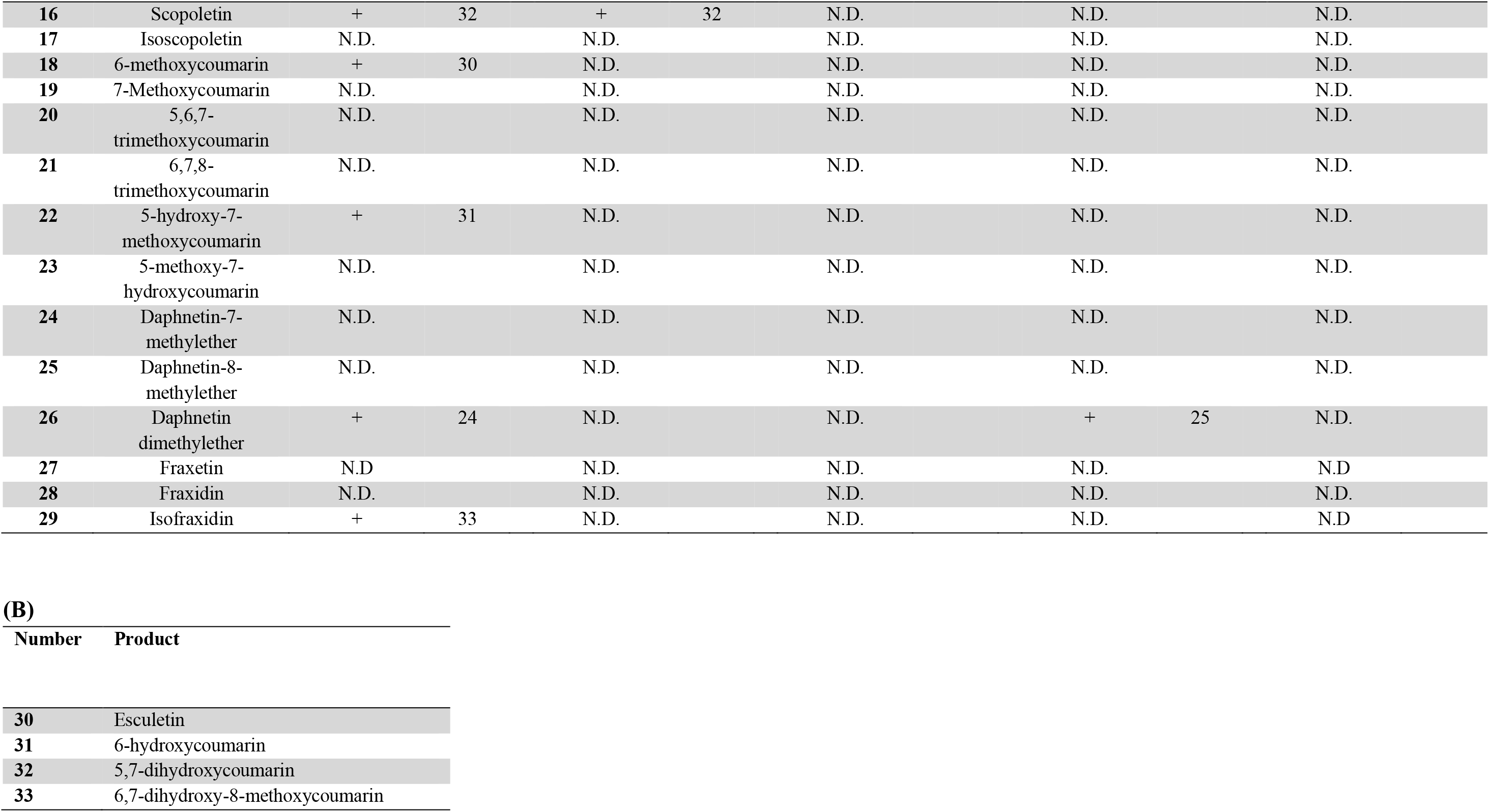
*O*-demethylated substrate specificity of the DOXC52 subclass from *Arabidopsis thaliana*. Phenylpropanes **(1–12)**, CoA ester: feruloyl-CoA **(13)**, flavonoid: isorhamnetin **(14)**, and coumarins **(15–29)** were tested as potential substrates **(A)** of S6OD1, S6OD2, AtOD3, AtOD4, and AtOD5. Reaction mixtures (100 µL) contained purified enzyme, 500 µM α-ketoglutarate, 500 µM ascorbic acid, 5 mM FeSO₄, and 200 µM *O*-methylated substrate, and were incubated for 20 h in a mixed buffer system (0.1 M acetic acid, 0.1 M Tris and 40 mM imidazole) at optimal temperature and pH (Supplementary figures). *O*-demethylated products are indicated either by the number of the tested substrate (**1–29**) or by the product numbers **(B)**: esculetin (**30**); 6-hydroxycoumarin (**31**); 5,7-dihydroxycoumarin (**32**); and 6,7-dihydroxy-8-methoxycoumarin (**33**). N.D., not detected. The chemical structures of tested substrates are shown in Supplementary Figures. Independent triplicate reactions yielded identical results.

## RESULTS

### Identification of *O*-demethylases candidates and phylogenetics

BLAST analysis of the T6ODM primary sequence enabled the identification of the closest dioxygenases within the Arabidopsis 2OGD family. Five candidate enzymes were selected based on a minimum and maximum sequence identity and similarity of 63-85 % and 81-91 %, respectively (**Supplementary Table S1**). For phylogenetic analysis of AtOD1-5, we searched for the homologs of AtOD1-5 in different plant genomes by sequence similarity. This analysis found that only in genomes of angiosperm species but not in those of a liverwort, a moss, and a green algae (**Supplementary File 1 and 2**), being consistent with the previously reported distribution of the DOXC subfamily (Kawai et al., 2014). In a phylogenetic tree of the DOXC52 subfamily (**Figure 2**), AtOD1-5 formed a clade with sequences from Fabids and Malvids (*Glycine max*, *Vigna unguiculata*, *Fragaria* × *ananassa*, *Theobroma cacao*, *Gossipium hirsutum*, *Citrus sinensis*, *Populus tricocarpa*), suggesting this orthologous group arose before the divergence of these taxa. From a more localized viewpoint, AtOD1-5 were found to be adjacent to each other in the phylogenetic tree, with all sequences from Malvales, the neighbor to Brassicales, being outside the AtOD1-5 cluster. This relationship suggests that the five paralogs have occurred through Brassicales-specific gene duplication events. Multiple sequence alignment of the five selected candidates together with all enzymatically characterized *O*-demethylases revealed the presence of the conserved catalytic motif typical of 2OGDs—His238-Xaa-Asp/Glu240-(Xaa)n-His295—which coordinates Fe(II) to form the catalytic triad. This motif is conserved in all aligned sequences, except in AtOD5, where Asp240 is substituted by Glu. In addition, residues involved in succinate, α-ketoglutarate, and substrate recognition are highly conserved (**Supplementary Figure S1**).

**Figure 2.**
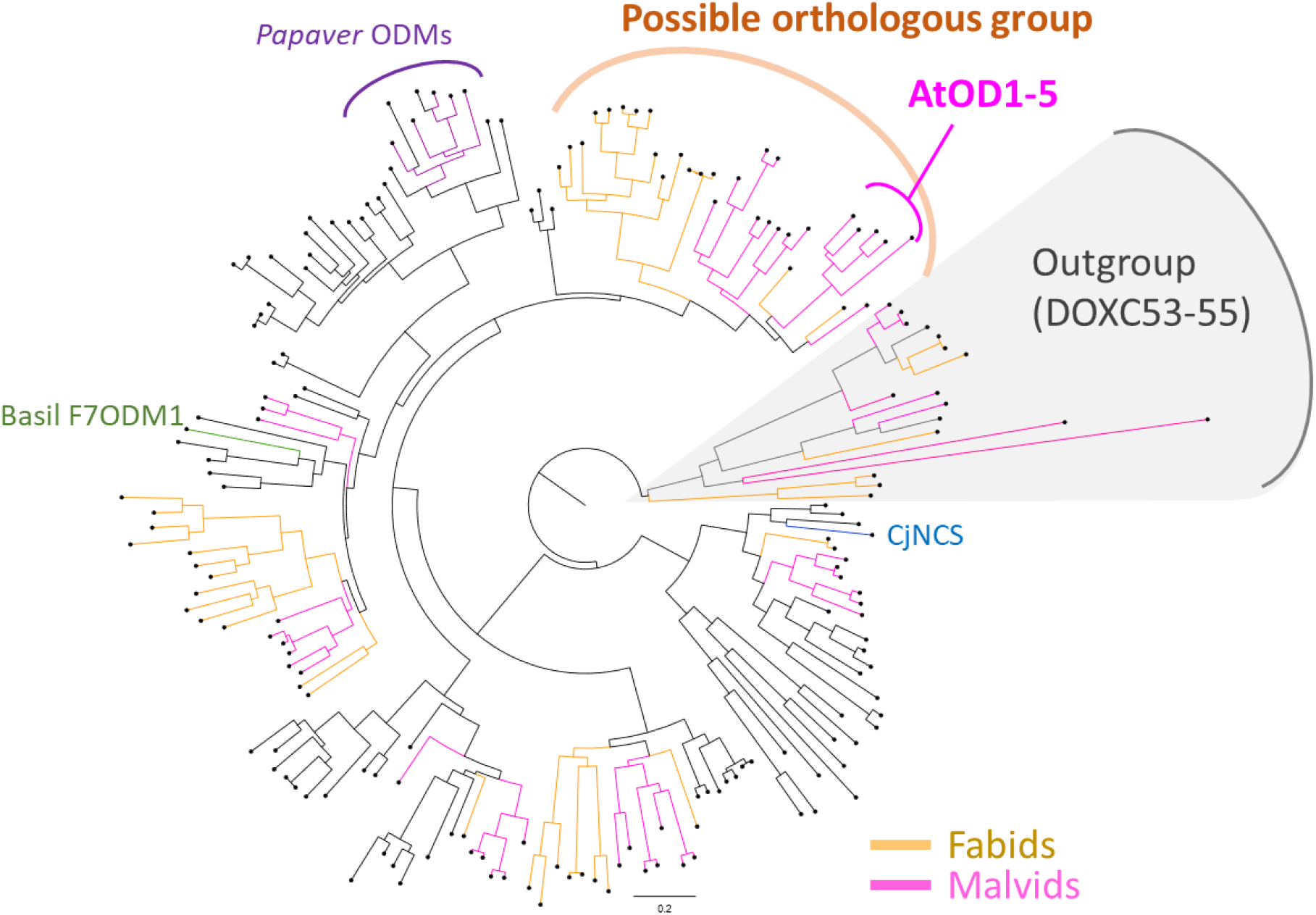
A phylogenetic tree of the DOXC52 subfamily. A maximum likelihood-based phylogenetic tree was constructed for the DOXC52 subfamily using DOXC53, 54, and 55 sequences as outgroups, based on MAFFT alignment and IQ-TREE inference (JTT+F+I+G4 model). The scale bar indicates an amino acid substitution rate per site of 0.20. The proteins from Fabids and Malvids are highlighted by orange and magenta, respectively. The previously-reported plant *O*-demethylases from *Papaver somniferum* and *Ocimum basilicum*, as well as norcoclaurine synthase catalyzing the condensation of dopamine and 4- hydroxyphenylacetaldehyde to form norcoclaurin in *Coptis japonica*, are shown in purple, green, and blue, respectively. The input sequences were listed in **Supplementary File 1.** The detailed phylogenetic tree, where each sequence name, and the results of the bootstrap test and the Shimodaira-Hasegawa approximate likelihood-ratio test are visualized, is given in **Supplementary File 2.**

### *In vitro* substrate specificity of Arabidopsis *O*-demethylases (AtODs)

Prior to enzymatic assays, the secondary structure of renatured enzymes was analyzed by circular dichroism spectroscopy. The S8H enzyme was used as a control (**Supplementary Figure S2 and Table S2**). Experimental spectra were compared with secondary structure predictions generated using JPred software. Both experimental data and predictions were consistent with a general folding characteristic of 2OGD family and were comparable to that of S8H (Siwinska et al., 2018).

To gain deeper insight into the substrate specificity of AtODs enzymes, we conducted a comprehensive screening using a panel of methoxylated compounds, comprising 12 phenylpropanes, 19 coumarins, one flavonol, and one CoA ester (**Supplementary Figure S3 and S4**). Enzymatic assays were carried out under optimal conditions (∼26 C, pH 6.0) (**Supplementary Figure S5 and S6**) in the presence of saturating α-ketoglutarate concentrations. These experimental parameters are in line with previously reported optimal conditions (Kai et al., 2008; Siwińska et al., 2018). Reactions were incubated overnight to enhance the detection of weak catalytic activities. Products were identified by UHPLC-DAD. The structural identity of the products was confirmed via MS² analysis and compared with authentic standards (**Figure 3 and Supplementary Figures S10 to S25**).

**Figure 3.**
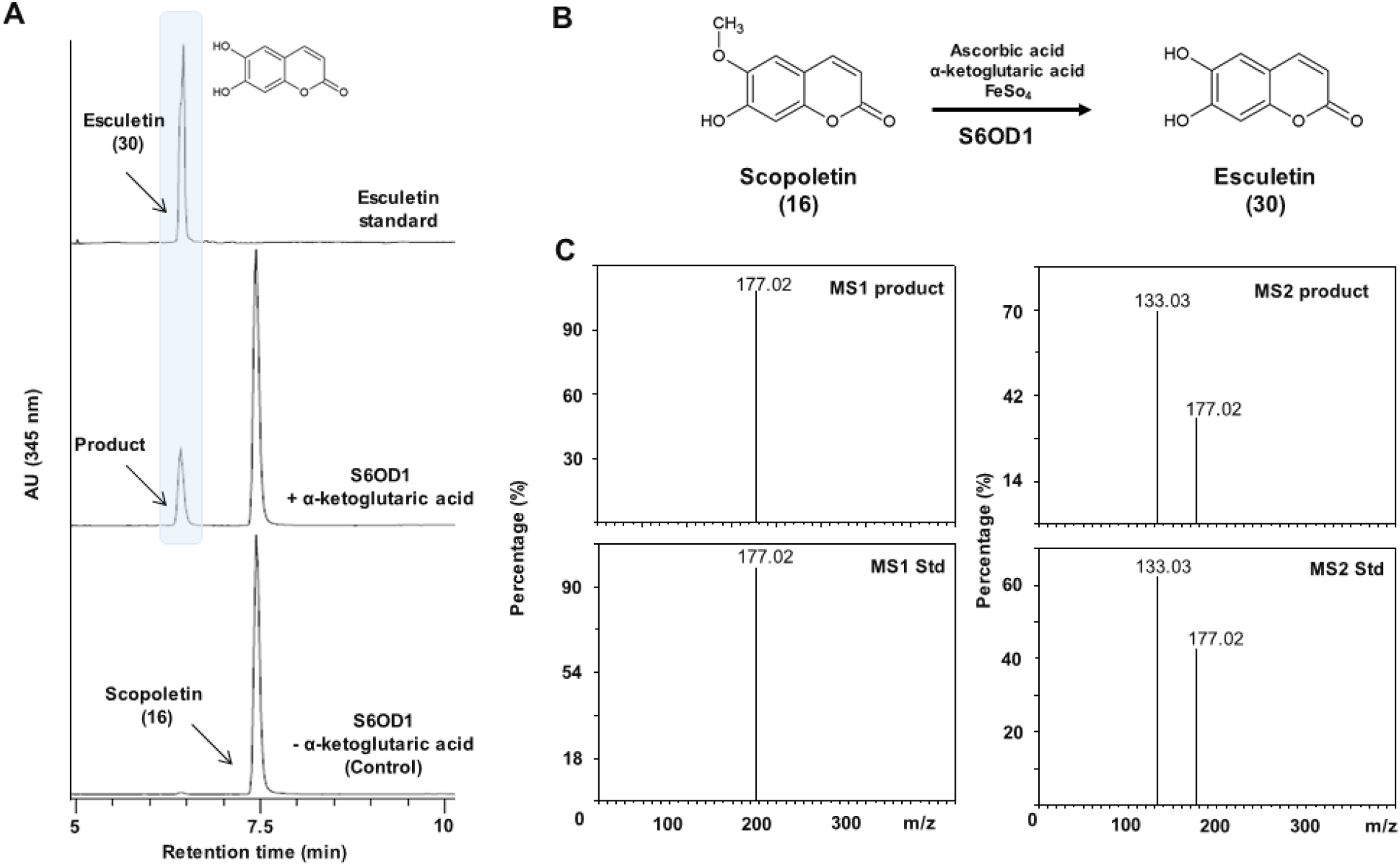
Biochemical characterization of the *O*-demethylase activity of S6OD1 towards scopoletin. (A) UV chromatograms at 345 nm of enzymatic reactions using renatured sonication pellets containing recombinant S6OD1. Reaction mixtures (100 µl) contained purified enzyme, 500 µM α-ketoglutarate, 500 µM ascorbic acid, 5 mM FeSO₄, and 200 µM *O*-methylated substrate, and were incubated in a mixed buffer system (acetic acid 0.1 M, Tris 0.1M, Imidazole 40 mM) under optimal temperature and pH conditions (Supplementary Figures X and Y) for 20 h. B) Schematic representation of the reaction converting scopoletin into esculetin. (C) Comparison of MS¹ and MS² spectra of the reaction product with those of authentic esculetin standard in negative ion mode using Orbitrap detection.

All AtODs except AtOD5 displayed *O*-demethylase activity toward at least one phenylpropane and one coumarin (**Supplementary Table S3**). Notably, only phenylpropanes bearing a carboxylic acid moiety were metabolized, whereas analogs containing aldehyde or alcohol groups remained unmodified, highlighting the function of the carboxyl group in substrate recognition. This is consistent with the essential role of Arginine305 in forming a hydrogen bond with the carboxylate of succinate and α-ketoglutarate (Kluza et al., 2018). In all active phenylpropanes, *O*-demethylation occurred at the para-methoxy group. However, none of these molecules have yet been described as endogenous metabolites in Arabidopsis. For coumarins, demethylation was observed at either the 6- or 7-position of the benzene ring, depending on the substrate, though no clear structure-activity relationship was evident regarding methoxy or hydroxy group positioning. All four active AtODs catalyzed the transformation of 3,4-dimethoxycinnamic acid and scoparone.

The most striking outcome of this study is the highly specific *O*-demethylation of scopoletin by AtOD1 and AtOD2 to yield esculetin, leading to their designation as S6OD1 and S6OD2, respectively (**Figure 3 and Supplementary Figures S7B, S8A, S10**). The apparent affinity parameters measured for both enzymes strongly support their physiological role *in planta* (**Figure 4 and Supplementary Table S3**). This hypothesis is further reinforced in case of S6OD1 by the predicted expression of the gene in root tissues (Klepikova et al., 2016), which our genotyping confirmed (**Supplementary Figure S26B**). Expression in this tissue supports a potential function in the metabolism of methoxylated coumarins, which are linked to Fe mobilization processes. S6OD1 and S6OD2 also converted 3,4-dimethoxycinnamic acid and scoparone into ferulic acid and isoscopoletin, respectively (**Supplementary Figures S7C, S8B, S11-S12, and Supplementary Figures S7D, S8C, S13-S14**, respectively). While ferulic acid is a central metabolite in Arabidopsis, the presence of its dimethoxylated precursor has not been reported *in planta*. Similarly, the lack of detectable isoscopoletin in roots questions the physiological relevance of this reaction. In addition, S6OD1 catalyzed the *O*-demethylation of herniarin to 6-hydroxycoumarin (**Supplementary Figures S7A, S15)** and showed trace activity on 5-hydroxy-7-methoxycoumarin (**Supplementary Figure S16) -** neither of which have been identified in Arabidopsis. Isofraxidin (Clemens and Weber, 2016) was also detected in trace amounts, but low signal intensity precluded kinetic characterization. Nevertheless, MS² analysis indicated that the enzymatic product was not fraxetin but likely isofraxetin (Clemens and Weber, 2016) (**Supplementary Figure S17**). Finally, S6OD2 converted 3,4,5-trimethoxycinnamic acid - a compound not reported in Arabidopsis - into sinapic acid (**Supplementary Figure S8D, S18**), a major endogenous metabolite. Despite favorable affinity values, the physiological relevance of this conversion remains uncertain, and the existence of an as-yet-unidentified native substrate structurally related to 3,4,5-trimethoxycinnamic acid is plausible. In contrast, although AtOD3 efficiently converted 3,4-dimethoxycinnamic acid (**Supplementary Figure S19**), its weak apparent affinity (**Supplementary Table S3**) and the absence of this compound in Arabidopsis tissues argue against a physiological role, suggesting that its native substrate remains to be identified. AtOD3 also catalyzed the conversion of scoparone (**Supplementary Figure S20**). AtOD4, in addition to acting on scoparone and 3,4-dimethoxycinnamic acid, catalyzed the demethylation of daphnetin dimethylether ether to produce daphnetin 8-methyl ether (**Supplementary Table S3 and Figures S21-S24**), a compound not detected in Arabidopsis, supporting a non-physiological transformation (leak activity). Similarly, AtOD4 showed activity with 3,4,5-trimethoxycinnamic acid (**Supplementary Figure S25**), despite the absence of this substrate in planta, and exhibited only limited activity toward scoparone (**Supplementary Table S3**). Although its apparent affinity for these substrates was relatively high (**Supplementary Table S3**), the restricted substrate scope and the lack of endogenous product detection suggest that, as for AtOD3, the physiological substrate of AtOD4 is yet to be elucidated.

**Figure 4.**
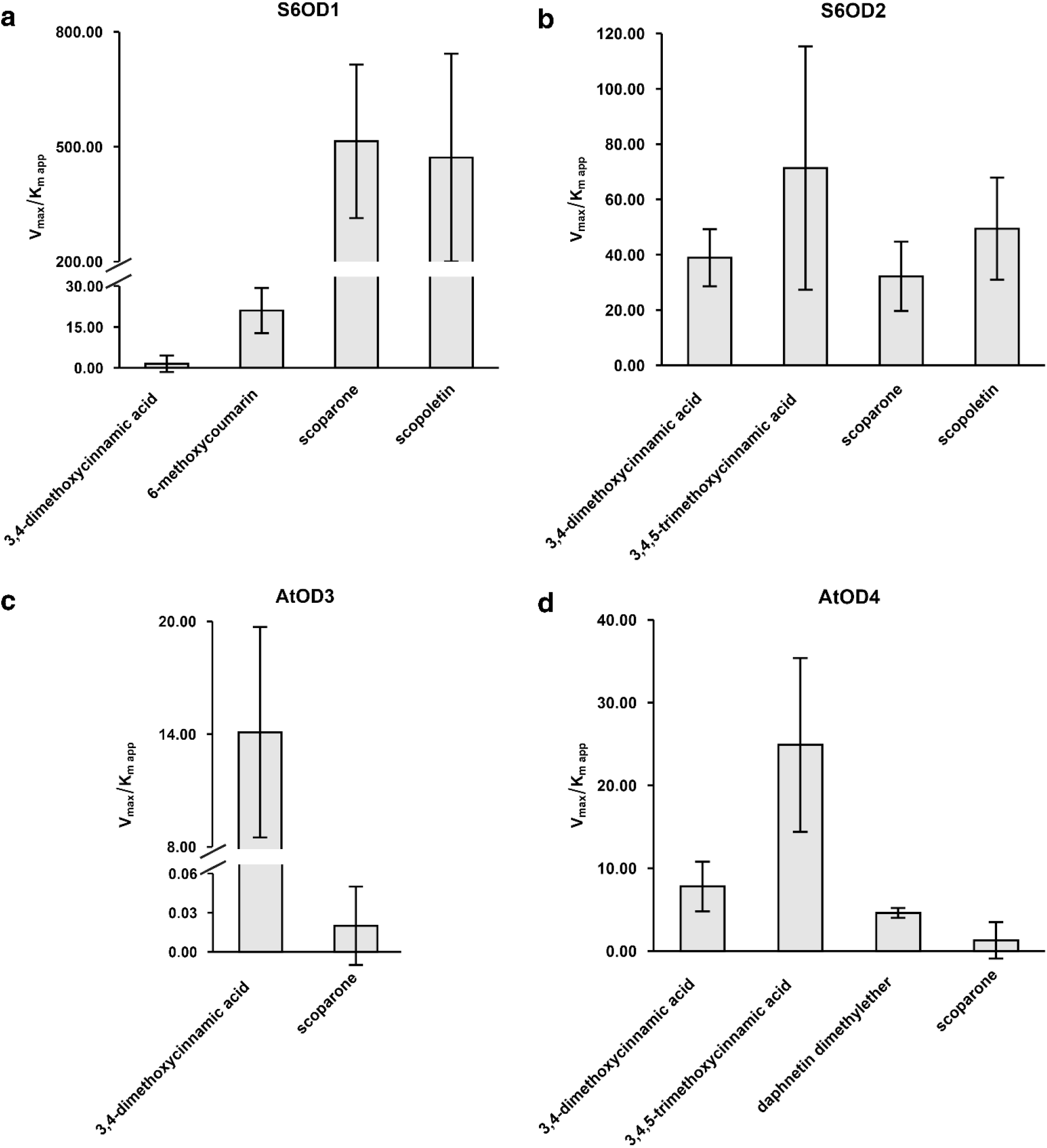
Comparison of catalytic efficiency of a S6OD1, b S6OD2, c AtOD3, and d AtOD4. Kinetic assays were performed under optimal temperature and pH conditions (Supplementary Figures X and Y). Reaction mixtures (100 µL) contained purified enzyme, 500 µM α-ketoglutarate, 500 µM ascorbic acid, 5 mM FeSO₄, and varying concentrations of *O*-methylated substrates, and were incubated in a mixed buffer system (acetic acid 0.1 M, Tris 0.1 M, Imidazole 40 mM). Products were detected at their maximal absorbance wavelength, and kinetic parameters were obtained by fitting the data with SigmaPlot 12.0 software. Catalytic efficiency was defined as Vmax/*Km app*, expressed in M⁻¹·s⁻¹. Each substrate concentration was tested in triplicate, and error bars represent standard error.

### S6OD1 catalyzes *O*-demethylation of scopoletin to esculetin in Arabidopsis roots

To verify whether the activity shown *in vitro* occurs *in planta*, for both *S6OD1* and *S6OD2* genes we identified two independent T-DNA insertion mutant lines, which were designated as *s6od1-1, s6od1-2* (**Supplementary Figure S26 and Tables S4-S5**), *s6od2-1* and s6od2-2 (**Supplementary Figure S27 and Tables S4-S5**). Generally, coumarin accumulation occurs at highest concentrations in roots, which is consistent with the Fe uptake function they have been shown to be crucial for (Schmid et al., 2014). Fe-responsive coumarins, such as scopoletin, esculetin, fraxetin and sideretin, accumulate differentially under Fe deficiency, but also in response to varying pH levels. Accumulation of esculetin has previously been shown to be promoted by both Fe deficiency and elevated pH (Paffrath et al., 2024). Based on this, we performed hydroponic experiments under Fe-deficient conditions at optimal (5.6) and increased (6.5) pH to identify genotype-dependent differences in root coumarin profiles.

Here, we cultivated Col-0, *s6od1* and *s6od2* mutants first in control Heeg hydroponic media (pH 5.6, 25 μM Fe^2+^-EDTA) and later transferred to pH 5.6 or 6.5 with 25 or 0 μM Fe^2+^-EDTA. The only line in which esculetin was consistently accumulated in quantifiable concentrations across conditions was WT Col-0 (**Figure 5**). In both *s6od1* mutant lines, esculetin was not detected in any of the conditions. In *s6od2-2* roots, depending on conditions, esculetin was not detected or present in trace amounts, whereas in line *s6od2-1* it was quantifiable, present in trace conditions or not detected. Levels of the glycosylated version of esculetin - esculin – were significantly lower (p-val < 0.05) in both *s6od1* lines than in Col-0 at pH 5.6 with no Fe present in the media. In the remaining conditions, the difference in esculin content between Col-0 and *s6od1-1* was significant, and in *s6od1-2* average esculin concentrations were lower than in Col-0, but differences remained not statistically significant (p-vals: 0.051 – 0.08). No statistically significant differences have been observed in esculin levels between Col-0 and *s6od2* lines, with the trend of higher values observed in the mutant lines.

**Figure 5.**
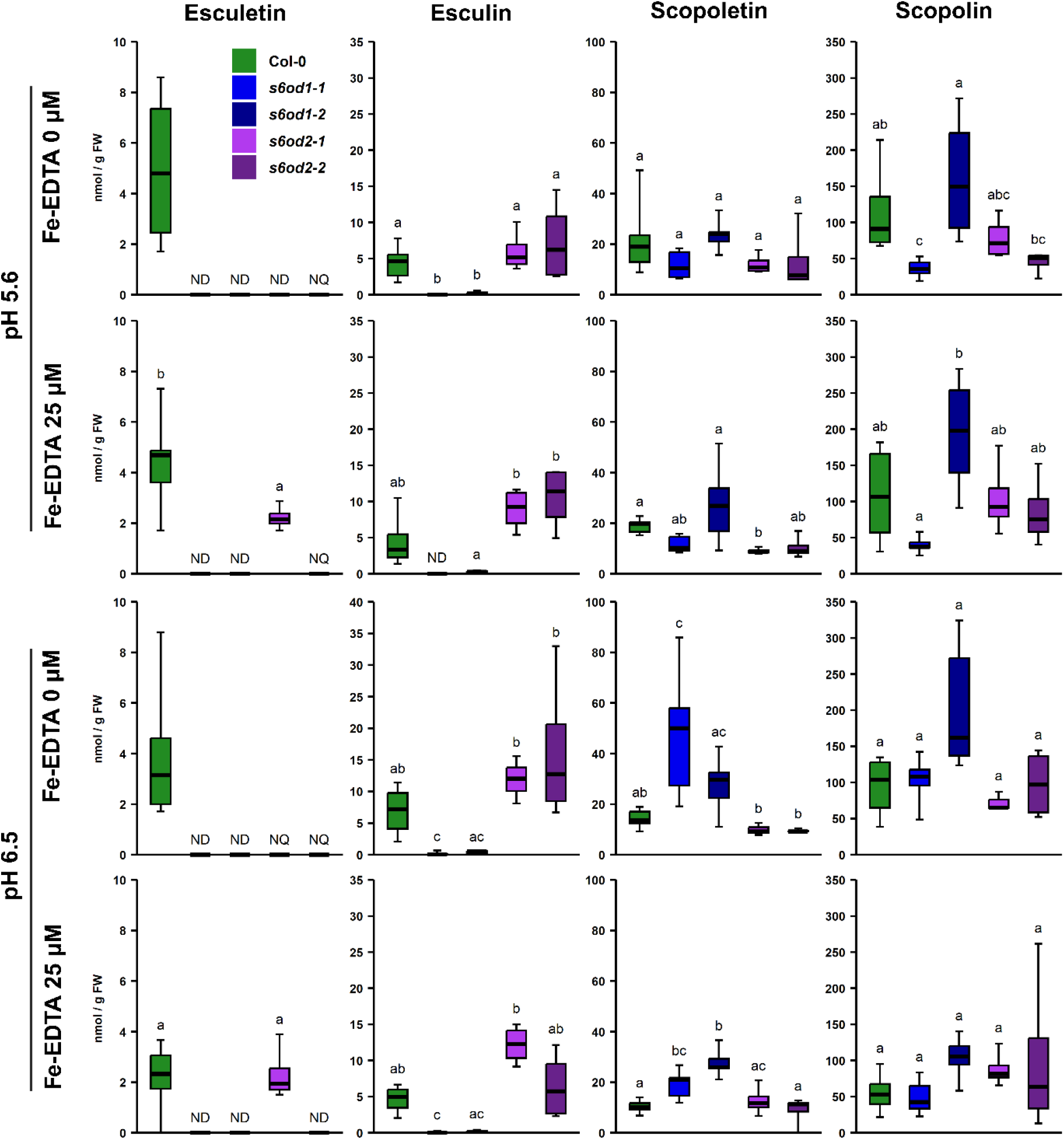
Esculetin, esculin, scopoletin and scopolin content in roots of Col-0, *s6od1* and *s6od2* mutants. Plants were grown for 10 days in control Heeg media (pH 5.7, 25 µM Fe^2+^-EDTA) before transferring to pH 5.7 or 6.5 with 25 or 0 µM Fe^2+^-EDTA, where they were cultivated for additional 5 days. ND – not detected, NQ – not quantifiable. n = 3 - 6. Statistical tests performed were ANOVA with Tukey’s post hoc or Kruskal-Wallis test with Dunn’s post hoc and Benjamini-Hochberg adjustment for multiple comparisons or two-tailed Student’s *t*-test, depending on data.

The concentrations of the reaction product, scopoletin, did not differ between Col-0 and *s6od1* lines at pH 5.6, however at pH 6.5 they were significantly higher for *s6od1-1* than Col-0 at both Fe conditions, whereas for *s6od1-2* the difference was significant only under Fe-depleted conditions. Scopolin concentrations did not differ between Col-0 and mutant lines (**Figure 5**). We observed increased scopoletin concentrations in *s6od1* mutant lines when compared to Col-0 under control conditions in both shoots and roots, but not exudates of plants grown in liquid half-strength Murashige-Skoog (MS) *in vitro* cultures with 3 % sucrose. The same was not observed when plants experienced Fe deficiency or mannitol-induced osmotic stress with the same 3 % sucrose-supplied media (**Supplementary Figures S28-S30**).

The results we obtained confirm the scopoletin *O*-demethylation activity of S6OD1 and S6OD2 *in planta*, showing no accumulation of the reaction product (esculetin) in *s6od1* lines and lower than in Col-0 in *s6od2* lines. As scopoletin concentrations in roots are approximately 10 times higher than those of esculetin, the lack of differences in scopoletin levels between Col-0 and knockout mutants at low pH is expected. We hypothesize that the increased concentrations of scopoletin in the mutant lines we observe at higher pH are not directly the effect of the disrupted scopoletin to esculetin conversion itself, but instead reflect the higher need for scopoletin to compensate for lack of esculetin’s activity.

It appears that in Arabidopsis roots, S6OD1 is the main enzyme responsible for biosynthesis of esculetin, with S6OD2 playing a supporting role, similarly to the relationship between F6’H1 and F6’H2 catalyzing the hydroxylation of feruloyl-CoA upstream in the coumarin biosynthetic pathway.

Interestingly, in our experimental conditions we did not observe statistically significant changes in esculetin accumulation based on higher pH or low Fe availability in Col-0, which was previously shown by Paffrath et al. (2024). This could be explained by either differences in experimental timepoints or the fact that the growth medium used (Heeg) has an overall lower concentration of nutrients than the very nutrient-rich Murashige-Skoog medium. Therefore, using the same molar Fe concentration, the ratio of Fe to other nutrients is higher in Heeg than in Murashige-Skoog media.

### Knockouts of *s6od1* and *s6od2* affect Arabidopsis performance

When root to shoot mass ratio of plants grown under the same conditions was compared, we noted differences between Col-0 and *s6od1* and *s6od2* mutant lines. Under all experimental conditions the mass ratio of roots to shoots of *s6od1-2* and *s6od2-2* mutant lines was significantly lower than that of Col-0. The *s6od1-1* mutant line sustained the trend, with the difference being significant at pH 5.6 with 25 μM Fe^2+^-EDTA (**Figure 6**).

**Figure 6.**
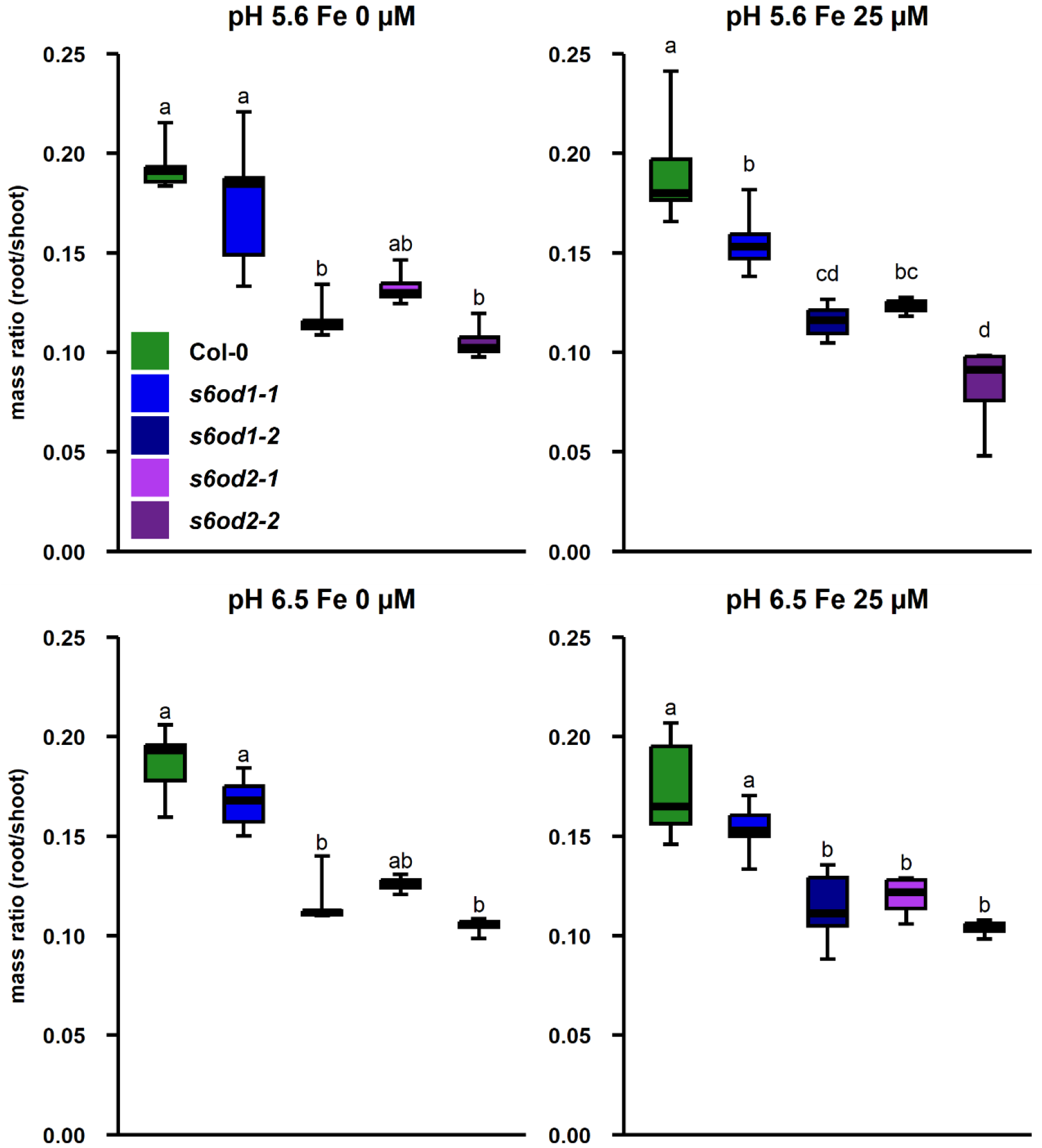
Root to shoot mass ratio of Col-0 and *s6od1* plants grown hydroponically at different pH and Fe availability conditions. Plants were grown for 10 days in control Heeg media (pH 5.7, 25 µM Fe^2+^-EDTA) before transferring to pH 5.7 or 6.5 with 25 or 0 µM Fe^2+^-EDTA where they were cultivated for additional 5 days. n = 4 – 6. Statistical test performed was ANOVA with Tukey’s post hoc.

The root to shoot mass ratios were fairly stable in the tested pH and Fe availability conditions, with no significant differences detected within line between conditions. Overall, we show that the knockout of either of the genes results in affected plant development. We propose that the effect points to a role of either esculetin or scopoletin in Arabidopsis development that is not related to Fe uptake function. This has been previously described when scopoletin-treated roots have shown cell and tissue abnormalities (Graña et al., 2017). Negative influence of both scopoletin and esculetin on root length has been shown for lettuce and arugula, but not barley (Facenda et al., 2023).

### Altered root coumarin localization profiles in flowering-stage *s6od1* and *s6od2* mutants

Spatial imaging of the glycosylated coumarins (scopolin, esculin, fraxin) performed by Robe et al., (2021) has shown that all three of them were present in root hairs of 7-day-old seedlings. We have failed to confirm expression of *S6OD2* using RNA extracted from rosette leaves, which is in agreement with available data. Literature suggests that under standard conditions expression of *S6OD2* does not occur in vegetative tissues (Jiang et al., 2021), but is limited to seeds (Klepikova et al., 2016). Root Cell Atlas (https://rootcellatlas.org/), where gene expression is mapped in five to seven-day-old primary root tips, indicates no expression of *S6OD2* in the roots, while reveals that *S6OD1* is principally expressed in the meristematic zone, particularly the root cap. This contrasts with the spatially distant expression of other genes known to be related to coumarin biosynthesis: *F6’H1, F6’H2* and *COSY* are expressed primarily in the differentiation and late elongation zone, and *S8H* as well as *CYP82C4* in differentiation zone (Shahan et al., 2022).

Here, we decided to investigate the general spatial coumarin accumulation profiles in roots of flowering-stage Col-0, *s6od1* and *s6od2* plants. For this, we first cultivated plants in standard peat-based soil conditions, with fertilizer added regularly. This was done to support physiological root development. At 3.5 weeks of growth, after removing the plant from soil and rinsing the roots, plants were transferred for two weeks to hydroponic conditions at pH 6.5, with 25 μM Fe^2+^-EDTA. At harvest, roots were cut longitudinally into three parts, equal in length, which we will here refer to as upper, middle and lower parts of the root.

Coumarin profiling has shown that the levels of esculin remained relatively consistent throughout the root parts in Col-0 (**Figure 7**), suggesting that either it is mobile and distributed throughout the root, or expression patterns of *S6OD1* change at later developmental stages. Esculin levels in *s6od1* mutants were non-detectable or were significantly lower than in Col-0 in all parts of the roots, except *s6od1*-*2* in the upper part of the root (here p-val = 0.08). Col-0 and *s6od2* lines have not shown statistically significant differences in esculin levels, although in the lower part of the root the concentration was on average lower by 66 % and 51 % in the *s6od2*-*1* and *s6od2*-*2* mutant lines than in Col-0, respectively (p-val = 0.06 and 0.15) (**Figure 7**), which could indicate that this is the part of the root where S6OD2 plays a supporting role in esculetin biosynthesis. Esculetin levels remained non-quantifiable or non-detectable due to low sample mass (**Supplementary Figure S31**).

**Figure 7.**
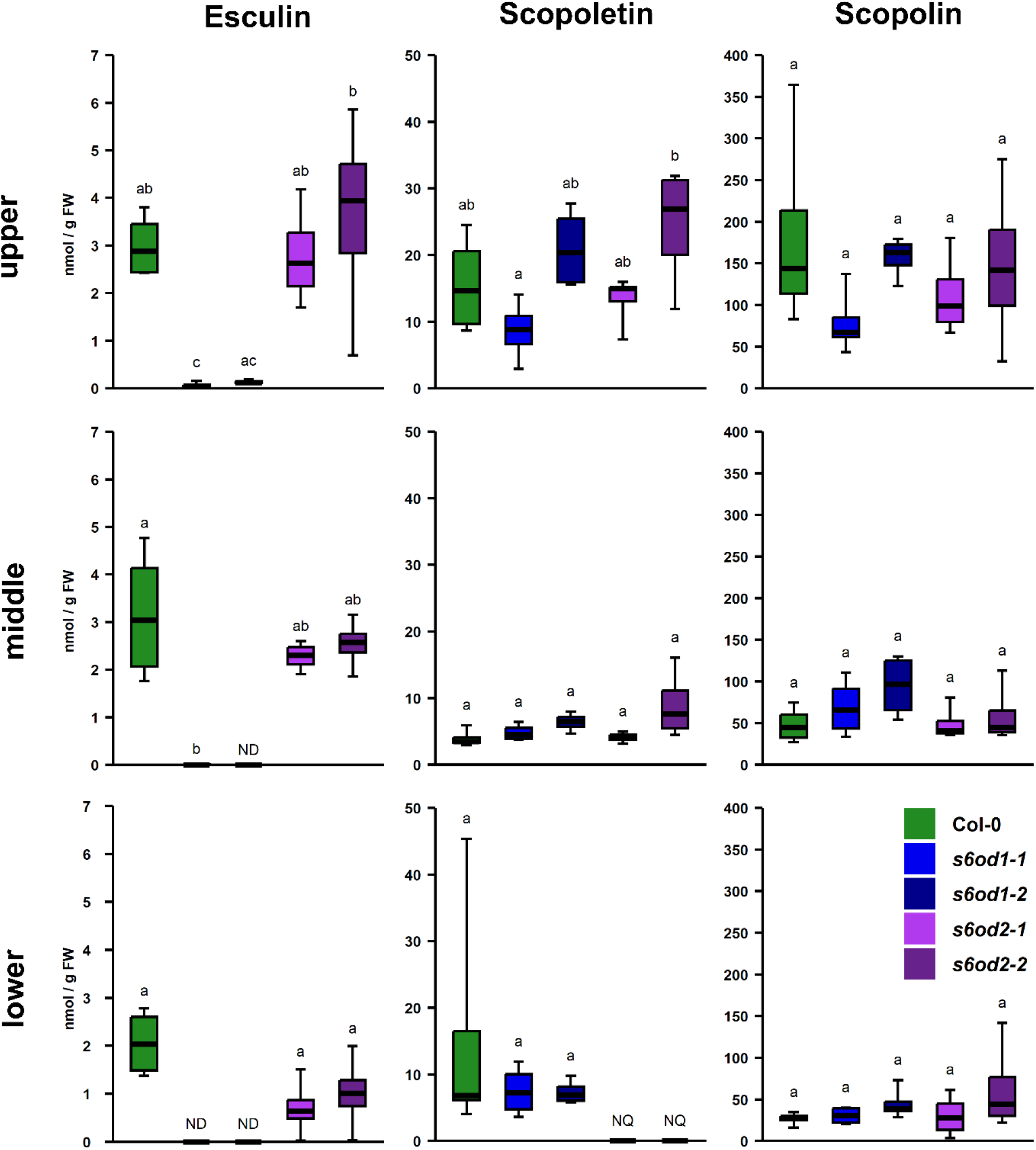
Esculin, scopoletin and scopolin content in parts of roots of Col-0, *s6od1* and *s6od2* mutants. Plants were grown first in soil with sufficient fertilization for 3.5 weeks and then hydroponically in Heeg medium (pH 6.5, 25 µM Fe^2+^-EDTA) for additional 2 weeks. Results are presented for upper, middle and lower part of the roots. ND – not detected, NQ – not quantifiable. n = 4 (each consisting of root parts of 3 independently grown plants). Statistical tests performed were ANOVA with Tukey’s post hoc or Kruskal-Wallis test with Dunn’s post hoc and Benjamini-Hochberg adjustment for multiple comparisons, depending on data.

In Col-0, the concentration of scopolin was higher in the upper part of the root than the lower. Similar trend was observed in the mutant lines’ roots (p-val = 0.13 (*s6od1-1*), 0.0009 (*s6od1-2*), 0.02 (*s6od2-1*), 0.6 (*s6od2-2*)). Scopoletin levels in Col-0 roots were the highest in the upper part of the root (p-val upper:middle = 0.02, upper:lower = 0.06). The same trend was observed in the mutant lines, although the differences were not statistically significant in *s6od1-2* line. Interestingly, scopoletin levels in both *s6od2* lines were non-quantifiable in the lower part of the root. No differences have been detected in scopoletin or scopolin levels between Col-0 and *s6od1* lines, which distinguishes these results from the ones obtained by analyzing coumarin content of plants grown hydroponically from seed (**Figure 5**). This could be explained by the difference in the duration of plant growth as well as by the initial growth in soil, which not only differed in nutritional availability, but also influence of the soil microbiome on the plant root. This plant-microbiome relationship has been shown to be important to coumarin metabolism (Stassen et al., 2021). It was recently shown that a fungal endophyte, *Macrophomina phaseolina* exhibits capacity to modify scopoletin into esculetin and enhance Fe nutrition (Van Dijck et al., 2025).

### S6OD1 affects plant performance at limited Fe availability under alkaline conditions

In experiments performed at acidic conditions, we have not observed any Fe deficiency-related phenotype distinctly associated with *s6od1* mutants. Considering the capacity of esculetin to mobilize and chelate Fe, especially its unique high efficiency at alkaline pH (Schmid et al., 2013, Paffrath et al., 2024), we decided to test how the substitution of the Fe source from chelated Fe^2+^ EDTA to unchelated Fe^3+^ in the form of FeCl_3_ at alkaline pH will affect plant growth in case of Col 0 and *s6od1* mutant lines.

When plants were cultivated on solid quarter-strength Murashige-Skoog medium supplemented with 20 µM FeCl_3_ at pH 7.4, coumarin profiling of Col-0 and *s6od1* mutants’ roots has shown lower levels of esculetin in the mutant lines, consistent with metabolic profiling performed for plants grown in chelated Fe. Here there was no increase in scopoletin or scopolin concentrations in *s6od1* lines (**Supplementary Figure S32**). We observed uneven phenotypes between plates and experimental replicates, which we speculate was due to low solubility of FeCl_3_, leading to precipitation and varying concentration of Fe on plates. Therefore, we decided to continue with experiments performed in liquid media.

Fraxetin-deficient mutant *s8h* was used as a marker line for Fe deficiency. Plants were first grown for 10 days in 2.5 µM Fe-EDTA (pH 7.4) and then cultivated for three weeks in either 2.5 µM Fe-EDTA or 2.5 µM FeCl_3_-supplemented media. Both conditions supply a sub-optimal level of Fe, however the unchelated Fe source results in lower availability and the demand for coumarin-facilitated Fe uptake.

In case of Col-0 grown with unchelated Fe, while loss in rosette mass was observed, there were very limited signs of chlorosis on some of the plants and the chlorophyll content was not significantly decreased. It has been previously shown that fraxetin plays a major role in the process of Fe-mobilization (Siwinska et al., 2018, Rajniak et al., 2018), which our results confirmed by the stark difference in phenotype, mass and chlorophyll content of *s8h* between the conditions. Similarly to Col-0 and *s8h*, both *s6od1* lines had decreased rosette mass when grown with unchelated Fe. Contrary to Col-0, *s6od1* lines exhibited a significant decrease in the chlorophyll content in the unchelated Fe condition. Some of the plants developed a clear chlorotic phenotype (**Figure 8**).

**Figure 8.**
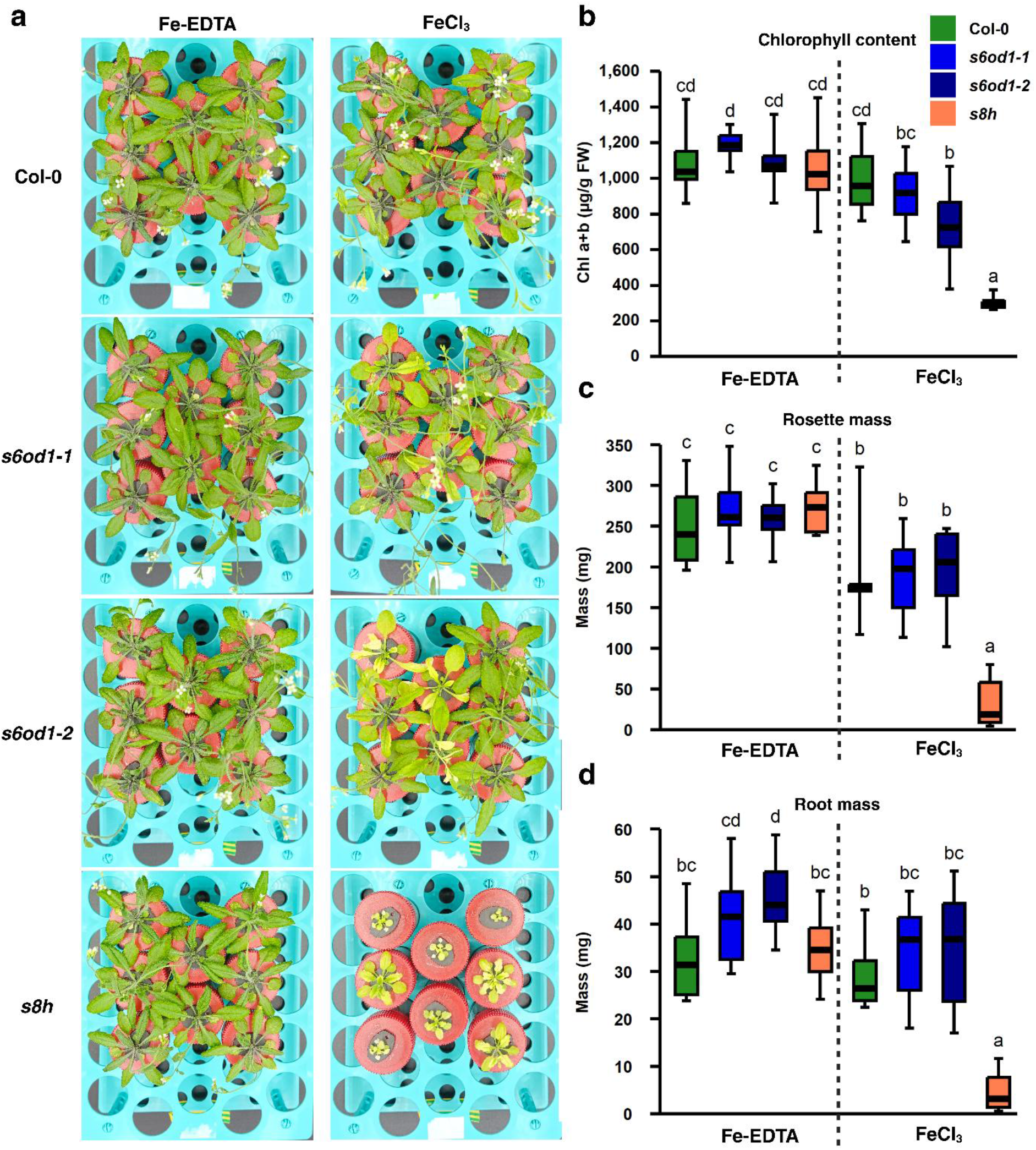
Growth of Col-0, *s6od1* and *s8h* knockout mutant plants in alkaline media with varying Fe availability. Plants were grown for 10 days in pH 7.4 Heeg media supplemented with 2.5 µM Fe^2+^-EDTA, then in media containing 2.5 µM Fe supplied either as Fe^2+^-EDTA or unchelated FeCl_3_ for additional three weeks. **a** plant phenotype 2.5 weeks after transfer, n = 8 **b** chlorophyll a + b concentration in rosettes at harvest, n = 5 for *s8h* cultivated in FeCl_3_ condition, 8 for all other groups **c** rosette mass at harvest, n = 8 **d** root mass at harvest, n = 8. Statistical test performed was Two-Way ANOVA with Benjamini-Hochberg post-hoc.

Our results suggest that esculetin affects Fe deficiency tolerance at limited Fe availability conditions at alkaline pH, with fraxetin playing a much more crucial role. We speculate that although it possesses high Fe-mobilizing activity, esculetin’s main role in Arabidopsis physiology does not lay in Fe deficiency response, although it seems to provide more tolerance under specific conditions of limited availability. Recent research suggests that coumarins have a significant function in plant-microbiome interactions (Stringlis et al., 2018; Gu et al., 2026). Due to its antioxidant properties esculetin could also play a role in response to a wide range of environmental stresses.

### Transient expression of AtS6OD1 in *Nicotiana benthamiana*

To test whether At*S6OD1* could catalyze the reaction of scopoletin *O*-demethylation to esculetin when expressed in a heterologous plant system, we transiently expressed the *S6OD1* alleles in *Nicotiana benthamiana* leaves. Here we utilized Agrobacterium-mediated transformation to express the *S6OD1* allele from Col-0, producing a functional copy of the protein, and one derived from another natural accession, Stobowa (Stw-0), which is characterized by an early translation termination allele of the *S6OD1* gene (**Supplementary Figure S33**). The experiment was carried out without or with additional scopoletin feeding to increase the concentration of the reaction substrate. We did not observe a clear AtS6OD1-dependent increase in esculetin or esculin accumulation (**Supplementary Figure S34**). This likely reflects the strong background of the concurrent endogenous esculetin metabolism in *N. benthamiana*, which was itself markedly affected by Agrobacterium infiltration, making it difficult to resolve enzyme-specific effects under these conditions. Interestingly, when scopoletin feeding was performed, the levels of esculin and scopolin were increased in all Agrobacterium-infiltrated groups compared to mock infiltration media. This highlights the role of these compounds in response to biotic factors.

## DISCUSSION

In this work, we present the first case of *O*-demethylation reaction occurrence in the model plant *Arabidopsis thaliana*. Although many instances of *O*-methylation have been showcased within the plant kingdom both within primary and secondary metabolism, the *O*-demethylation reaction has previously been only shown in two plant species, 1) *Papaver somniferum* (opium poppy) in the highly specialized isoquinoline alkaloid pathway, leading to the biosynthesis of morphine (Farrow and Facchini, 2013), and 2) *Ocimum basilicum* (basil), in the lipophilic flavone network (Berim et al., 2013). It is our expectation that *O*-demethylation will prove to be more widespread in the plant kingdom than previously assumed, occurring in various species and metabolic pathways. This reaction may either produce final products (as in the opium poppy for morphine biosynthesis) or, as we demonstrate in our case with Arabidopsis, participate in the reverse reaction of *O*-methylation. This reveals a new dynamic dimension to metabolic pathways traditionally considered unidirectional. The possibility of *O*-demethylation should therefore be considered when the biosynthesis of a plant-produced compound remains undescribed. In this way, we hope that our findings will pave way to further discoveries of *O*-demethylation within plant biosynthetic pathways, both in Arabidopsis and other plant species.

Notably, plant *O*-demethylases could serve as valuable tools in metabolic pathway engineering, particularly if they exhibit substrate promiscuity or on the contrary substrate specificity and regioselectivity of reaction. These enzymes could enable the production of rare hydroxylated compounds from more abundant O-methylated precursors, which are typically more stable and less reactive. This approach would leverage the natural metabolic versatility of plants for bioengineering applications. As we originally identified the Arabidopsis *O*-demethylases through their sequence similarity to the opium poppy *O*-demethylases, the same approach could lead to discoveries of *O*-demethylation in other plant species. With the rapid development of protein structure prediction tools, such as AlphaFold, the search can be widened by comparisons of protein structure on the 3D level.

From an applied perspective, the identification of new *O*-demethylases in a model plant expands the metabolic engineering toolbox by enabling flux redirection toward valuable hydroxylated metabolites. As esculetin and esculin are currently obtained mainly from *Fraxinus* spp. or *Aesculus hippocastanum*, identification of their biosynthetic genes could facilitate sustainable production. Although engineering of downstream glycosylation can enhance esculin yields (Zhang et al. 2026), further pathway optimization requires better control of upstream steps leading to esculetin formation. Our identification of a key enzymatic step therefore provides a molecular framework for engineering esculetin-derived compounds.

We considered three possible routes leading to biosynthesis of esculetin: 1) hydroxylation of caffeoyl-CoA to 6-hydroxycaffeoyl CoA with subsequent isomerization and lactonization to esculetin, as shown previously *in vitro* (Vanholme et al., 2019); 2) hydroxylation of umbelliferone and 3) the *O*-demethylation of scopoletin, which we are showing in this work. This third possibility was previously suggested based on metabolite profiling of *f6’h1* mutants (Schmid et al., 2014), but remained experimentally unconfirmed. Non-enzymatic formation in soil has been reported, for example *via* oxidation of caffeic acid (Deiana et al., 1995). Recently, the fungal endophyte *Macrophomina phaseolina* has been shown to convert scopoletin to esculetin (Van Dijck et al., 2025). The physiological relevance of such alternative routes for plants remains to be determined and is likely to depend on environmental conditions, particularly in the rhizosphere. Our hydroponic experiments with different Fe sources further indicate that S6OD1-dependent esculetin formation contributes to Fe-responsive physiology *in planta*, as *s6od1* plants showed genotype-specific changes in chlorophyll content depending on Fe form, whereas the wild type remained largely stable, consistent with partial redundancy of Fe acquisition systems. It remains to be explored whether the same reaction, *O*-demethylation of scopoletin, leads to biosynthesis of esculetin in other plant species. An approach worth consideration would be to investigate homologs of *S6OD1* and *S6OD2* genes, as well as structurally similar proteins, particularly in agronomically important Fabids and Malvids, where AtOD orthologs are present. This is especially relevant for crops such as cassava (*Manihot esculenta*), where scopoletin and esculetin accumulation is linked to postharvest tuber deterioration (Buschmann et al., 2000). Our phylogenetic analyses, together with recent functional reports from *Fraxinus* (Kong et al. 2026), indicate that coumarin *O*-demethylation is not restricted to Arabidopsis but likely distributed across diverse plant lineages, suggesting repeated recruitment of this chemistry during plant evolution. Such cases of convergent evolution have been previously reported in other plant metabolic pathways (Colinas et al., 2025; Huang et al., 2016).

Relatively high concentrations of esculin in shoots were observed here and in previous work (Perkowska et al., 2021). Given that *S6OD1* and *S6OD2* are not expressed, or expressed at low levels in shoots, we propose that esculin in aerial tissues is either transported from the roots, formed nonenzymatically from 6-hydroxycaffeoyl-CoA under light, or produced via COSY-catalyzed conversion (Vanholme et al., 2019). The trans–cis isomerization followed by a lactonization activity of COSY does not however appear to be of significance in roots, as it is not able to complement *s6od1* knockout. Whether it plays a role in esculetin biosynthesis in shoots remains to be confirmed, as this activity has so far only been shown *in vitro*, and the reaction kinetics remain unexplored. It is unclear whether the reaction occurs *in planta* and whether the required substrate concentration is physiological. The substantial shift in scopoletin accumulation observed in *s6od1* suggests that this step affects broader flux distribution within the coumarin network rather than acting as a simple linear conversion, indicating pathway-level homeostatic regulation.

It has been shown previously as well as in this work that esculetin and esculin, among other coumarins, are accumulated in response to both abiotic and biotic stresses in various plant species. This includes Fe deficiency (Paffrath et al, 2024) drought stress (Fini et al., 2012) and pathogens (Witzel et al, 2024., Stringlis et al., 2019). Taking into consideration that coumarins are able to modulate rhizosphere microbiome by selectively inhibiting growth of various bacterial and fungal species (Stringlis et al., 2018; Voges et al, 2019) it stands to reason that further research is needed to understand the mechanisms at play. In this context, S6OD1-dependent remodeling of coumarin composition may influence root–microbe interactions, as coumarins including scopoletin are involved in both beneficial fungal associations and microbial competition in the rhizosphere, linking this metabolic branch to ecological outcomes beyond Fe homeostasis (Van Dijck et al. 2025).

As esculetin accumulates at concentrations much lower than other Fe-responsive coumarins, the chief difficulty in working with this compound lies in obtaining concentrations exceeding the limit of quantification. The highly reactive catecholic motif may contribute to this challenge by limiting its stability and detectability. Low accumulation levels of esculetin, compared to those of other Fe-responsive coumarins, together with the atypical spatial expression profile (expression of *S6OD1* in the meristematic part of the root and expression of *S6OD2* is dry seeds) might suggest that esculetin plays a regulatory or protective, possibly antioxidative, role in the selected tissues or cell types in which expression of the responsible genes is higher. Expression of *S6OD2* in seeds might suggest a role of esculetin in seed maturation or protection of seeds against stressors such as plant pathogens. It has been shown that exogenously applied coumarins inhibit seed germination of different plant species (Chen et al., 2019; Zhang et al., 2023). Very little is known however about coumarin profile of Arabidopsis seeds. Experiments performed with *s6od1* and *s6od2* mutants clearly demonstrate the essential role of S6OD1 and S6OD2 in esculetin biosynthesis in roots. At the same time, under the experimental conditions we tested, esculetin is non-essential, as scopoletin and the catecholic coumarin fraxetin appear to compensate for its functional role in Fe deficiency response.

Based on our *in vitro* substrate metabolization results and catalytic parameter determinations, none of the tested compounds appear to be physiological substrates for AtOD3 and AtOD4, and the observed substrate affinity (>100%M) falls outside the typical range for DOXC enzymes. The function of AtOD5 remains unknown. Although all AtODs belong to the DOXC class of 2OGDs, their placement in the DOXC52, distinct from coumarin biosynthesis related DOXC30 enzymes, could suggest different substrate specificity. As seen in other DOXC clades (e.g. DOXC47, with flavonol vs anthocyanidin synthases; Kawai et al., 2014), closely related enzymes can act on different metabolite classes despite their structural similarity. We can hypothesize that AtOD substrates may belong to different chemical classes, while sharing a benzene ring backbone (as demonstrated for alkaloid and flavonoid *O*-demethylation). These substrates could therefore participate in distinct metabolic pathways. Purified AtOD3 protein was shown to be able to catalyze hydroxylation of melatonin to 3-hydroxymelatonin *in vitro* with K_m_ value of 100 µM (Lee and Back, 2022). Based on publicly available expression data, expression of AtOD3 appears to be most strongly, out of the 5 AtOD genes, responsive to pathogens (Arabidopsis eFP Browser data; Whitham et al., 2003; Ascencio-Ibáñez et al., 2008), which supports its role in secondary metabolism.

The physiological analyses of plants grown in hydroponic cultures with different Fe sources suggest that the identified activity contributes to Fe-dependent responses *in planta*. Genotype-specific changes in chlorophyll content under different Fe sources support a link between S6OD1-dependent coumarin metabolism and plant responses to Fe availability. Given the complexity and partial redundancy of Fe uptake mechanisms in Arabidopsis, the subtle nature of the phenotype is not surprising. S6OD1 activity may also influence other processes mediated by coumarins, as esculetin and esculin have been implicated in biotic and abiotic stress responses, as well as in interactions with the root-associated microbiota. The altered scopoletin accumulation observed in *s6od1* mutants exceeding expectations based on a simple substrate–product relationship, suggests that S6OD1 contributes to coumarin homeostasis. The evolutionary context further highlights the potential significance of this pathway. Together with recent evidence from *Fraxinus*, phylogenetic analyses indicate that the responsible enzyme family is broadly distributed across plant lineages, suggesting that this activity is not restricted to Arabidopsis and may represent a more general feature of plant specialized metabolism.

Taken together, our results identify a previously unrecognized branch of coumarin metabolism with measurable physiological consequences. The combined biochemical, genetic, physiological and phylogenetic evidence supports a role for this pathway in shaping coumarin homeostasis and plant responses to environmental conditions. Our findings illustrate that important components of specialized metabolism remain to be discovered even in one of the most extensively studied model plant species and suggest that *O*-demethylation represents a more widespread, yet still underexplored, component of plant specialized metabolism across the plant kingdom.

## METHODS

### Plant material

*Arabidopsis thaliana* (Arabidopsis) Col-0 accession was used as the wild-type. The mutant lines: *s6od1-1* (SAIL_606_F09) and *s6od1-2* (WiscDsLox353A01), as well as *s6od2-1* (SALK_056936) and *s6od2-2* (SAIL_641_B03) and *s8h-2* (SM_3.23443) T-DNA insertional mutant lines were obtained from the Nottingham Arabidopsis Stock Centre (http://arabidopsis.info/). *Nicotiana benthamiana* seeds were kindly gifted by Dr Etienne Herbach (INRAE, Colmar, France).

### Growth conditions

#### Hydroponic cultures

Media used in hydroponic cultures was 1 x Heeg (Siwinska et al., 2018) buffered with MES hydrate (Sigma-Aldrich, 2.5 mM) for pH 5.6 or 6.5 and with MOPS (EMD Milipore, 1.25 mM) for pH 7.4. Seeds were placed on microcentrifuge tube lids filled with 0.65 % Heeg-based agar, which were then placed in liquid medium-filled plates. Seeds were stratified by placing the plates in the dark at 4 °C for 3 to 4 days and then transferred to controlled conditions growth chambers.

#### Hydroponic culture on plates

Plants were cultivated for 10 days in control Heeg media (pH 5.7, 25 µM Fe2+-EDTA) before transferring onto standard 90 mm diameter plates with dividers with 15 ml of either pH 5.7 or 6.5 with 25 or 0 µM Fe^2+-^EDTA. Four plants were placed in each half-plate. Plants were cultivated for additional 5 days, then harvested and flash frozen before processing, with 4 plants growing in each half-plate combined into one sample for metabolomic analysis.

Culture in 50 ml tubes. Plants were cultivated for 10 days in pH 7.5, 2.5 µM Fe^2+^-EDTA Heeg before transferring into opaque 50 ml falcon-type tubes, one plant per tube. Media used were pH 7.5, 2.5 µM Fe^2+^-EDTA and pH 7.5, 2.5 µM FeCl3. Plants were then cultivated for additional three weeks with media replenished (added without discarding) according to needs and then harvested and frozen. Rosettes were later used for chlorophyll content analysis.

#### Soil-to-hydroponics transferred plants

To stratify the seeds, they were placed on damp Whatman paper and kept in the dark at 4 °C for 3 to 4 days. Seeds were then sown into pots filled with peat-based commercial soil (COMPO SANA Substrate for sowing and transplanting) mixed 3:1 v:v with 2-6 mm vermiculite. Plants were watered according to their needs, general use fertilizer (Substral) was added once a week as per manufacturer’s recommendations. After 3.5 weeks of growth, plants were gently removed from soil and roots were rinsed with deionized water. Plants were then placed in opaque falcon-type 50 ml tubes filled with pH 6.5, 25 µM Fe^2+^-EDTA liquid Heeg medium, where they were grown for additional 2.5 weeks with media being replenished (added without discarding) every 2-3 days. Plants were then harvested with roots cut crosswise into 3 equal lengths (constituting upper, middle and lower section of the root). Root sections derived from 3 plants were pooled together as one sample for metabolomic analysis. Four samples were used. Samples were flash frozen before processing.

### Phylogenetic analysis

Protein primary structures for phylogenetic analysis were collected from Phytozome v13 (https://phytozome-next.jgi.doe.gov/) **(Supplementary File 1)** and Plant Garden (https://plantgarden.jp/ja/index) for *Papaver somniferum* using DOXC52 primary structures as queries. As for *Papaver somniferum*, *Ocimum basilicum*, and *Coptis japonica*, reported functional primary structures were also added in the list. Overlapped primary structures and too short sequences were removed (**Supplementary Figures S35**). The remaining primary structures were subjected to construct a multiple alignment in MAFFT ver. 7 (default setting) (https://mafft.cbrc.jp/alignment/server/index.html) (Kuraku et al., 2013; Katoh et al., 2019) and used to construct a phylogenetic tree by maximum likelihood method with 1,000 bootstrap tests and 1,000 Shimodaira-Hasegawa approximate likelihood-ratio test in IQtree (https://iqtree.github.io/) (default setting). The tree was visualized by Figtree v.1.4.4 (https://github.com/rambaut/figtree/releases/tag/v1.4.4). Primary structures were aligned with Bioedit Software.

### Plant RNA extraction for cloning and coding sequence synthesis

Total RNA was extracted from seeds, seedlings, or roots of Arabidopsis Col-0 accession using the E.Z.N.A.® SP Plant RNA Kit (Omega Bio-tek) according to the manufacturer’s instructions. cDNA synthesis of *At1g17020* (*AtOD1*), *At4g25310* (*S6OD1*), and *At1g78550* (*AtOD3*) from total RNA extracts was performed using the High-Capacity RNA-to-cDNA™ Kit (Applied Biosystems), following the manufacturer’s protocol. Coding sequences of *At1g17010* (*S6OD2*) and *At4g25300* (*AtOD2*) were obtained by de novo synthesis in the pUC57 vector (GenScript).

### Construction of Gateway™ recombinant plasmid pCold I and subcloning of coding sequences

The pCold™ I vector (Takara) was modified by insertion of a Gateway™ destination cassette. Briefly, the Gateway™ fragment was PCR-amplified using PrimeSTAR Max DNA Polymerase (Takara) and subsequently cloned by restriction–ligation between the *NdeI* and *XhoI* restriction sites. PCR amplification (**Supplementary Table S4**) of all coding sequences (from cDNA or pUC57 templates) was performed using PrimeSTAR Max DNA Polymerase. PCR products were subcloned into the pCR™8/GW/TOPO™ vector using the TA Cloning Kit (Invitrogen™) according to the manufacturer’s instructions. Recombination into the pCold I Gateway™ expression vector was carried out using Gateway™ LR Clonase™ II Enzyme Mix (Invitrogen™), following the manufacturer’s protocol. Recombinant plasmids were used to transform *E. coli* Rosetta 2 strain for heterologous expression of 6×His-tagged proteins.

### Heterologous expression and purification of DOXC52

*E. coli* Rosetta 2 strain transformed with pCOLD:*S6OD1* was cultured at 37 °C overnight in 10 mL LB supplemented with 100 mg/L ampicillin and 33 mg/L chloramphenicol. A 2 ml pre-culture was transferred to 1 L of fresh LB containing ampicillin 100 mg/L and chloramphenicol 33 mg/L. Transformed cells were cultured at 37 °C until OD_600nm_ reached 0.6. Overexpression was induced by an addition of 1 mM of IPTG and the culture was transferred to 15 °C. To purify the overexpressed protein a protocol of denaturation and renaturation was adapted from Olry et al., (2001). The supernatant was concentrated using an Amicon® Ultra Filter (cut off 10 kDa) and centrifuged for 10 min (15 000 g). The filtrate was discarded and the residual solution remaining in the column was transferred into an Eppendorf tube. At multiple points in the purification protocol an aliquot was taken for SDS-PAGE analysis to confirm protein solubility (**Supplementary Figure S36**).

### Enzymatic activities measurements

Previous to enzymatic studies of renaturated 2OGDs, the enzymatic preparation was followed to be stable over time up to 2 h with scopoletin as a substrate. For standard scopoletin *O*-demethylation activity measurement, cofactors were set as follows: Vitamin C 5 mM, α-ketoglutarate 5 mM and FeSO_4_ 500 µM. Optimal temperature and pH determination were determined with saturated concentration of 3,4 dimethoxycinnamic acid (400 µM each) and cofactors and in the range of 12 to 37 °C and 6.0 to 9.0, respectively. For substrate screening, substrate concentration was set at 200 µM. For kinetic parameters determinations, coumarin/phenolic acid apparent affinity were determined at saturating concentration of cofactors. Substrate concentration varied from 0.1 *k*_m_ to 10 *k*_m_. All the reactions were followed during 1 h at optimal pH and temperature. All the reactions were performed in buffer 100 mM Tris-HCl in a volume of 100 µl.

### Circular Dichroïsm

Circular dichroism (CD) spectra of the enzymes (5 μM in phosphate buffer 10 mM at pH 7.1) were obtained using using a X spectrometer. The spectra were scanned at 25°C with 1-nm steps from 260 to 190 nm and averaged over six scans.

### Methanolic extract preparation

Extracts were prepared by homogenizing plant tissue samples, sonicating for 10 mins and incubating overnight at 4 °C in 500 µL of 80 % methanol supplemented with 5 µM 4-methylumbelliferone as internal standard. Sample was then centrifuged at 18 000 g and supernatant transferred to a new tube. Additional 450 µL of 80 % methanol was then added to the sample, followed by sonication and centrifugation. Supernatant was added to the previously transferred batch. Samples were then dried in a vacuum centrifuge and later used for HPLC-MS analysis.

### Extraction of root exudates from nutrient solutions

Nutrient solutions from Arabidopsis hydroponic and liquid *in vitro* cultures were processed as described in Siwinska et al., 2018. Briefly, phenolic compounds were retained in a BAKERBOND™ C18 column (J. T. Baker Chemical Co., Phillipsburg, NJ, USA), eluted from the cartridge with 3 ml of 100 % methanol, and dried in a centrifugal evaporator. Dry extracts were stored at –20 °C until further analysis.

### UHPLC and LTQ LC/MS analysis

For UPLC analysis of the products, a NEXERA UPLC system (Shimadzu, Japan) equipped with a photodiode array (PDA) and a detector (SPDM20A, Shimadzu) was used. The chromatographic column was a C18 Core-Shell reverse phase (Kinetex XB-C18, Phenomenex, USA) 150 × 2.1 mm 2.6 µm. The products were quantified after separation with a gradient of Water + 0,1% Formic Acid (A) and Methanol + 0.1 % Formic Acid (B). The elution gradient was (A:B; v/v): 90:10 at 0 min, 80:20 at 0.74 min, 40:60 at 5.88 min, 10:90 at 10 min, 0:100 between 12 and 16 min, and 90:10 from 16.01 to 20 min. The total analysis lasted 20 min and a PDA range of 200–400 nm and the products were detected at their maximum wavelength. MS/MS analyses were performed on a Thermo Vanquish UHPLC Chain equipped with a Thermo Orbitrap XL detector (Thermo Electron, USA). We used a Phenomenex Kinetex XB-C18 150x2,1mm, 2,6µm (Phenomenex, USA) column to separate compounds with a 30 mn method using Water + 0,1% Formic acid (A) and Methanol +0,1% Formic Acid (B) following the gradient (from 0 to 1mn 10% (B), increasing to 95% B at 20mn and maintaining 95% B until 23mn, then back to initial conditions in 1 min, Column kept at 40°C during all t). For the MS/MS detector, the parameters were scan between 100 and 1000m/z in both positive and negative mode for the first MS then automatic MS/MS for the first three most intense MS signal in both positive and negative mode at each cycle time (every second) (with isolation width = 2 and normalized collision energy at 35). We also used the same chromatographic conditions to quantify coumarins in samples using MS1 measured area corrected by Taxifolin (internal standard reference) area toward a standard curve ranging from 1.25 to 40 µM. Coumarins were quantified with the use of a standard curve.

### Chlorophyll content analysis

Rosettes were homogenized with mortars and tissue was transferred to a tube (∼50 mg or less whenever shoot mass wasn’t high enough). Chlorophylls were extracted by adding 500 µl of 80% acetone with 2.5 mM Na_2_CO_3_ followed by 10 min incubation in darkness at 4°C and centrifugation (10 min, 11 000 rcf). Supernatant was transferred to another tube. The process was repeated and the pooled supernatant was used to measure absorbance at 646.6, 663.6 and 750 nm. Chlorophyll concentrations were calculated according to (Porra et al., 1989). Samples were pooled together in case of *s8h-2* plants grown in FeCl_3_-supplemented media. n = 5 for *s8h-2* plants grown in FeCl_3_, n = 8 in all other cases.

### Data processing and statistical analysis

All treatments included at least 3 biological replicates. Data processing and statistical analysis were done in Excel 365 (Microsoft) or using R version 4.5.0. (R Core Team (2021), https://www.R-project.org/). Large language models (LLM) were used to support the process of writing of the R scripts for analysis and visualisation (GPT-5, Claude). Data visualisation was performed using R, BioEdit, BioRender and ChemDraw.

Kinetic parameters as well as optimal pH and temperature were determined from reactions performed in triplicate in two independent assays using different purified enzyme preparations. Results are expressed as means ± standard errors from three independent experiments. Kinetic data were fitted to the Michaelis–Menten equation, while optimal pH and temperature profiles were modeled using a two-parameter Gaussian function.

## SUPPLEMENTARY METHODS

### Homozygous knockout mutant isolation

Genotyping of homozygous mutants was performed by PCR. DNA was extracted from rosette leaves of Col-0 and mutant plants using a commercial DNA extraction kit (EURx Plant and Fungi DNA Purification Kit). PCR reaction was performed using EURx TiOptiTaq polymerase, reaction conditions are shown in **Supplementary Table S4**, reaction primers shown in **Supplementary Table S5**. Lack of *S6OD1* gene transcript in *s6od1-1* and *s6od1-2* and of *S6OD2* gene in *s6od2-1* and *s6od2-2* mutant background was confirmed by RT-PCR. RNA was extracted from Col-0 and mutant plant roots (for *s6od1* knockout) and from seeds (for *s6od2* knockout) using a commercial RNA extraction kit (EURx Universal RNA Purification Kit). Reverse transcription reactions were performed using LunaScript® RT Master Mix Kit (New England Biolabs), standardising the quantity by extracted RNA concentration. RT-PCR reactions were performed using DreamTaq Green DNA Polymerase (Thermo Fisher Scientific), reaction conditions are shown in **Supplementary Table S4**, reaction primers shown in Supplementary Table 3. PCR and RT-PCR reaction results were visualised by electrophoresis in 1 % agarose gel and ethidium bromide staining. The T-DNA insert localization was determined by sequencing, which was performed by an outside company (Genomed S.A., Warsaw) using primers as specified in **Supplementary Table S5**.

### In vitro liquid cultures

Arabidopsis seeds were surface sterilized by soaking in 70 % ethanol for 2 minutes, next in 5.25 % calcium hypochloride solution for 8 minutes followed by rinsing with sterile water 3 times. Next, they were suspended in 0.1 % agar and sown onto plates using 1 mL pipette tips. Seeds were sown onto Petri dishes containing 0.7 % agar half-strength Murashige and Skoog’s (MS) medium with macro-elements, micro-elements and vitamins in half-strength (Sigma-Aldrich). For stratification, plates were kept in the dark at 4 °C for 4 days and then placed under defined growth conditions (16 h light (∼5000 lux) at 22 °C and 8 h dark at 20 °C).

Ten days old seedlings from agar plates were transferred into 250 mL glass flasks containing 25 mL of sterile self-made 0.5 MS liquid medium (half strength Murashige and Skoog’s medium with macro-elements in half strength and micro-elements/vitamins in full strength), 5 seedlings per flask. Media contained 3 % sucrose, supplemented with 4 mg/L glycine, 200 mg/L myo-inositol, 1 mg/L thiamine hydrochloride, 0.5 mg/L pyridoxine hydrochloride and 0.5 mg/L nicotinic acid. No buffering agent was added and pH was adjusted to 5.7. Plants grown in liquid cultures were incubated on rotary platform shakers at 120 rpm. Two weeks after transfer, 5 mL of fresh medium was added to all flasks. After an additional week of culture, plants and growth media were harvested and flash frozen before processing. Roots of 5 plants grown together in a flask were treated as one sample.

### Construction of binary vector and Agrobacterium tumefaciens strains

*S6OD1* coding sequence was amplified (primer sequences shown in **Supplementary Table S6**) and cloned into pCR8 plasmid using the pCR®8/GW/TOPO® TA cloning kit (Invitrogen) (Vialart et al., 2012). The sequence was then subcloned into a pBIN vector using Gateway technology. The following pBIN:S6OD1 vector was then introduced into the GV3101 *A. tumefaciens* strain via electroporation and used for transient expression in *N. benthamiana* leaves.

### Transient expression of S6OD1 in tobacco leaves

*Nicotiana benthamiana* seeds were sown into pots and after 2 weeks transplanted and cultured for 3 additional weeks in growth chambers under a photoperiod of 16 h light (120 μmol m^−2^ s^−1^) at 24 °C and 8 h dark. N*. benthamiana* transformation was performed as described in Klińska-Bąchor et al., 2024. Briefly, *A. tumefaciens* carrying the pBIN:S6OD1 vector alongside strain containing p19 vector and strain containing vector carrying green fluorescent protein (GFP) were used to syringe-infiltrate N*. benthamiana* leaves. Plants inoculated with the *A. tumefaciens* suspension were then cultivated for another 5 days, after which expression of GFP was confirmed by examination under a binocular microscope. After this time, 3 cm diameter circles were cut out from the leaves and infiltrated with 500 µM solution of scopoletin or mock solution of water. Samples were flash frozen after 2 hours of incubation on the bench and later used for metabolite extraction. 8 samples from each condition were compared, each consisting of 1-2 leaf fragments.

### Col-0 and Stw-0 S6OD1 coding sequence alignment

Col-0 and Stw-0 *S6OD1* coding sequences were determined by sequencing (Genomed S.A., Warsaw). Translation and amino acid sequence alignment (**Supplementary Figures S33)** was performed with MEGA X programme with default settings.

## Acknowledgements

A.I. acknowledges support for the research of this work from the Polish National Science Centre [grant number UMO-2022/47/B/NZ2/01835] and the Polish National Agency for Academic Exchange [grant number BPN/BFR/2022/1/00039]. A.O. acknowledges support of the Scientific Council ‘Agronomie, Agroalimentaire et Forêt’, University of Lorraine.

## Contributions

A.D. designed and performed the experiments; collected samples; analyzed data; wrote and edited the manuscript; revised and approved the manuscript. C.C. performed experiments; analyzed data; edited the manuscript. I.P. collected samples; performed experiments; analyzed data. R. M. designed and performed the experiments; analyzed data; edited the manuscript. J.G. performed experiments. A.H. supervised the project. E.L. supervised the project; revised and approved the manuscript. A.I. designed the experiments; analyzed data; conceived and supervised the project; wrote and edited the manuscript; revised and approved the manuscript; provided funding. A.O. designed and performed the experiments; analyzed data; conceived and supervised the project, wrote the manuscript; revised and approved the manuscript; provided funding. All authors approved the final version of the manuscript.

## Data availability

All data generated and analysed in this study are included in this published article, and its Supplementary Data.

