## SUPPLEMENTARY FIGURES AND TABLES for "First *O*-demethylation activity in Arabidopsis specialized metabolism resolves the missing step in esculetin biosynthesis"

### Dobek et al.

### Supplementary Figures and Tables

**Table S1.** Pairwise primary structure identity **(A)** and similarity **(B)** among 5 AtODs (S6OD1, S6OD2, AtOD3, AtOD4, AtOD5) determined by ClustalW multiple alignment. (A) Identity dataset showing values ranging from 63 % to 85 %. (B) Similarity dataset showing values ranging from 81 % to 91 %.

**(A)**

| | **AtOD2 (S6OD2)** | **AtOD3** | **AtOD4** | **AtOD1 (S6OD1)** | **AtOD5** |
|---|---|---|---|---|---|
| **AtOD2 (S6OD2)** | **100** | **77** | **72** | **72** | **63** |
| **AtOD3** | **77** | **100** | **75** | **75** | **66** |
| **AtOD4** | **72** | **75** | **100** | **85** | **65** |
| **AtOD1 (S6OD1)** | **72** | **75** | **85** | **100** | **65** |
| **AtOD5** | **63** | **66** | **65** | **65** | **100** |

**(B)**

| | **AtOD2 (S6OD2)** | **AtOD3** | **AtOD4** | **AtOD1 (S6OD1)** | **AtOD5** |
|---|---|---|---|---|---|
| **AtOD2 (S6OD2)** | 100 | 87 | 85 | 84 | 82 |
| **AtOD3** | 87 | 100 | 88 | 86 | 82 |
| **AtOD4** | 85 | 88 | 100 | 91 | 81 |
| **AtOD1 (S6OD1)** | 84 | 86 | 91 | 100 | 82 |
| **AtOD5** | 82 | 82 | 81 | 82 | 100 |

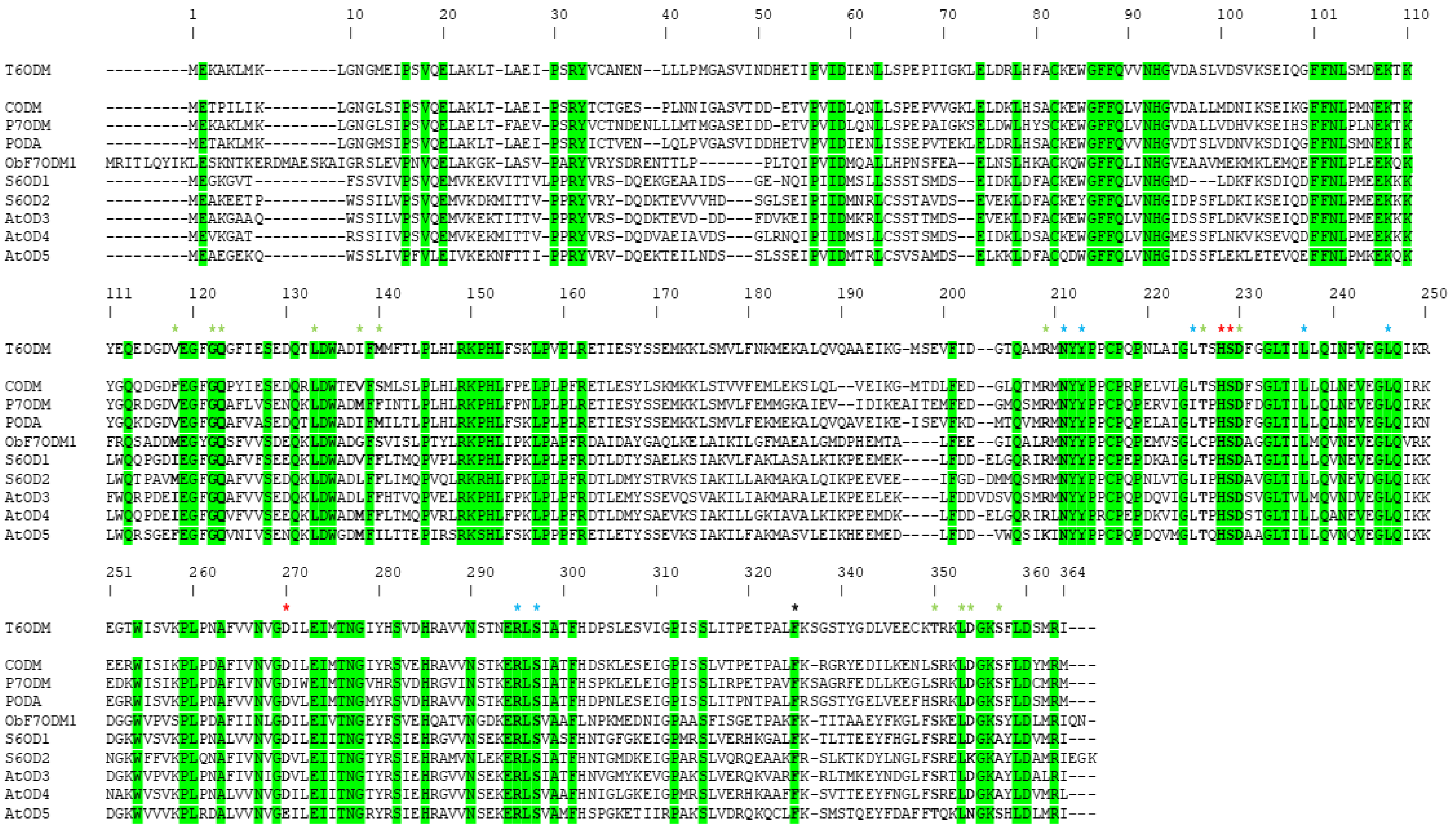

**Figure S1. Alignment of the primary structures of previously characterized 2-OGDs from *Papaver somniferum* (Hagel et al., 2010; Farrow et al., 2013; Farrow et al., 2015) and *Ocimum basilicum* (Berim et al., 2014), alongside members of the *Arabidopsis thaliana* DOXC52 *O*-demethylase subfamily.** T6ODM: thebaine *O*-demethylase; CODM: codeine *O*-demethylase; P7ODM: papaverine *O*-demethylase; PODA: protopine *O*-demethylase; ObF7ODM1: gardenin B 7-*O*-demethylase / 8-hydroxysalvigenin-7-*O*-demethylase; S6OD1: scopoletin 6-*O*-demethylase 1; S6OD2: scopoletin 6-*O*-demethylase 2; AtOD3–5*:* Arabidopsis *O*-demethylases 3 to 5. Amino-acid numbering refers to T6ODM. Red stars mark residues of the catalytic triad; blue stars indicate residues involved in α-ketoglutarate and succinate binding; green stars indicate residues involved in substrate recognition. Amino acids highlighted in green represent canonical residues.

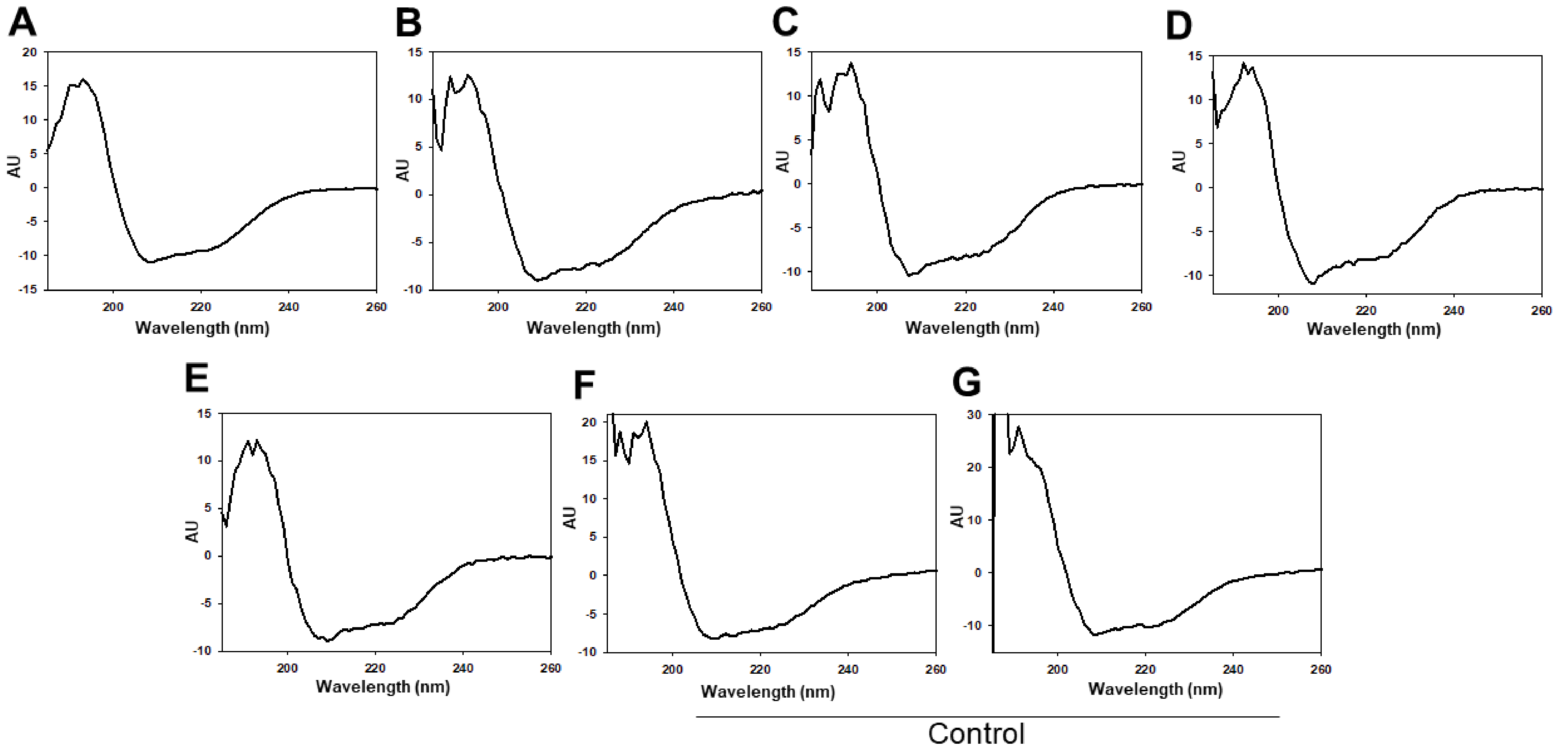


**Figure S2. CD spectra of O-demethylases. A: S6OD1, B: S6OD2, C: AtOD3, D: AtOD4, E: AtOD5, F: S8H under Native condition, G: S8H under renaturated condition.** CD spectra were scanned from 260 to 190 nm in potassium phosphate buffer 10 mM (pH 7.1) at 25 °C, and at an enzyme concentration of 50 µM.

**Table S2.** Prediction of secondary structure of S6OD1 to AtOD5 under renaturated condition; S8H Native and S8H under renaturated condition.

| | | | | | | Control | |
|---|---|---|---|---|---|---|---|
| **Enzymes** | **S6COD1** | **S6COD2** | **AtOD3** | **AtOD4** | **AtOD5** | **S8H Native** | **S8H Renatured** |
| % α-helix | 28.8 | 21.7 | 22.3 | 22.5 | 21.6 | 22.8 | 25.3 |
| % extended strand | 23.3 | 38.1 | 37.0 | 36.5 | 39.0 | 34.1 | 28.1 |
| % Others | 41.1 | 58.5 | 57.3 | 57.1 | 58.1 | 58.6 | 56.2 |

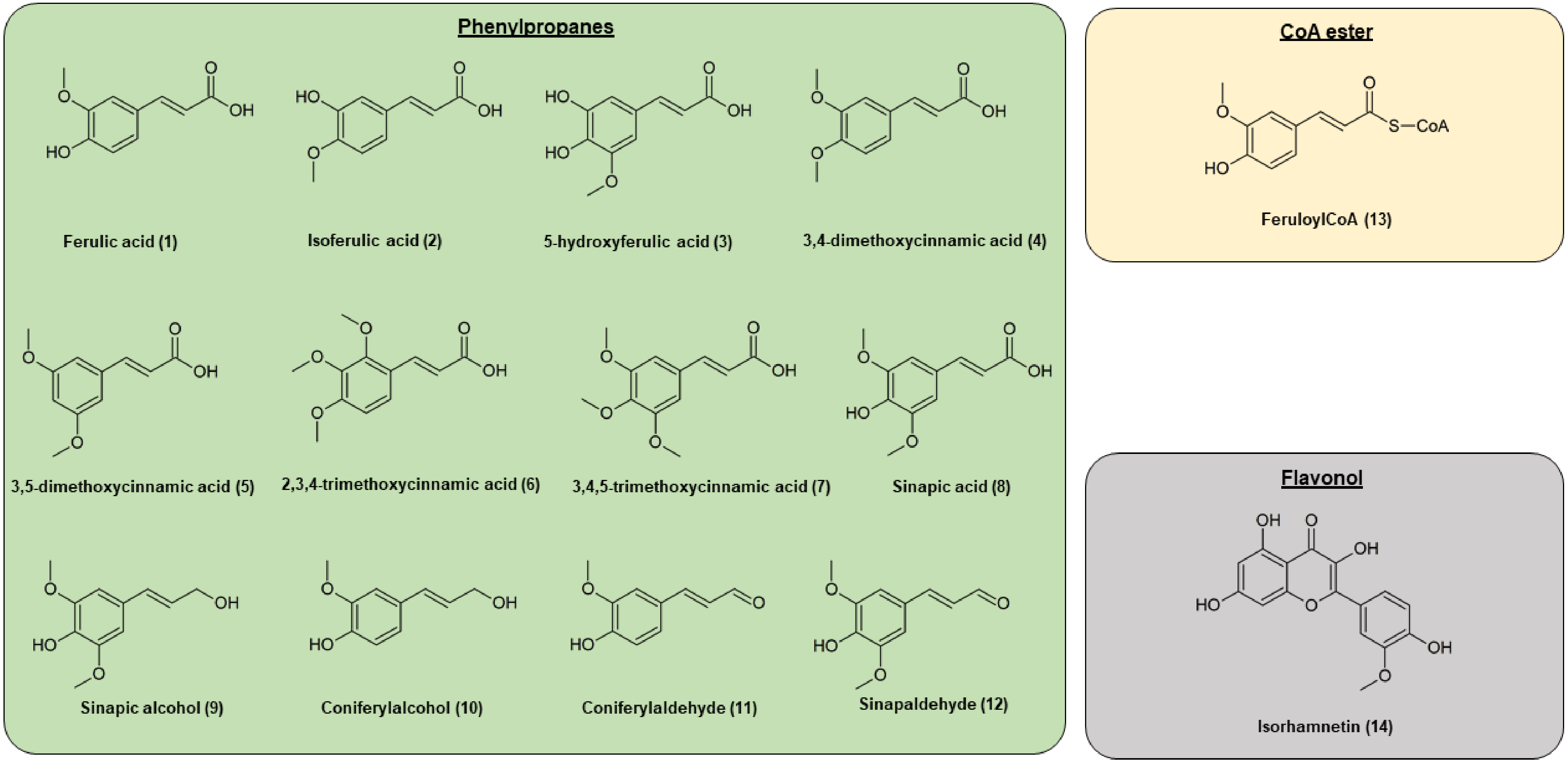


**Figure S3. Chemical structures of the *O*-methylated compounds tested as substrates of DOXC52 subclass of *Arabidopsis thaliana*.** The panel illustrates the set of molecules used to assess the catalytic activity of S6OD1, S6OD2, AtOD3, AtOD4, AtOD5. Structures include phenylpropans (green box), CoA ester feruloyl CoA (yellow box), flavonoid isorhamnetin (grey box) selected for their structural diversity and relevance in the Arabidopsis phenylpropanoid pathway.

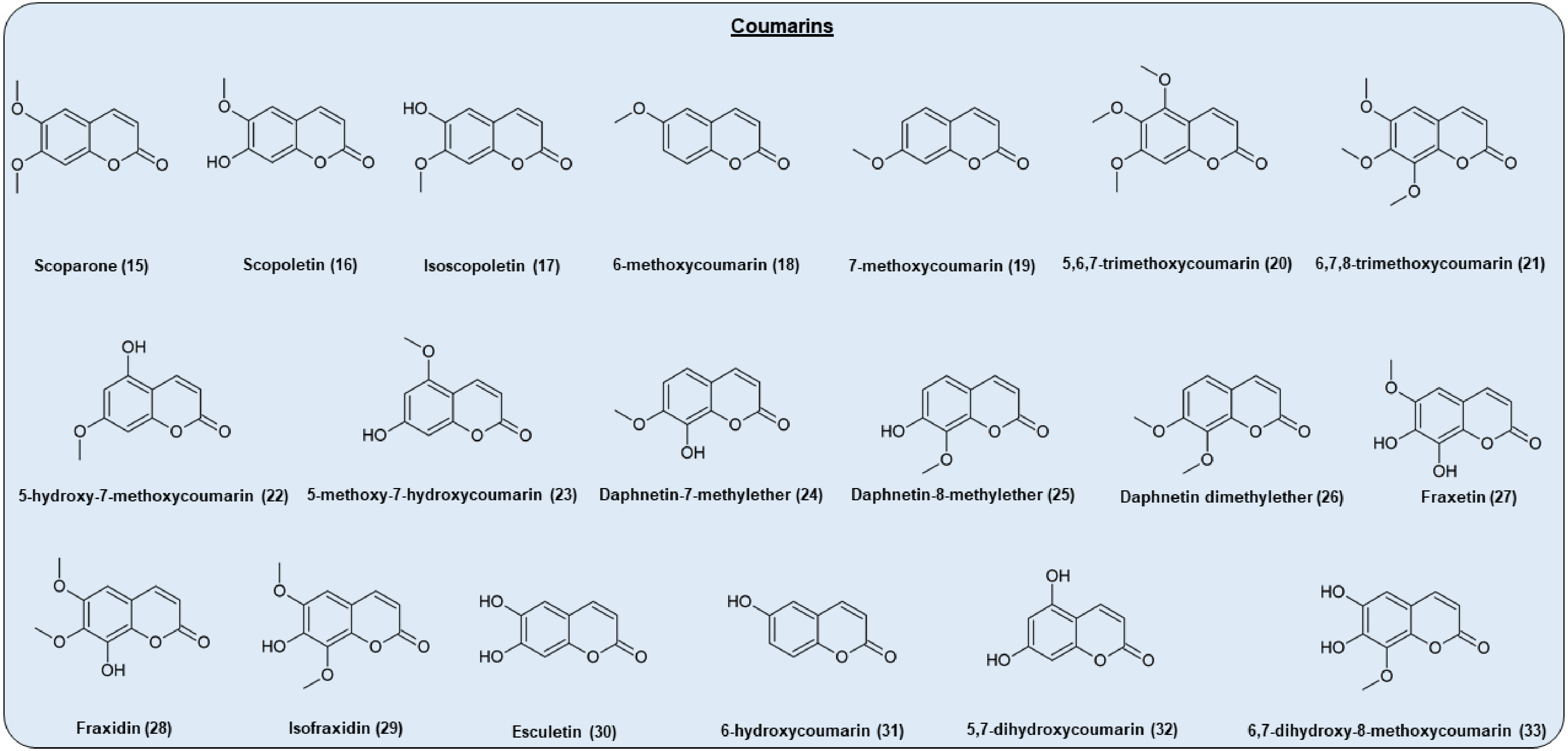


**Figure S4. Chemical structures of the *O*-methylated coumarins tested as substrates of DOXC52 subclass of *Arabidopsis thaliana*.** The panel illustrates the set of molecules used to assess the catalytic activity of S6OD1, S6OD2, AtOD3, AtOD4, AtOD5. Coumarins were selected for their structural diversity and relevance in the Arabidopsis phenylpropanoid pathway.

**Table S3. Kinetics of *O*-demethylation activities of SOD1,SOD2,AtOD3,AtOD4.** The reactions were performed with the purified enzymes, which enabled calculation of the relative *Vmax* values of the different substrates. The concentrations of substrates ranged from 0.1 to 10 Km depending on the substrate and the enzyme. The concentration of α-ketoglutarate and ascorbic acid were set at 500 µM, the concentration of $FeSO_4$ was set at 5 mM. All results are expressed as means ± standard errors of three independent experiments. The units of $V_{max}$ and $K_m$ are pmol.sec$^{-1}$.µg of purified enzyme and µM, respectively.

| | Substrate | 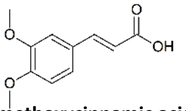 3,4-dimethoxycinnamic acid | 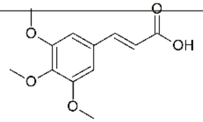 3,4,5-trimethoxycinnamic acid | Scoparone | 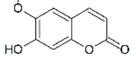 Scopoletin | 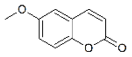 6-methoxycoumarin | 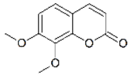 Daphnetin dimethyether |
|---|---|---|---|---|---|---|---|
| | Product | 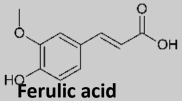 Ferulic acid | 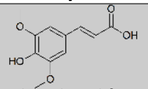 Sinapic acid | 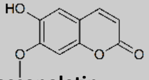 Isoscopoletin | 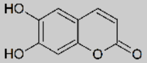 Esculetin | 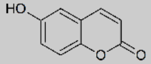 6-hydroxyxycoumarin | 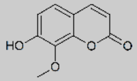 Daphnetin 8-methylether |
| | Position ring | 4 | 4 | 6 | 6 | 6 | 6 |
| S6OD1 | $V_{max}$ | 316.7 +/- 38.8 | N.A. | 1543 +/- 195 | 1604 +/- 196 | 460.2 +/- 28.9 | N.A. |
| | $K_{m\ app}$ | 203.8 +/- 46.7 | N.A. | 3.0 +/- 1.3 | 3.4 +/- 1.3 | 21.9 +/- 5.2 | N.A. |
| | $V_{max}$ / $K_{m\ app}$ | 1.5 +/- 3.0 | N.A. | 514.3 +/- 200.5 | 471.7 +/- 271,4 | 21.0 +/- 8,3 | N.A. |
| S6OD2 | $V_{max}$ | 175.3 +/- 6.8 | 385.4 +/- 31.0 | 119.3 +/- 5.9 | 103.7 +/- 5.0 | N.A. | N.A. |
| | $K_{m\ app}$ | 4.5 +/- 0.8 | 5.4 +/- 1.8 | 3.7 +/- 0.9 | 2.1 +/- 0.5 | N.A. | N.A. |
| | $V_{max}$ / $K_{m\ app}$ | 38.9 +/- 10.3 | 71.3 +/- 44.0 | 32.2 +/- 12.5 | 49.4 +/- 18,5 | N.A. | N.A. |
| AtOD3 | $V_{max}$ | 2266 +/- 237 | N.A. | 66.2 +/- 33.4 | N.A. | N.A. | N.A. |
| | $K_{m\ app}$ | 160.5 +/- 33.9 | N.A. | 335.6 +/- 270.1 | N.A. | N.A. | N.A. |
| | $V_{max}$ / $K_{m\ app}$ | 14.1 +/- 5.6 | N.A. | 0.02 +/- 0.03 | N.A. | N.A. | N.A. |
| AtOD4 | $V_{max}$ | 602 +/- 52 | 951.2 +/- 72.8 | 159 +/- 40 | N.A. | N.A. | 291.3 +/- 9.8 |
| | $K_{m\ app}$ | 77.3 +/- 16.9 | 38.2 +/-9.3 | 126.2 +/- 69.9 | N.A. | N.A. | 63.8 +/- 5.8 |
| | $V_{max}$ / $K_{m\ app}$ | 7.8 +/- 3.0 | 24.9 +/- 10.5 | 1.3 +/- 2.2 | N.A. | N.A. | 4.6 +/- 0.6 |
| AtOD5 | $V_{max}$ | N.A. | N.A. | N.A. | N.A. | N.A. | N.A. |
| | $K_{m\ app}$ | N.A. | N.A. | N.A. | N.A. | N.A. | N.A. |
| | $V_{max}$ / $K_{m\ app}$ | N.A. | N.A. | N.A. | N.A. | N.A. | N.A. |

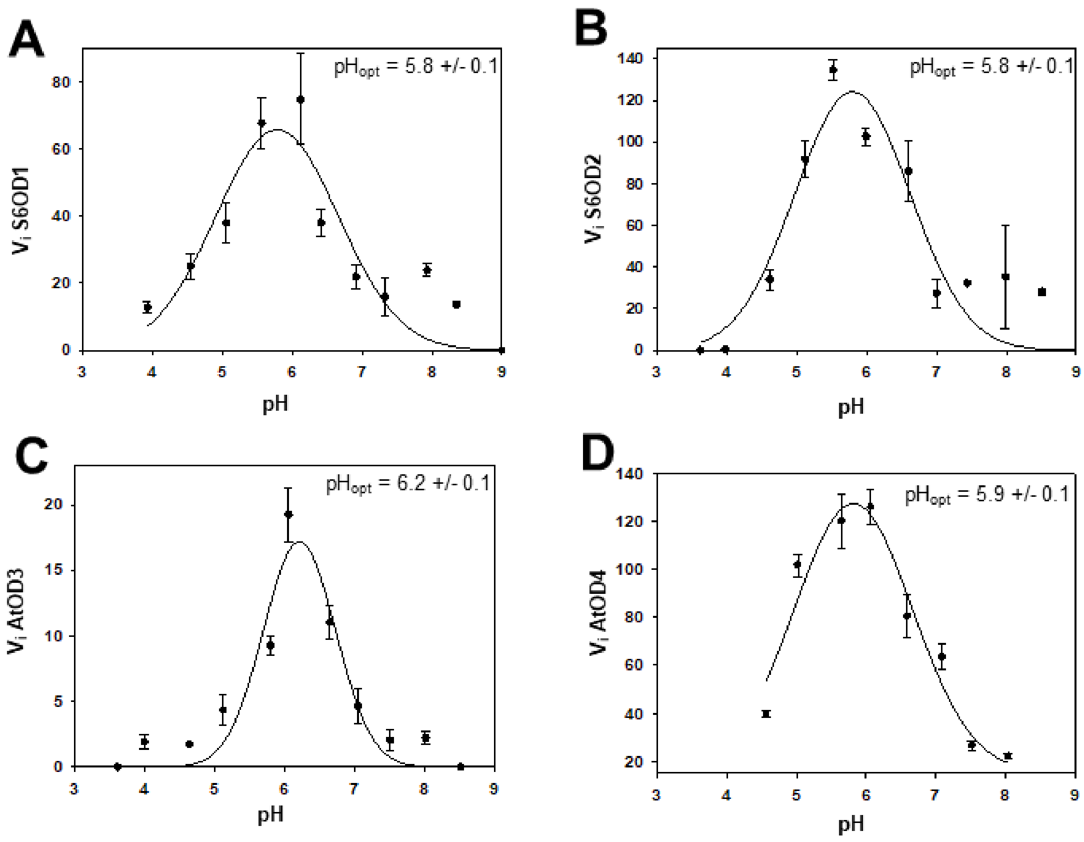


**Figure S5. pH dependence of enzymatic activity.** Relative activities of the four enzymes (S6OD1, S6OD2, AtOD3 and AtOD4) were measured at optimal temperature across a pH gradient. Reaction mixtures (100 µL) contained purified enzyme, 500 µM α-ketoglutarate, 500 µM ascorbic acid, 5 mM $FeSO_4$, and 200 µM scoparone, and were incubated in a mixed buffer system (0.1 M acetic acid, 0.1 M Tris, 40 mM imidazole). Products were monitored at their maximal absorbance wavelength. Data points represent mean values of three independent measurements; error bars indicate standard error. Curves were fitted using SigmaPlot to estimate pH–activity profiles. The unit of $V_i$ is pmol.sec$^{-1}$.µg of purified enzyme.

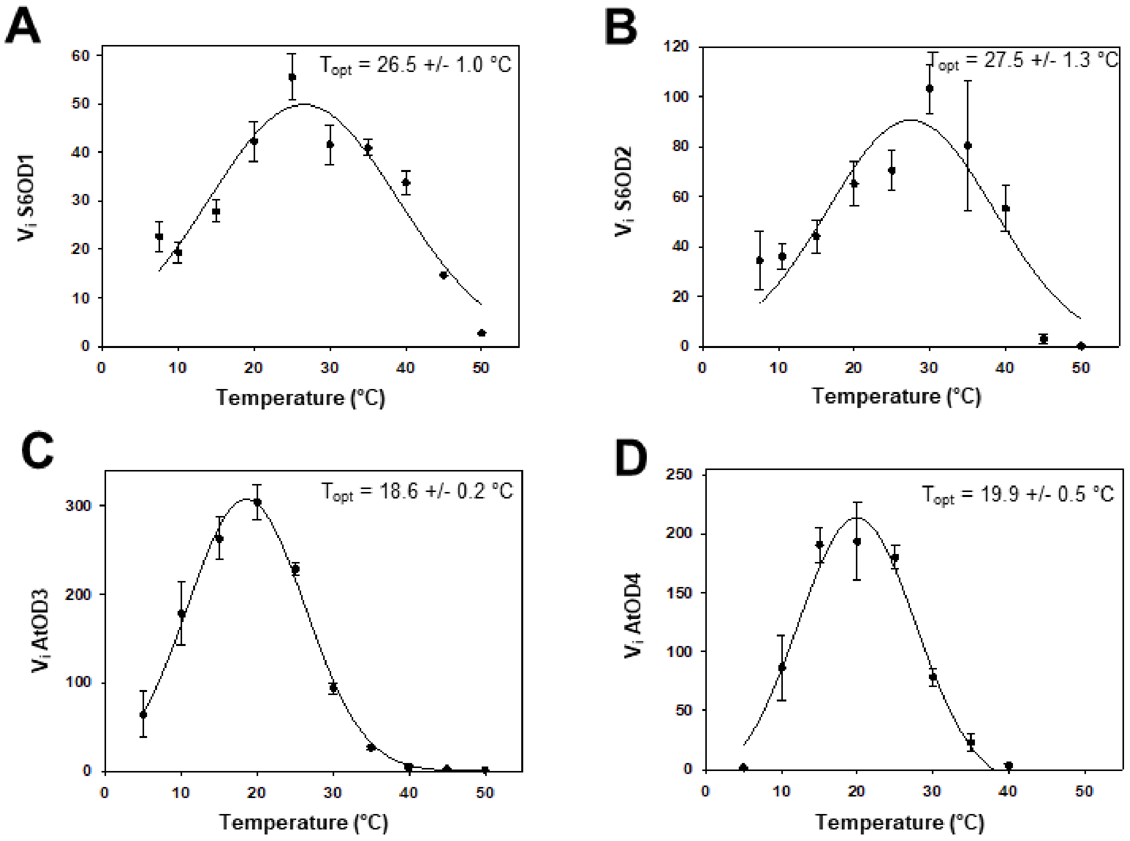


**Figure S6. Temperature dependence of enzymatic activity.** Relative activities of the four enzymes (S6OD1, S6OD2, AtOD3 and AtOD4) were measured at optimal pH across a temperature gradient. Reaction mixtures (100 µL) contained purified enzyme, 500 µM α-ketoglutarate, 500 µM ascorbic acid, 5 mM $FeSO_4$, and 200 µM scoparone, and were incubated in a polybuffer system (0.1 M acetic acid, 0.1 M Tris, 40 mM imidazole). Products were detected at their maximal absorbance wavelength. Data points represent mean values of three independent measurements; error bars indicate standard error. Curves were fitted using SigmaPlot to estimate pH–activity profiles. The unit of $V_i$ is pmol.sec$^{-1}$.µg of purified enzyme.

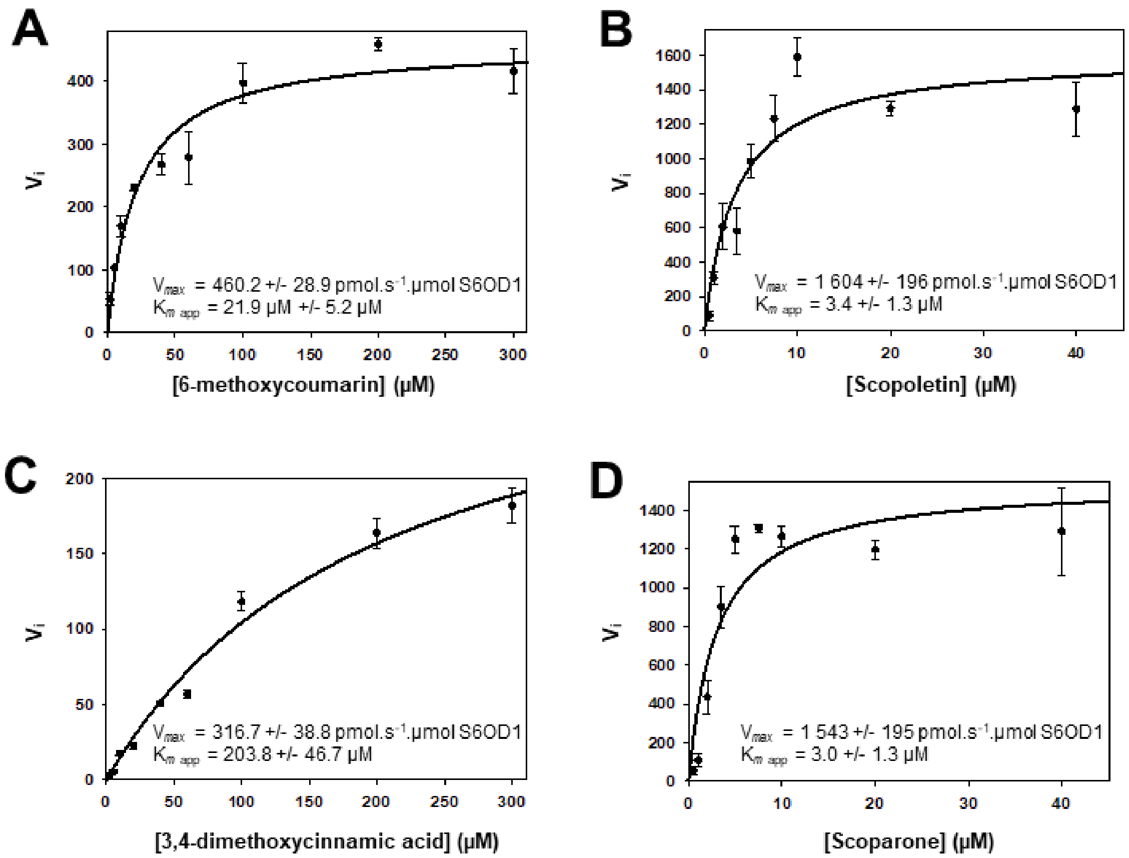


**Figure S7. Michaelis Menten regression of S6OD1 metabolization of A) 6-methoxycoumarin, B) Scopoletin, C) 3,4-dimethoxycinnamic acid and D) Scoparone substrates.** Reactions were carried out at optimal pH and temperature (Supplementary Fig. SX and SX) in polybuffer Tris 0.1 M acetic acid 0.1 M Imidazole 40 mM. The concentrations of substrates ranged from 1–300 µM for 6-methoxycoumarin and 3,4-dimethoxycinnamic acid, 5–40 µM for scopoletin and scoparone. The concentration of α-ketoglutarate and ascorbic acid were set at 500 µM, the concentration of $FeSO_4$ was set at 5 mM. All results are expressed as means ± standard errors of three independent experiments. Figures presented were obtained by fitting the experimental data to the Michaelis-Menten equation. The unit of $V_i$ is pmol.sec$^{-1}$.µg of purified enzyme.

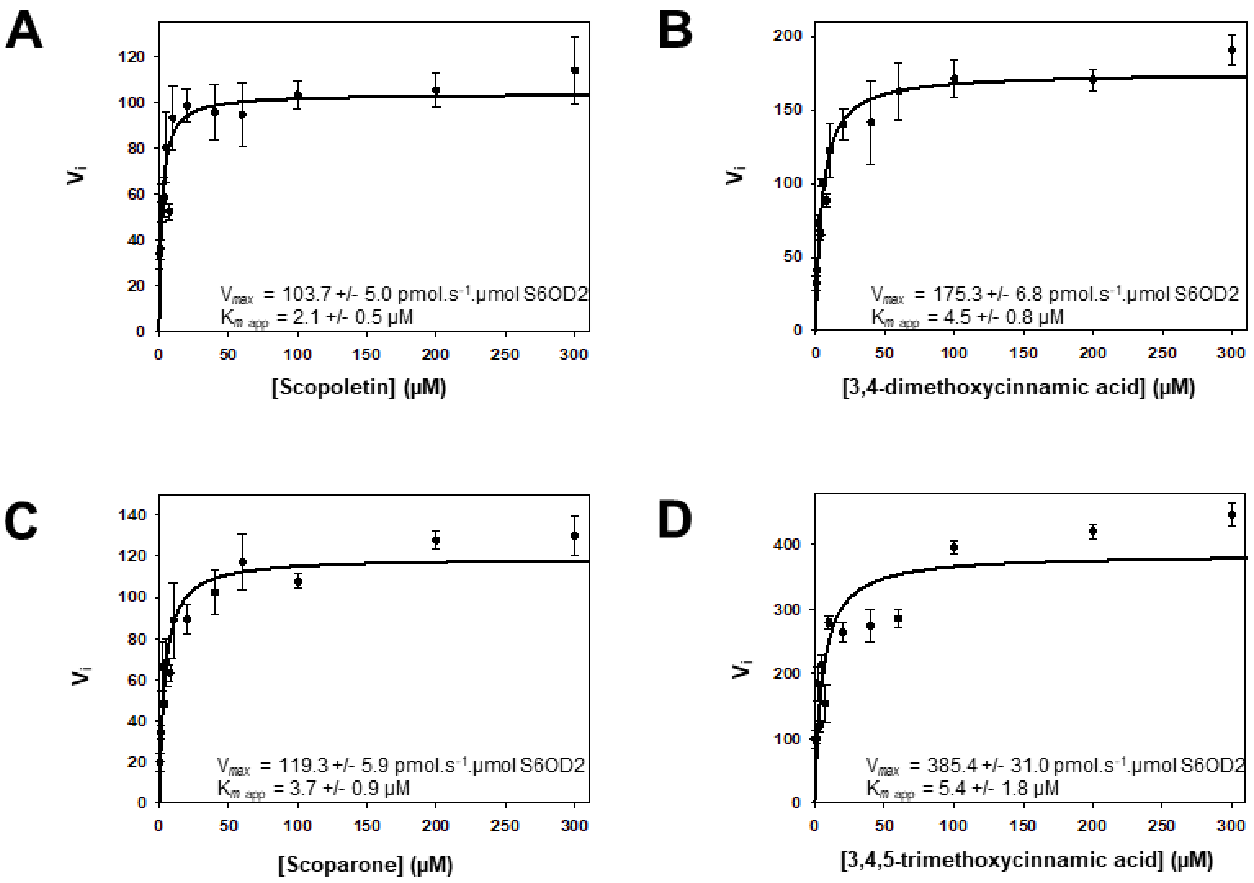


**Figure S8. Michaelis Menten regression of S6OD2 metabolization of A) Scopoletin, B) 3,4-dimethoxycinnamic acid, C) scoparone and D) 3,4,5-trimethoxycinnamic acid substrates.** Reactions were carried out at optimal pH and temperature (Supplementary Fig. SX and SX) in polybuffer Tris 0.1 M acetic acid 0.1 M Imidazole 40 mM. The concentrations of substrates ranged from 1–300 µM for scopoletin, scoparone, 3,4-dimethoxycoumarin and 3,4,5-trimethoxycoumarin. The concentration of α-ketoglutarate and ascorbic acid were set at 500 µM, the concentration of $FeSO_4$ was set at 5 mM. All results are expressed as means ± standard errors of three independent experiments. Figures presented were obtained by fitting the experimental data to the Michaelis-Menten equation. The unit of $V_i$ is pmol.sec$^{-1}$.µg of purified enzyme.

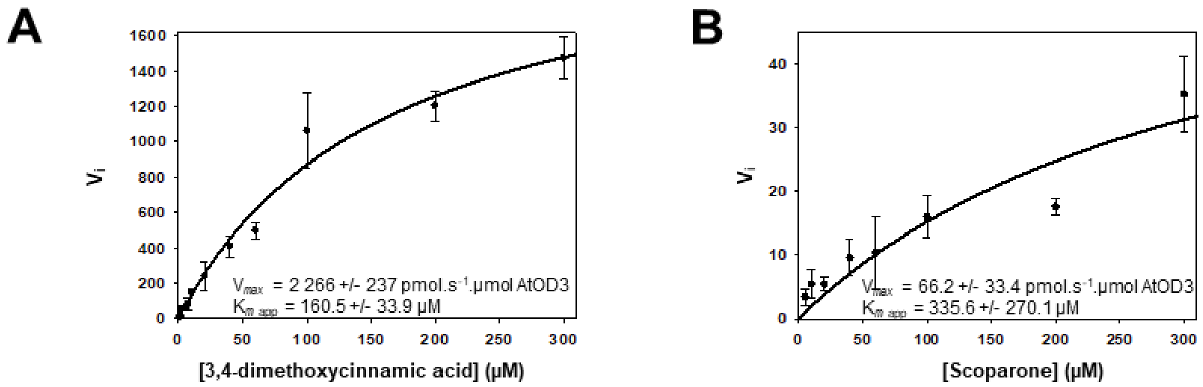


**Figure S9. Michaelis Menten regression of S6OD2 metabolization of A) 3,4-dimethoxycinnamic acid and B) scoparone substrates.** Reactions were carried out at optimal pH and temperature (Supplementary Fig. SX and SX) in polybuffer Tris 0.1 M acetic acid 0.1 M Imidazole 40 mM. The concentrations of substrates ranged from 1–300 µM for scoparone and 3,4-dimethoxycoumarin. The concentration of α-ketoglutarate and ascorbic acid were set at 500 µM, the concentration of $FeSO_4$ was set at 5 mM. All results are expressed as means ± standard errors of three independent experiments. Figures presented were obtained by fitting the experimental data to the Michaelis-Menten equation. The unit of $V_i$ is pmol.sec$^{-1}$.µg of purified enzyme.

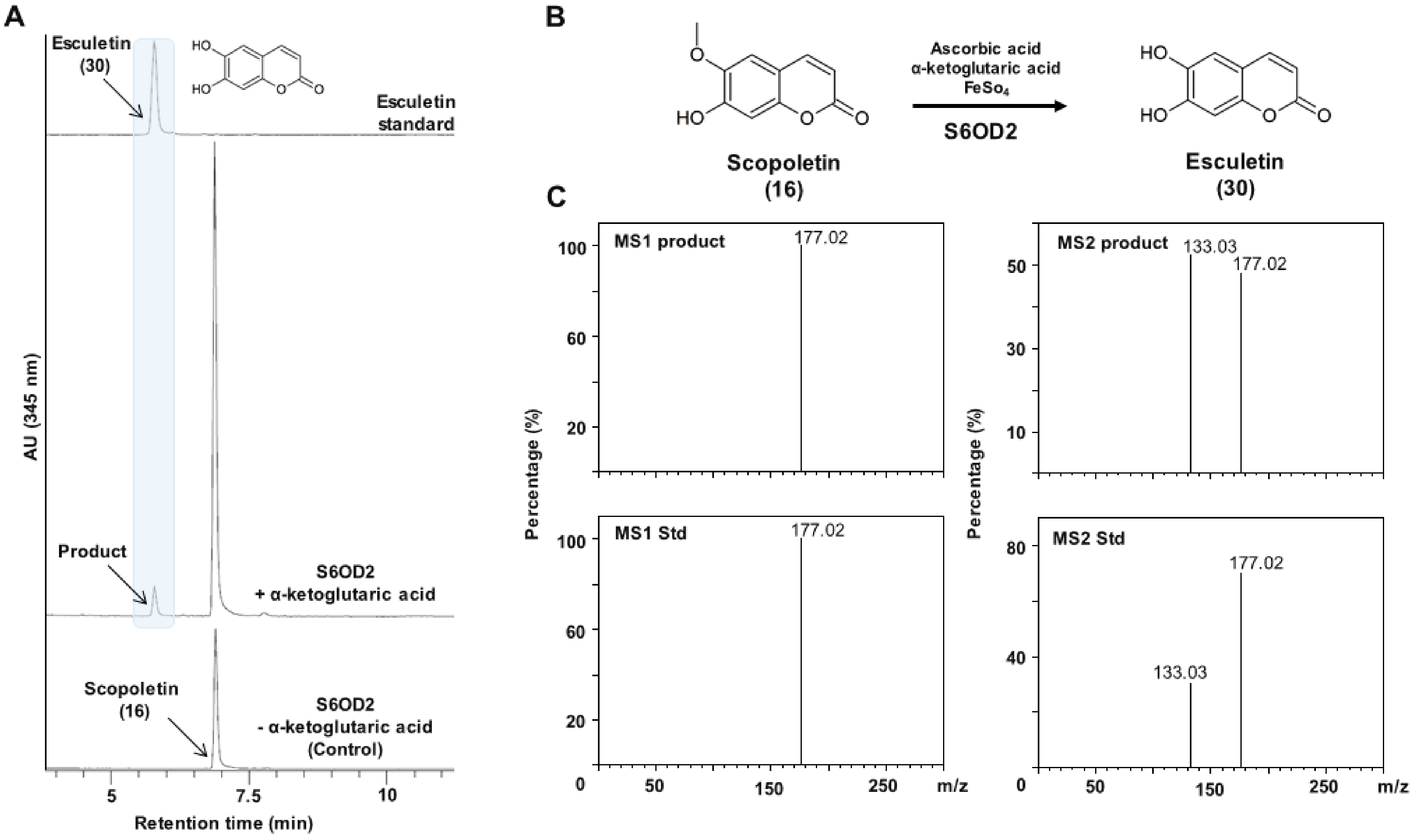


**Figure S10. Biochemical characterization of the *O*-demethylase activity of S6OD2 towards scopoletin.** (A) UV chromatograms at 345 nm of enzymatic reactions using renatured sonication pellets containing recombinant S6OD2. Reaction mixtures (100 µL) contained purified enzyme, 500 µM α-ketoglutarate, 500 µM ascorbic acid, 5 mM $FeSO_4$, and 200 µM *O*-methylated substrate, and were incubated in a mixed buffer system (acetic acid 0.1 M, Tris 0.1 M, Imidazole 40 mM) under optimal temperature and pH conditions (Supplementary Figures X and Y) for 20 h. B) Schematic representation of the reaction converting scopoletin to esculetin. (C) Comparison of $MS^1$ and $MS^2$ spectra of the reaction product with those of authentic ferulic acid standard in negative ion mode using Orbitrap detection.

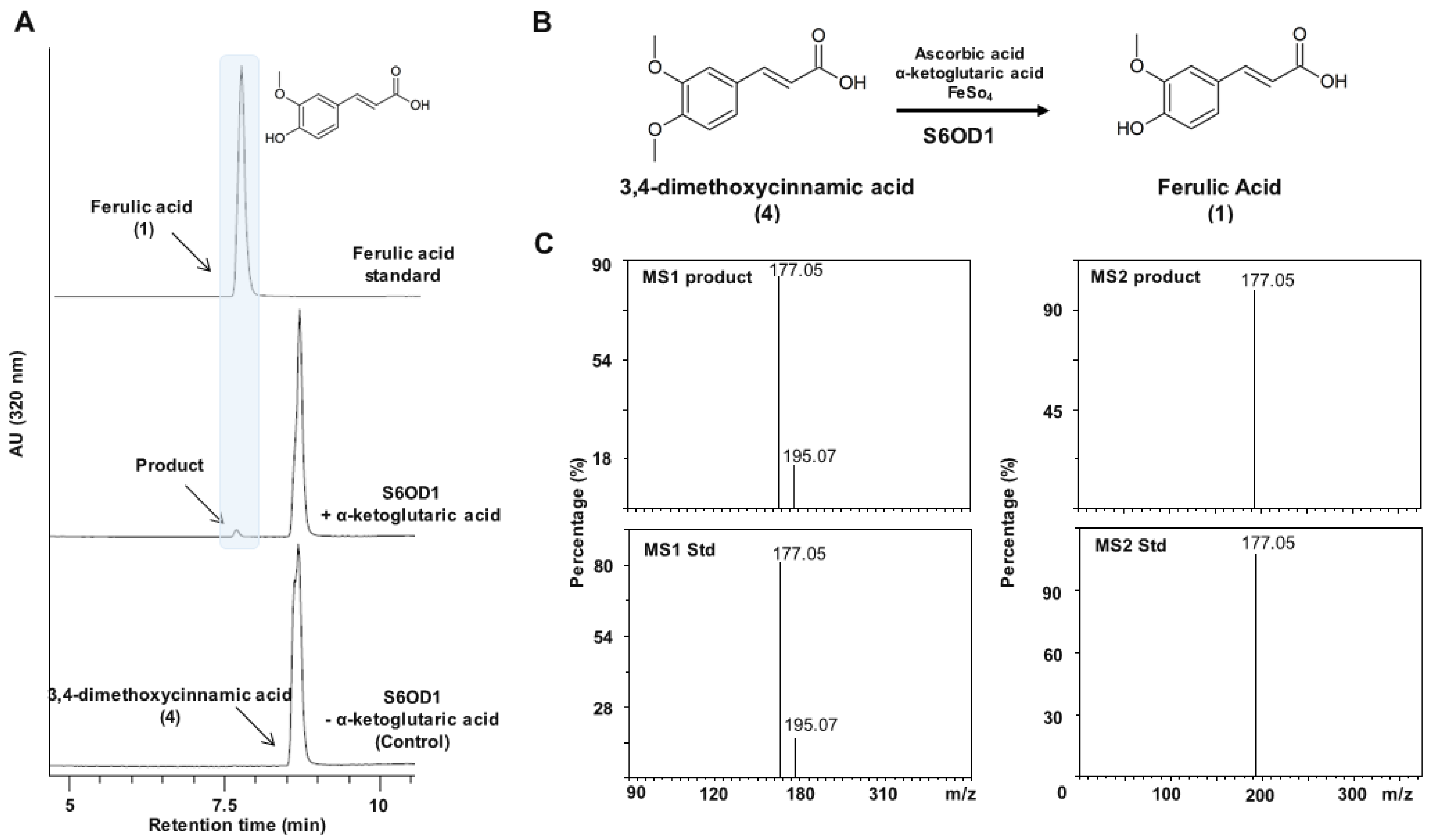


**Figure S11. Biochemical characterization of the *O*-demethylase activity of S6OD1 towards 3,4-dimethoxycinnamic acid.** (A) UV chromatograms at 320 nm of enzymatic reactions using renatured sonication pellets containing recombinant S6OD1. Reaction mixtures (100 µL) contained purified enzyme, 500 µM α-ketoglutarate, 500 µM ascorbic acid, 5 mM $FeSO_4$, and 200 µM *O*-methylated substrate, and were incubated in a mixed buffer system (acetic acid 0.1 M, Tris 0.1 M, Imidazole 40 mM) under optimal temperature and pH conditions (Supplementary Figures X and Y) for 20 h. B) Schematic representation of the reaction converting 3,4-dimethoxycinnamic acid to ferulic acid. (C) Comparison of $MS^1$ and $MS^2$ spectra of the reaction product with those of authentic ferulic acid standard in negative ion mode using Orbitrap detection.

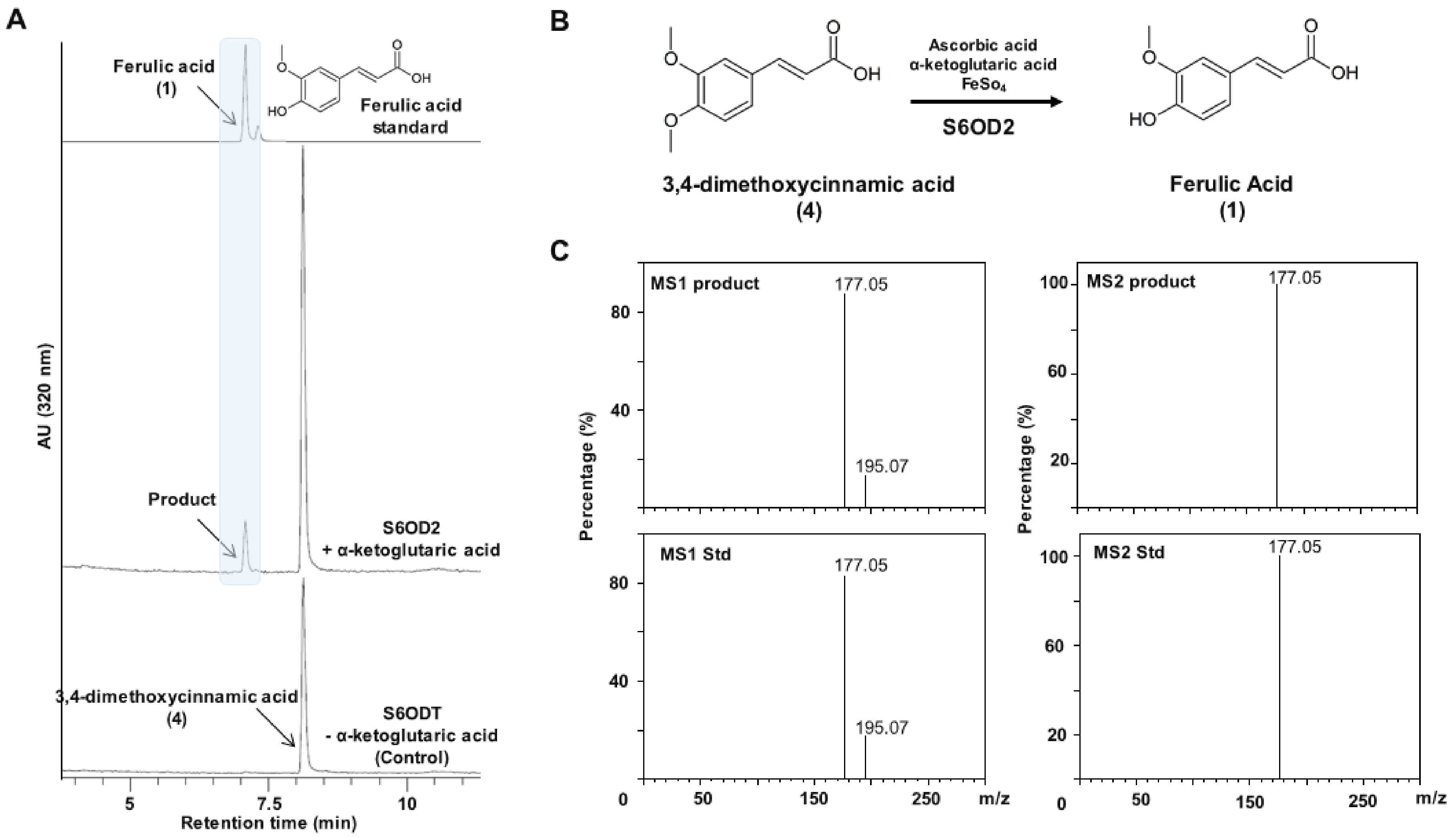


**Figure S12. Biochemical characterization of the *O*-demethylase activity of S6OD2 towards 3,4-dimethoxycinnamic acid.** (A) UV chromatograms at 320 nm of enzymatic reactions using renatured sonication pellets containing recombinant S6OD2. Reaction mixtures (100 µL) contained purified enzyme, 500 µM α-ketoglutarate, 500 µM ascorbic acid, 5 mM $FeSO_4$, and 200 µM *O*-methylated substrate, and were incubated in a mixed buffer system (acetic acid 0.1 M, Tris 0.1 M, Imidazole 40 mM) under optimal temperature and pH conditions (Supplementary Figures X and Y) for 20 h. B) Schematic representation of the reaction converting 3,4-dimethoxycinnamic acid to ferulic acid. (C) Comparison of $MS^1$ and $MS^2$ spectra of the reaction product with those of authentic ferulic acid standard in negative ion mode using Orbitrap detection.

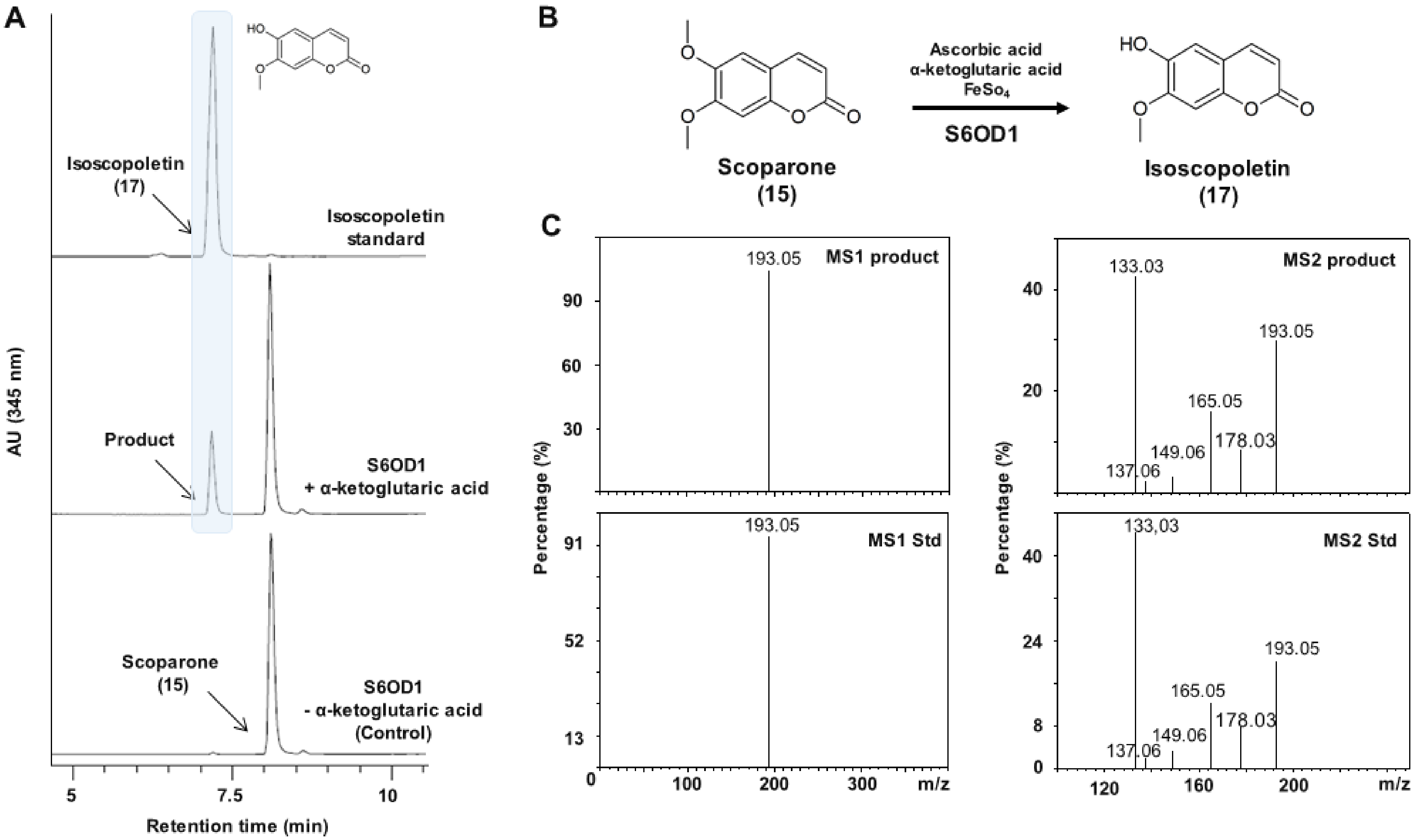


**Figure S13. Biochemical characterization of the *O*-demethylase activity of S6OD1 towards scoparone.** (A) UV chromatograms at 345 nm of enzymatic reactions using renatured sonication pellets containing recombinant S6OD1. Reaction mixtures (100 µL) contained purified enzyme, 500 µM α-ketoglutarate, 500 µM ascorbic acid, 5 mM $FeSO_4$, and 200 µM *O*-methylated substrate, and were incubated in a mixed buffer system (acetic acid 0.1 M, Tris 0.1 M, Imidazole 40 mM) under optimal temperature and pH conditions (Supplementary Figures X and Y) for 20 h. B) Schematic representation of the reaction converting scoparone to isoscopoletin. (C) Comparison of $MS^1$ and $MS^2$ spectra of the reaction product with those of authentic isoscopoletin standard in negative ion mode using Orbitrap detection.

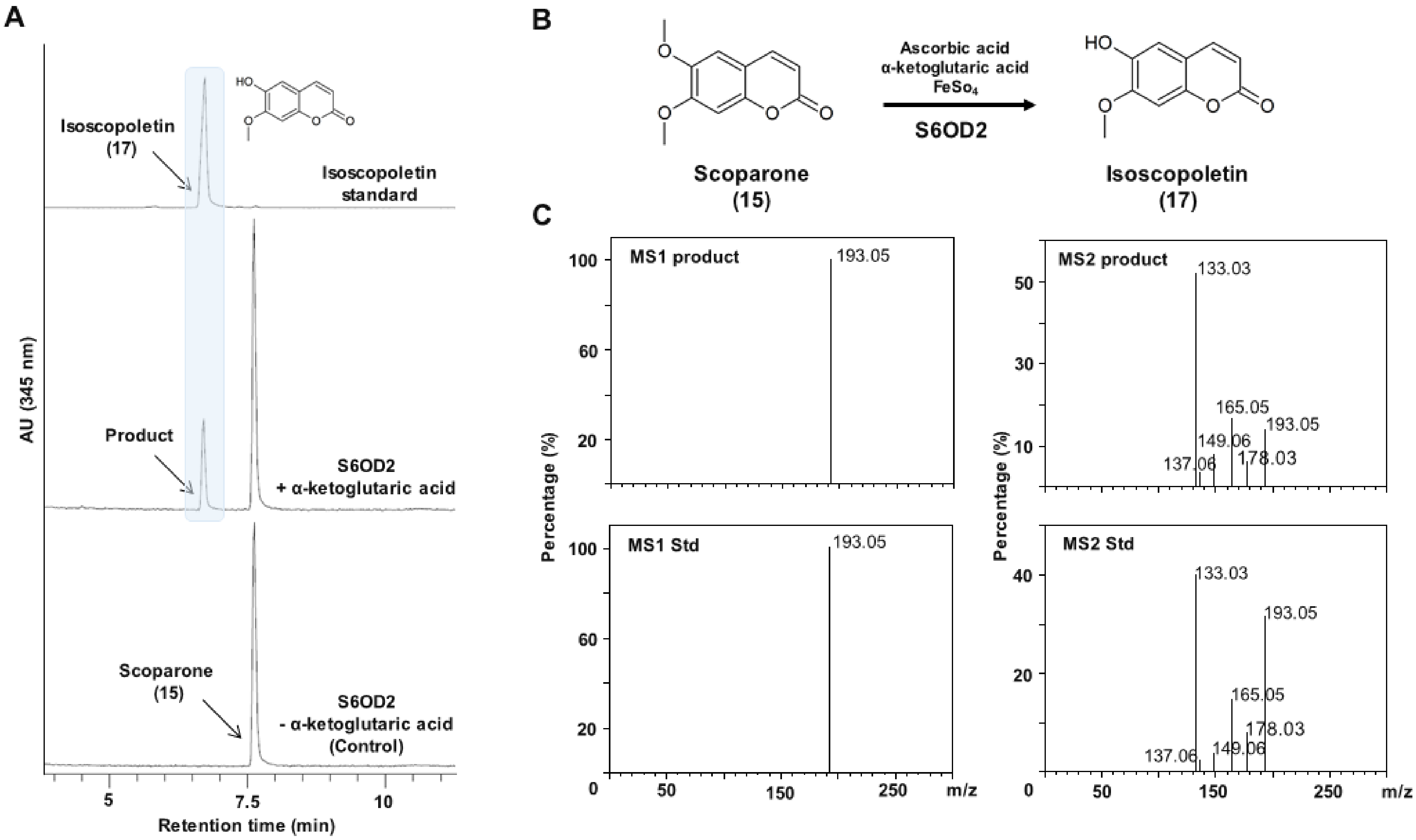


**Figure S14. Biochemical characterization of the *O*-demethylase activity of S6OD2 towards scoparone.** (A) UV chromatograms at 345 nm of enzymatic reactions using renatured sonication pellets containing recombinant S6OD2. Reaction mixtures (100 µL) contained purified enzyme, 500 µM α-ketoglutarate, 500 µM ascorbic acid, 5 mM $FeSO_4$, and 200 µM *O*-methylated substrate, and were incubated in a mixed buffer system (acetic acid 0.1 M, Tris 0.1 M, Imidazole 40 mM) under optimal temperature and pH conditions (Supplementary Figures X and Y) for 20 h. B) Schematic representation of the reaction converting scoparone to isoscopoletin. (C) Comparison of $MS^1$ and $MS^2$ spectra of the reaction product with those of authentic isoscopoletin standard in negative ion mode using Orbitrap detection.

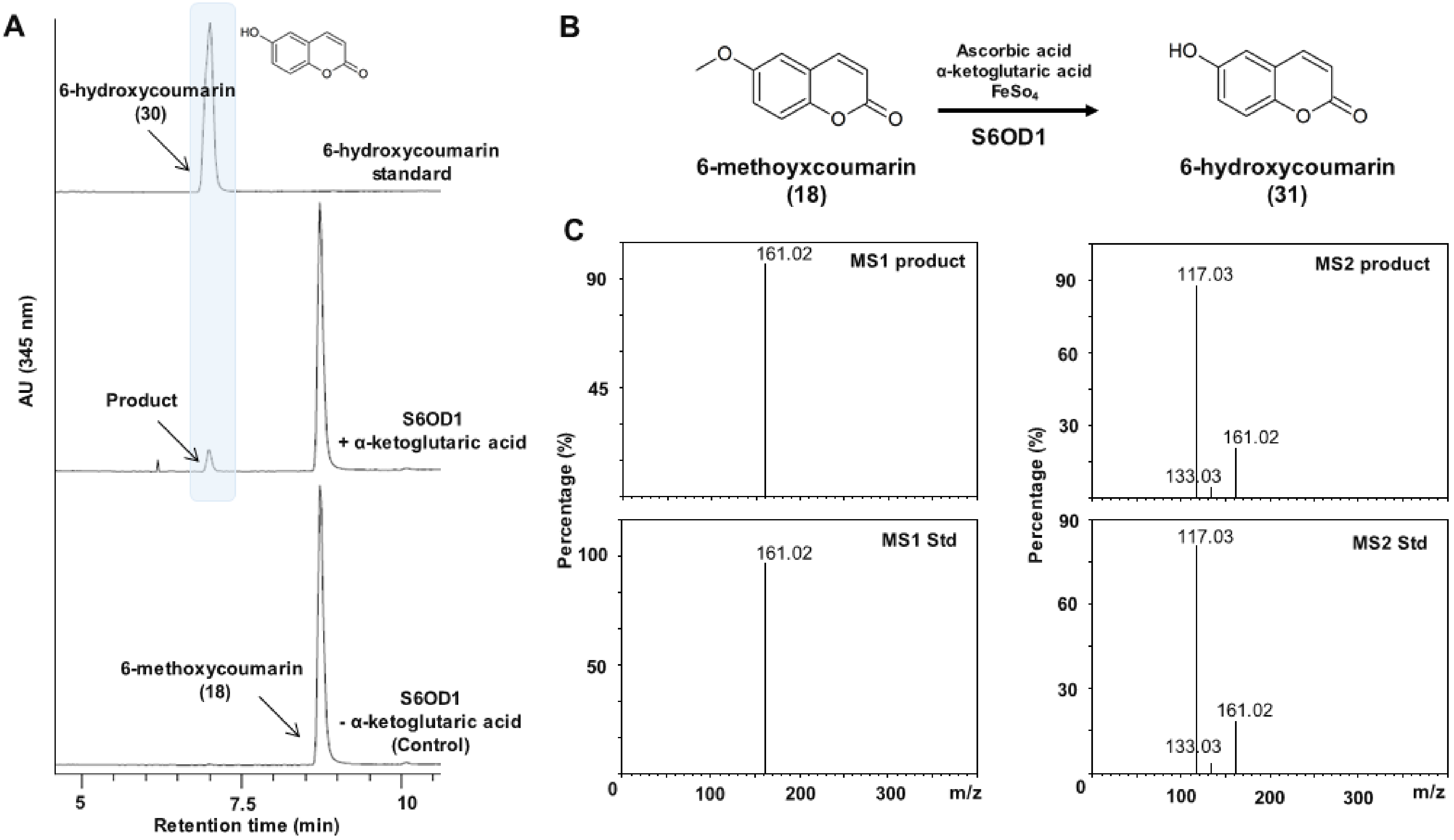


**Figure S15. Biochemical characterization of the *O*-demethylase activity of S6OD1 towards 6-methoxycoumarin.** (A) UV chromatograms at 345 nm of enzymatic reactions using renatured sonication pellets containing recombinant S6OD1. Reaction mixtures (100 µL) contained purified enzyme, 500 µM α-ketoglutarate, 500 µM ascorbic acid, 5 mM $FeSO_4$, and 200 µM *O*-methylated substrate, and were incubated in a mixed buffer system (acetic acid 0.1 M, Tris 0.1 M, Imidazole 40 mM) under optimal temperature and pH conditions (Supplementary Figures X and Y) for 20 h. B) Schematic representation of the reaction converting 6-methoxycoumarin to 6-hydroxycoumarin. (C) Comparison of $MS^1$ and $MS^2$ spectra of the reaction product with those of authentic 6-hydroxycoumarin standard in negative ion mode using Orbitrap detection.

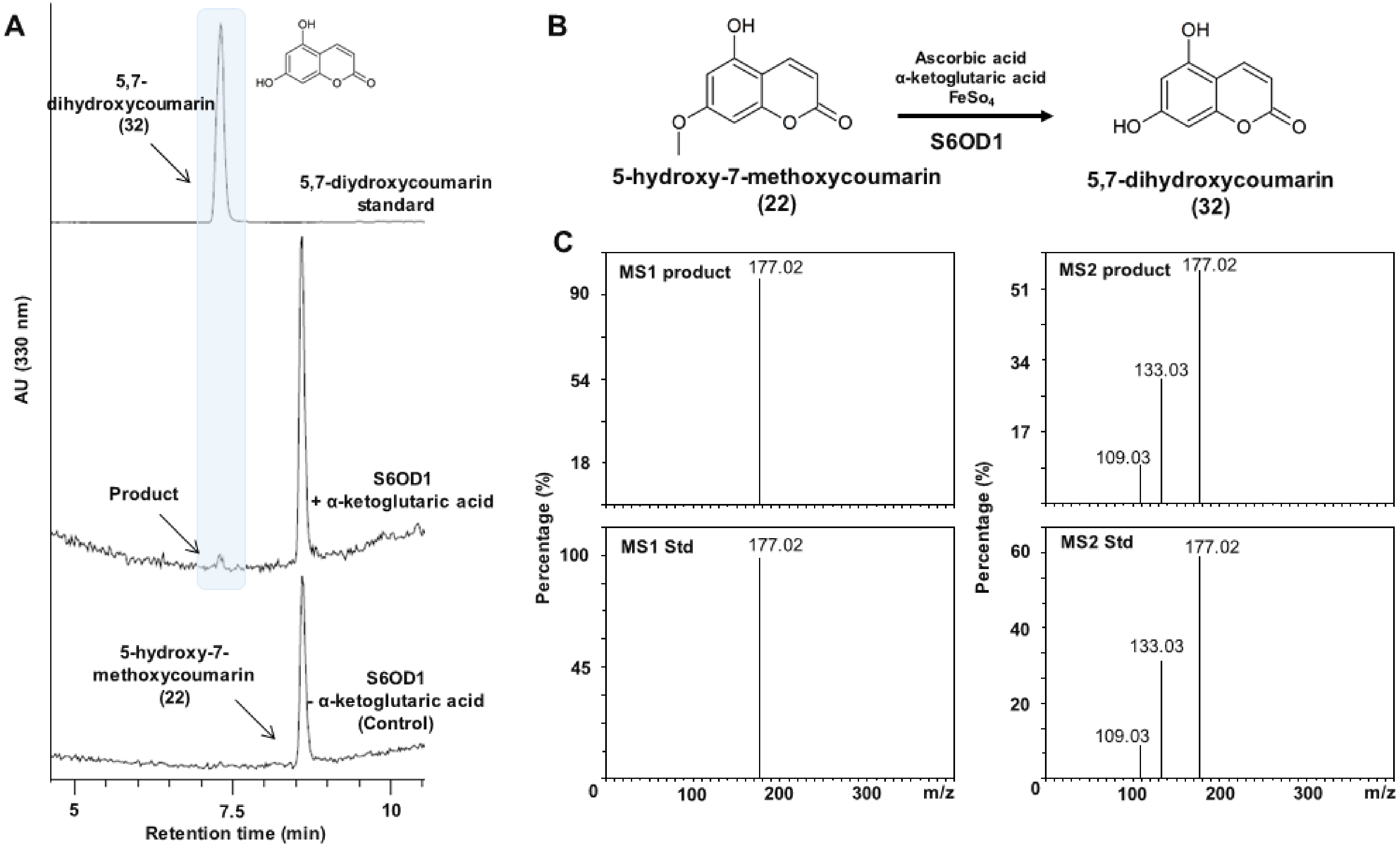


**Figure S16. Biochemical characterization of the *O*-demethylase activity of S6OD1 towards 5-hydroxy-7-methoxycoumarin.** (A) UV chromatograms at 330 nm of enzymatic reactions using renatured sonication pellets containing recombinant S6OD1. Reaction mixtures (100 µL) contained purified enzyme, 500 µM α-ketoglutarate, 500 µM ascorbic acid, 5 mM $FeSO_4$, and 200 µM *O*-methylated substrate, and were incubated in a mixed buffer system (acetic acid 0.1 M, Tris 0.1 M, Imidazole 40 mM) under optimal temperature and pH conditions (Supplementary Figures X and Y) for 20 h. B) Schematic representation of the reaction converting 5-hydroxy-7-methoxycoumarin to 5,7-dihydroxycoumarin. (C) Comparison of $MS^1$ and $MS^2$ spectra of the reaction product with those of authentic 5,7-dihydroxycoumarin standard in negative ion mode using Orbitrap detection.

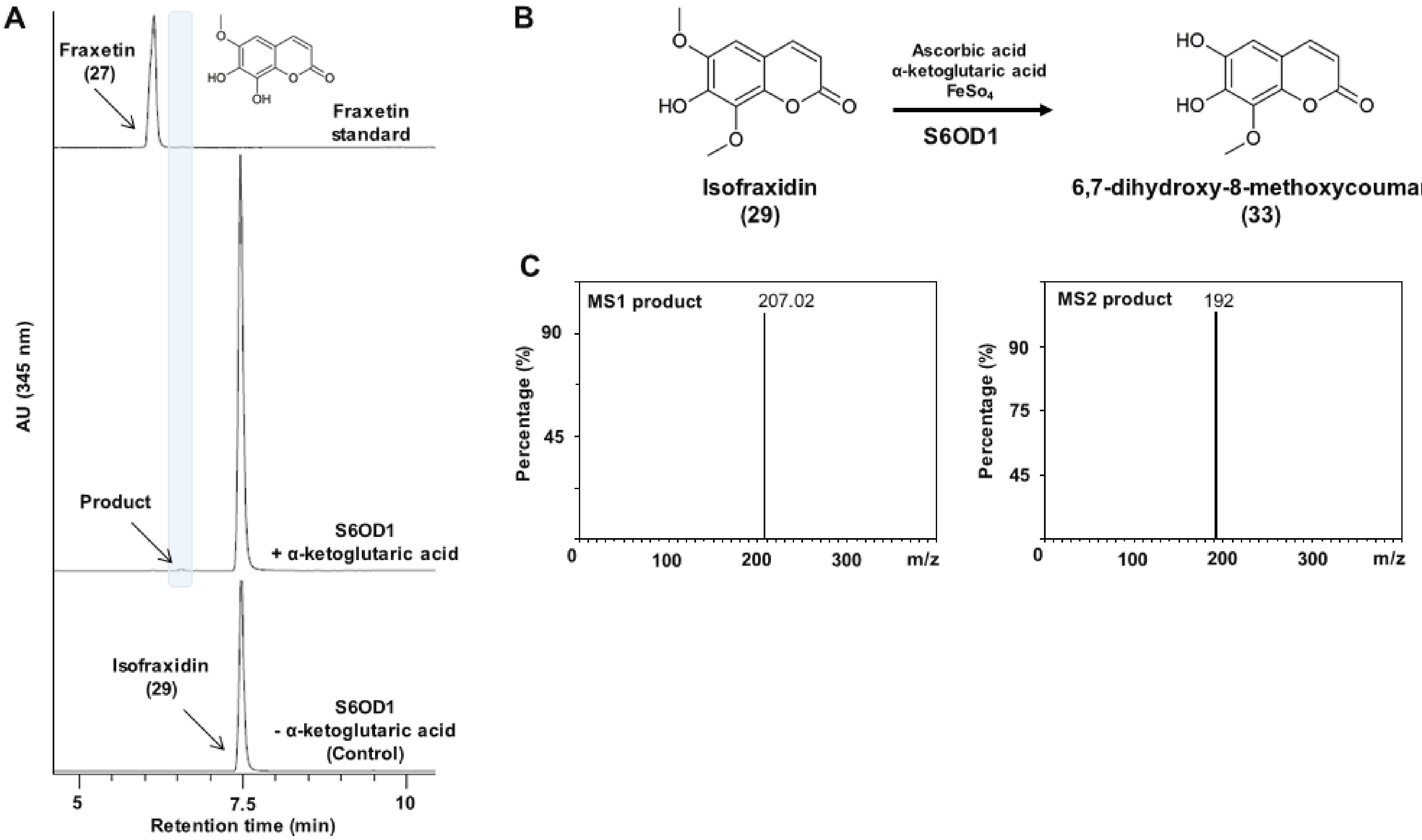


**Figure S17. Biochemical characterization of the *O*-demethylase activity of S6OD1 towards 3,4-dimethoxycinnamic acid.** (A) UV chromatograms at 345 nm of enzymatic reactions using renatured sonication pellets containing recombinant S6OD1. Reaction mixtures (100 µL) contained purified enzyme, 500 µM α-ketoglutarate, 500 µM ascorbic acid, 5 mM $FeSO_4$, and 200 µM *O*-methylated substrate, and were incubated in a mixed buffer system (acetic acid 0.1 M, Tris 0.1 M, Imidazole 40 mM) under optimal temperature and pH conditions (Supplementary Figures X and Y) for 20 h. B) Schematic representation of the reaction converting isofraxidin to 6,7-8-methoxycouymarin. (C) Comparison of $MS^1$ and $MS^2$ spectra of the reaction product in negative ion mode using Orbitrap detection.

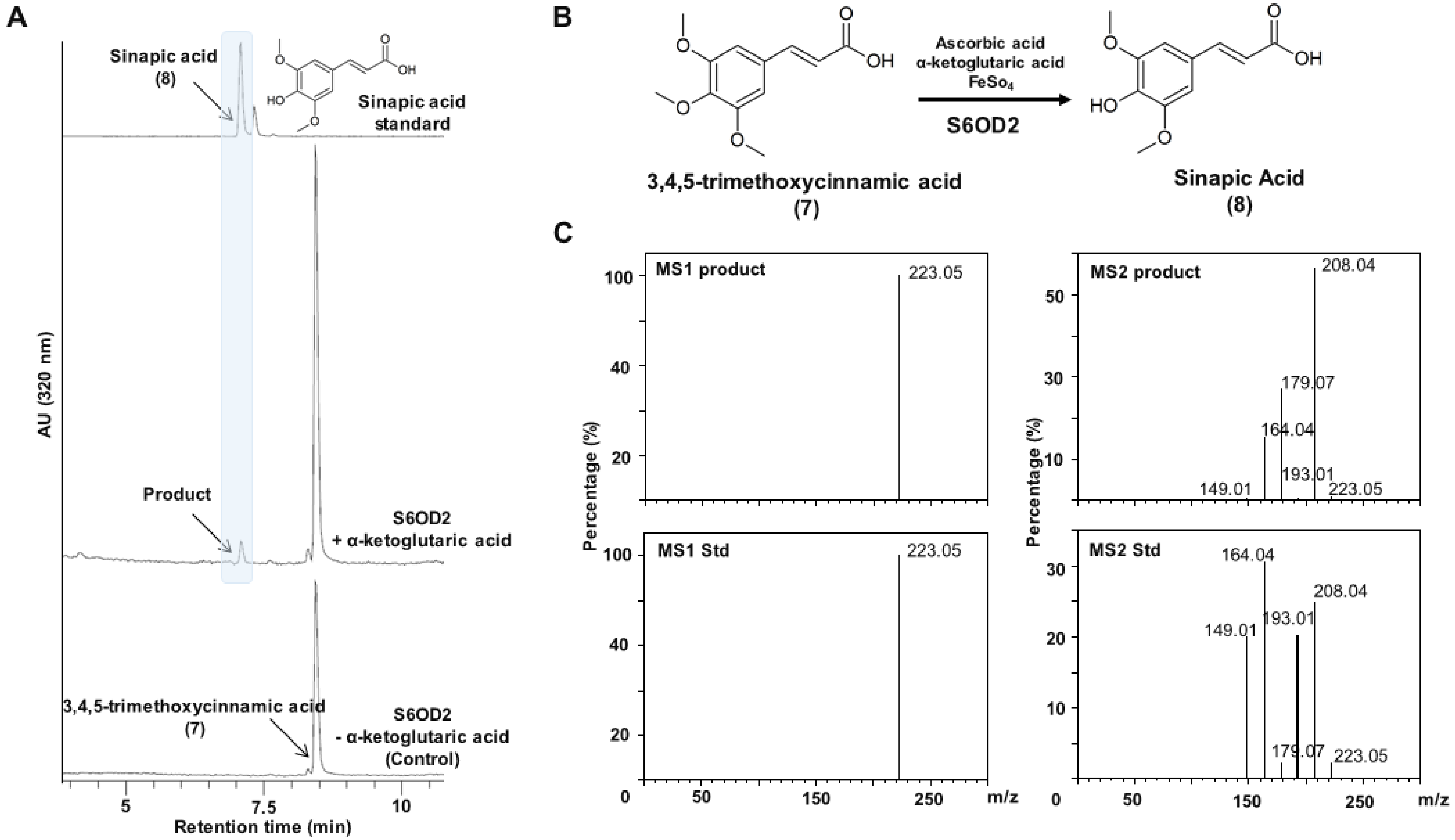


**Figure S18. Biochemical characterization of the *O*-demethylase activity of S6OD2 towards 3,4,5-trimethoxycinnamic acid.** (A) UV chromatograms at 320 nm of enzymatic reactions using renatured sonication pellets containing recombinant S6OD2. Reaction mixtures (100 µL) contained purified enzyme, 500 µM α-ketoglutarate, 500 µM ascorbic acid, 5 mM $FeSO_4$, and 200 µM *O*-methylated substrate, and were incubated in a mixed buffer system (acetic acid 0.1 M, Tris 0.1 M, Imidazole 40 mM) under optimal temperature and pH conditions (Supplementary Figures X and Y) for 20 h. B) Schematic representation of the reaction converting 3,4,5-trimethoxycinnamic acid to sinapic acid. (C) Comparison of $MS^1$ and $MS^2$ spectra of the reaction product with those of authentic sinapic acid standard in negative ion mode using Orbitrap detection.

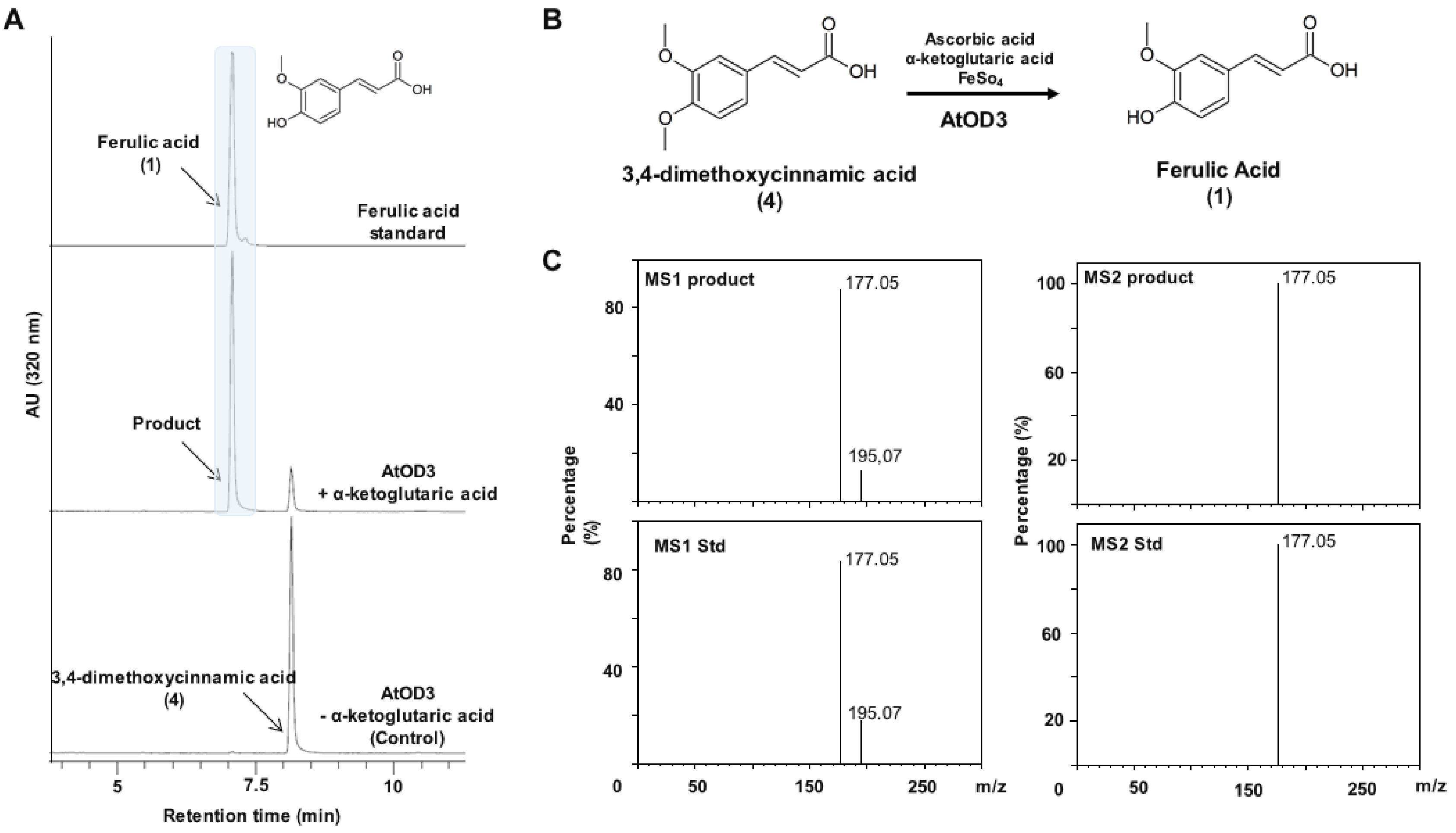


**Figure S19. Biochemical characterization of the *O*-demethylase activity of AtOD3 towards 3,4-dimethoxycinnamic acid.** (A) UV chromatograms at 320 nm of enzymatic reactions using renatured sonication pellets containing recombinant AtOD3. Reaction mixtures (100 µL) contained purified enzyme, 500 µM α-ketoglutarate, 500 µM ascorbic acid, 5 mM $FeSO_4$, and 200 µM *O*-methylated substrate, and were incubated in a mixed buffer system (acetic acid 0.1 M, Tris 0.1 M, Imidazole 40 mM) under optimal temperature and pH conditions (Supplementary Figures X and Y) for 20 h. B) Schematic representation of the reaction converting 3,4-dimethoxycinnamic acid to ferulic acid. (C) Comparison of $MS^1$ and $MS^2$ spectra of the reaction product with those of authentic ferulic acid standard in negative ion mode using Orbitrap detection.

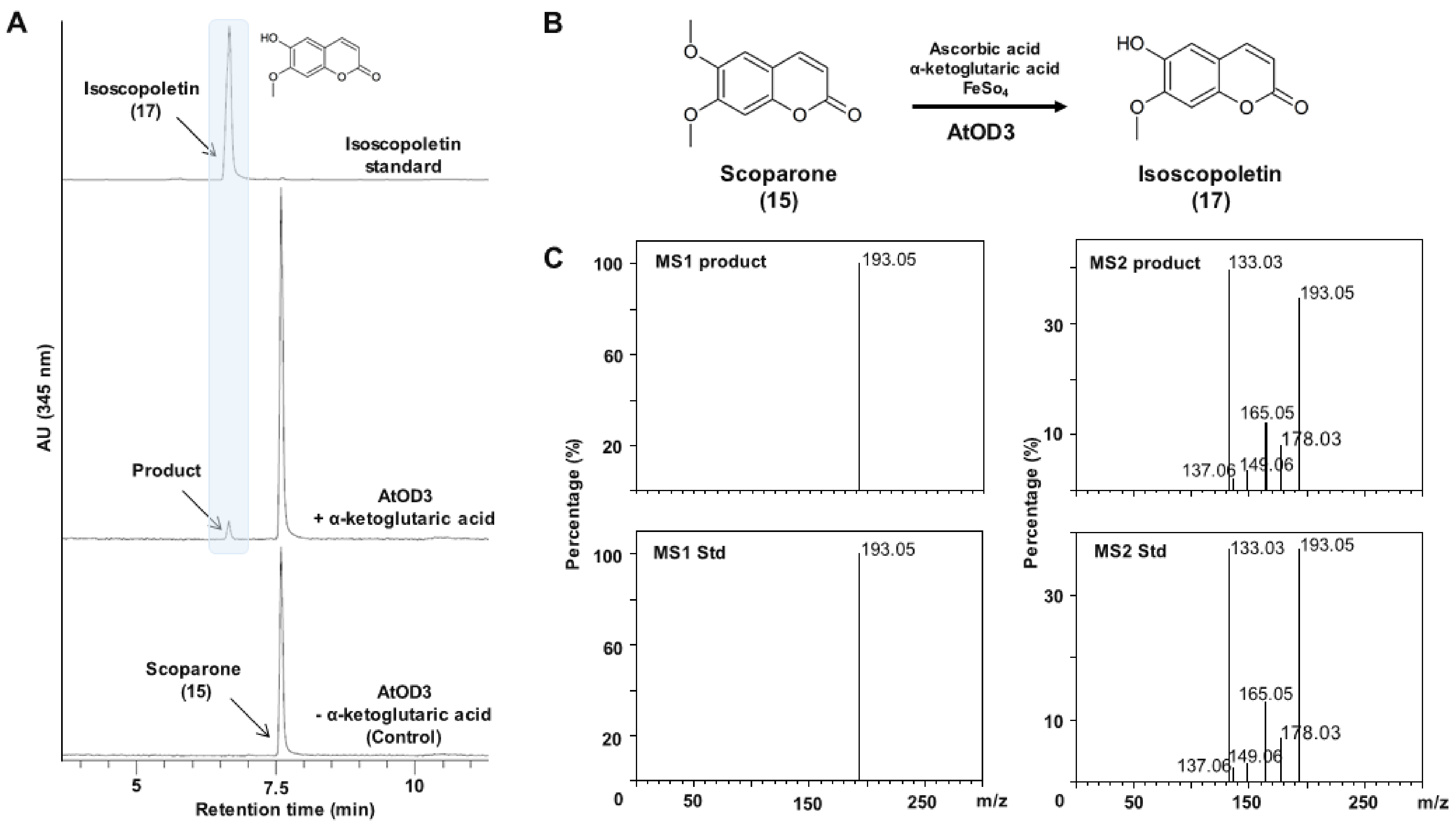


**Figure S20. Biochemical characterization of the *O*-demethylase activity of AtOD3 towards scoparone.** (A) UV chromatograms at 345 nm of enzymatic reactions using renatured sonication pellets containing recombinant AtOD3. Reaction mixtures (100 µL) contained purified enzyme, 500 µM α-ketoglutarate, 500 µM ascorbic acid, 5 mM $FeSO_4$, and 200 µM *O*-methylated substrate, and were incubated in a mixed buffer system (acetic acid 0.1 M, Tris 0.1 M, Imidazole 40 mM) under optimal temperature and pH conditions (Supplementary Figures X and Y) for 20 h. B) Schematic representation of the reaction converting scoparone to isoscopoletin. (C) Comparison of $MS^1$ and $MS^2$ spectra of the reaction product with those of authentic isoscopoletin standard in negative ion mode using Orbitrap detection.

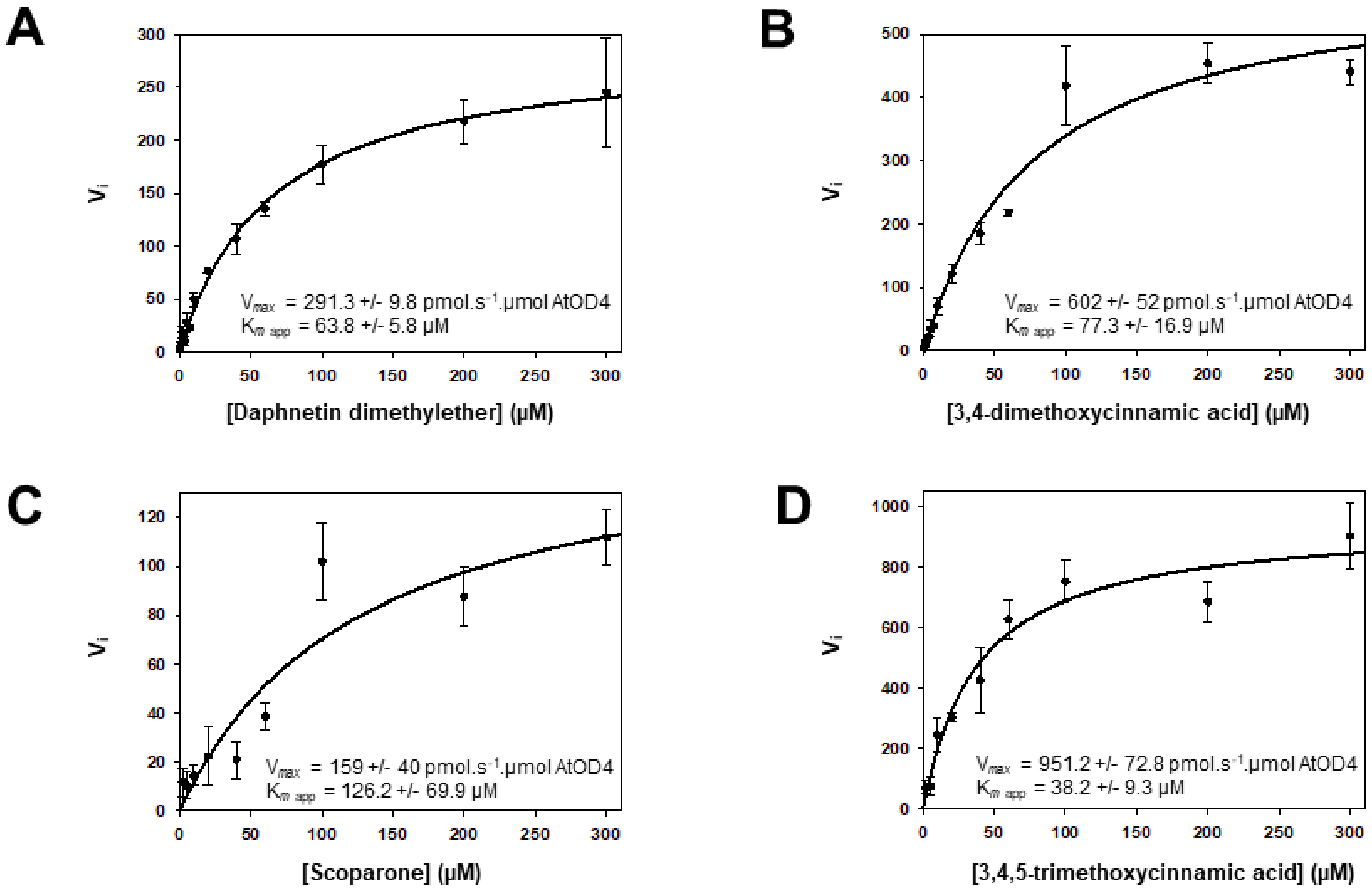


**Figure S21. Michaelis Menten regression of AtOD4 metabolization of A) daphnetin-dimethylether, B) 3,4-dimethoxycinnamic acid, C) scoparone and D) 3,4,5 trimethoxycinnamic acid substrates.** Reactions were carried out at optimal pH and temperature (Supplementary Fig. SX and SX) in polybuffer Tris 0.1 M acetic acid 0.1 M Imidazole 40 mM. The concentrations of substrates ranged from 1–300 µM for daphnetin dimethylether, scoparone, 3,4-dimethoxycoumarin and 3,4,5-trimethoxycoumarin. The concentration of α-ketoglutarate and ascorbic acid were set at 500 µM, the concentration of $FeSO_4$ was set at 5 mM. All results are expressed as means ± standard errors of three independent experiments. Figures presented were obtained by fitting the experimental data to the Michaelis-Menten equation.

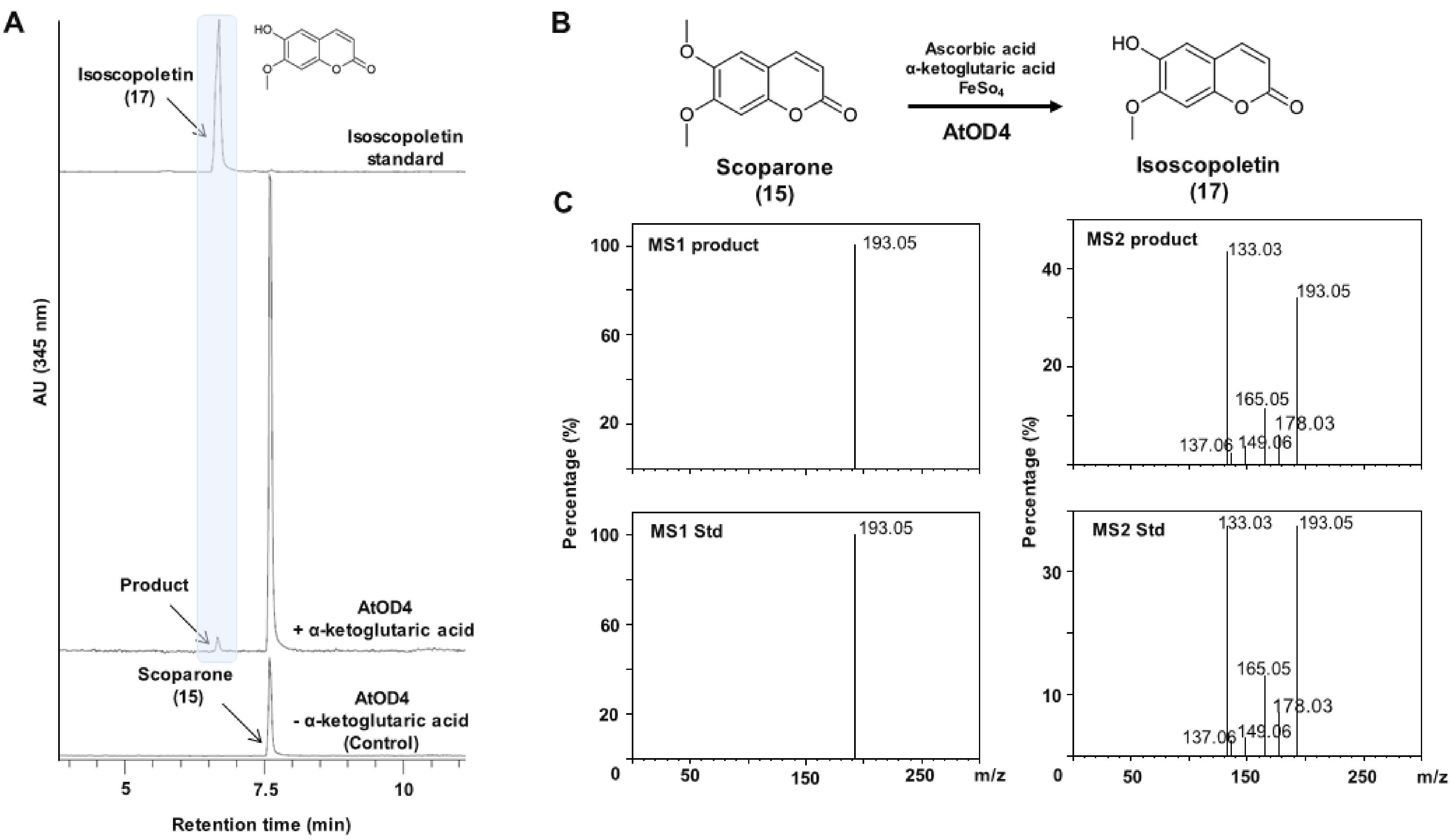


**Figure S22. Biochemical characterization of the *O*-demethylase activity of AtOD4 towards scoparone.** (A) UV chromatograms at 345 nm of enzymatic reactions using renatured sonication pellets containing recombinant AtOD4. Reaction mixtures (100 µL) contained purified enzyme, 500 µM α-ketoglutarate, 500 µM ascorbic acid, 5 mM $FeSO_4$, and 200 µM *O*-methylated substrate, and were incubated in a mixed buffer system (acetic acid 0.1 M, Tris 0.1 M, Imidazole 40 mM) under optimal temperature and pH conditions (Supplementary Figures X and Y) for 20 h. B) Schematic representation of the reaction converting scoparone to isoscopoletin. (C) Comparison of $MS^1$ and $MS^2$ spectra of the reaction product with those of authentic isoscopoletin standard in negative ion mode using Orbitrap detection.

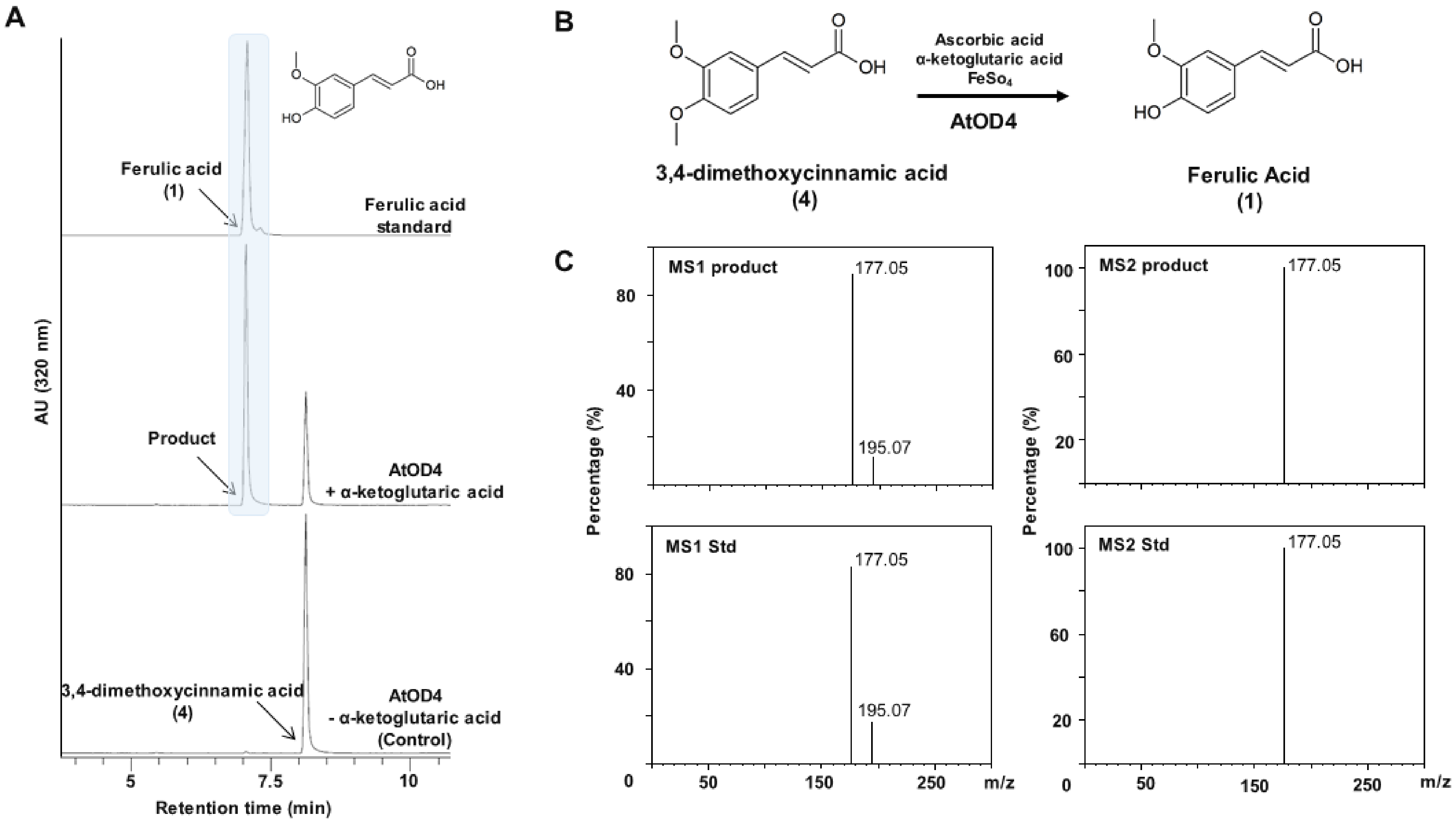


**Figure S23. Biochemical characterization of the *O*-demethylase activity of AtOD4 towards 3,4-dimethoxycinnamic acid.** (A) UV chromatograms at 320 nm of enzymatic reactions using renatured sonication pellets containing recombinant AtOD4. Reaction mixtures (100 µL) contained purified enzyme, 500 µM α-ketoglutarate, 500 µM ascorbic acid, 5 mM $FeSO_4$, and 200 µM *O*-methylated substrate, and were incubated in a mixed buffer system (acetic acid 0.1 M, Tris 0.1 M, Imidazole 40 mM) under optimal temperature and pH conditions (Supplementary Figures X and Y) for 20 h. B) Schematic representation of the reaction converting 3,4-dimethoxycinnamic acid to ferulic acid. (C) Comparison of $MS^1$ and $MS^2$ spectra of the reaction product with those of authentic ferulic acid standard in negative ion mode using Orbitrap detection.

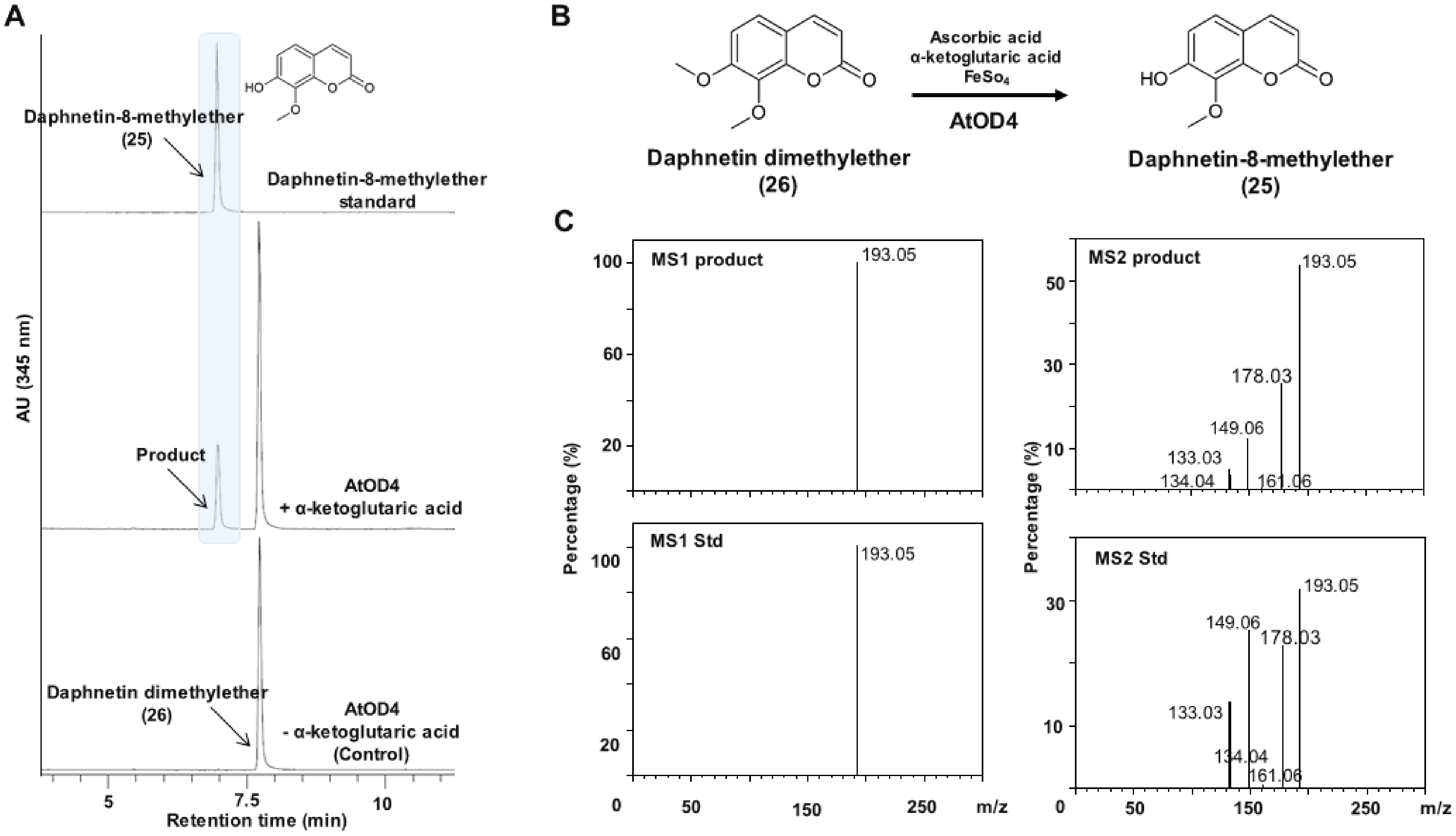


**Figure S24. Biochemical characterization of the *O*-demethylase activity of AtOD4 towards daphnetin dimethylether.** (A) UV chromatograms at 345 nm of enzymatic reactions using renatured sonication pellets containing recombinant S6OD1. Reaction mixtures (100 µL) contained purified enzyme, 500 µM α-ketoglutarate, 500 µM ascorbic acid, 5 mM $FeSO_4$, and 200 µM *O*-methylated substrate, and were incubated in a mixed buffer system (acetic acid 0.1 M, Tris 0.1 M, Imidazole 40 mM) under optimal temperature and pH conditions (Supplementary Figures X and Y) for 20 h. B) Schematic representation of the reaction converting daphnetin dimethylether to daphnetin-8-methylether. (C) Comparison of $MS^1$ and $MS^2$ spectra of the reaction product with those of authentic daphnetin-8-methylether standard in negative ion mode using Orbitrap detection.

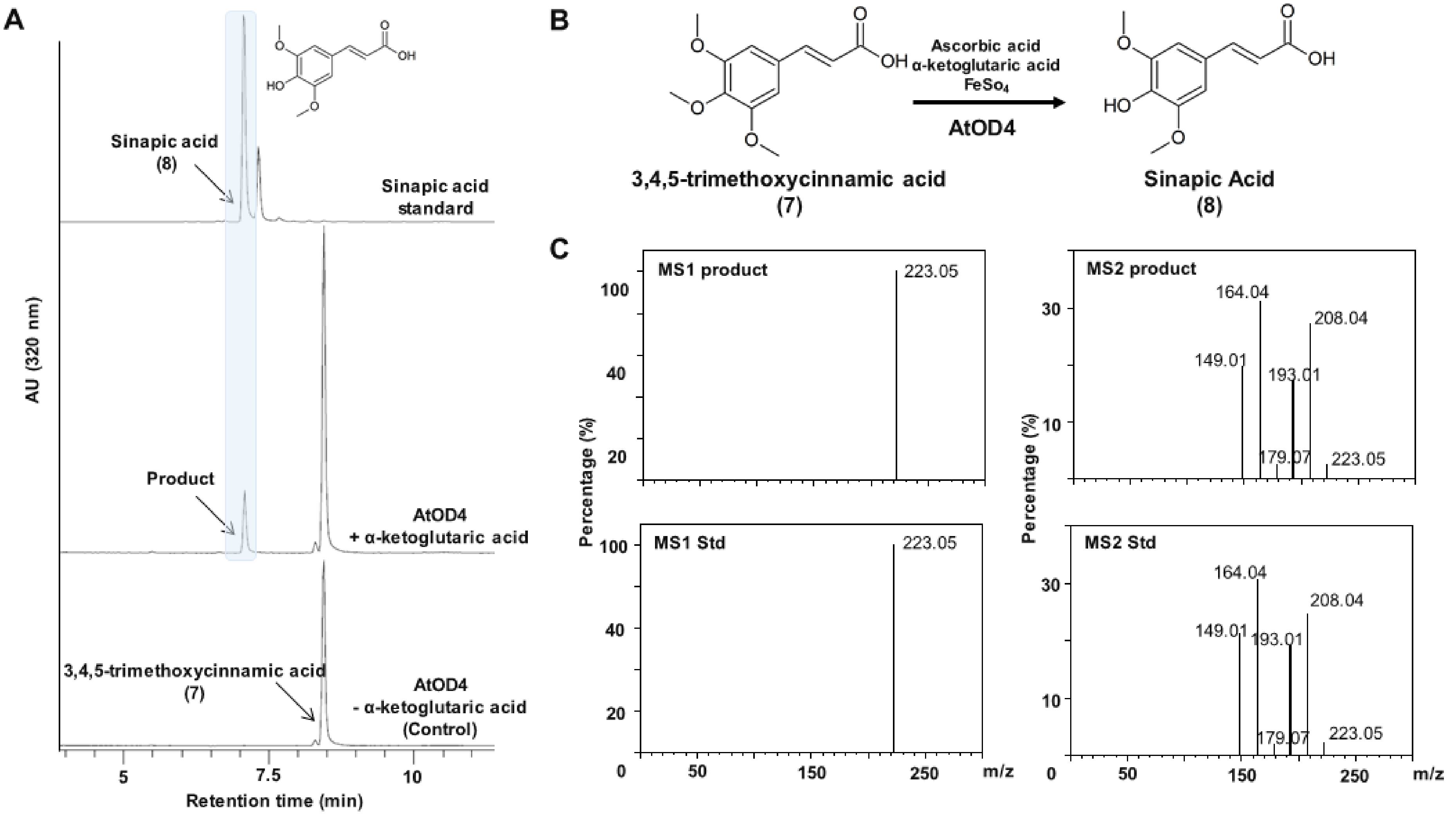


**Figure S25. Biochemical characterization of the *O*-demethylase activity of AtOD4 towards 3,4,5-trimethoxycinnamic acid.** (A) UV chromatograms at 320 nm of enzymatic reactions using renatured sonication pellets containing recombinant AtOD4. Reaction mixtures (100 µL) contained purified enzyme, 500 µM α-ketoglutarate, 500 µM ascorbic acid, 5 mM $FeSO_4$, and 200 µM *O*-methylated substrate, and were incubated in a mixed buffer system (acetic acid 0.1 M, Tris 0.1 M, Imidazole 40 mM) under optimal temperature and pH conditions (Supplementary Figures X and Y) for 20 h. B) Schematic representation of the reaction converting 3,4,5-trimethoxycinnamic acid to sinapic acid. (C) Comparison of $MS^1$ and $MS^2$ spectra of the reaction product with those of authentic sinapic acid standard in negative ion mode using Orbitrap detection.

**Table S4. PCR and RT-PCR conditions used for genotyping.**

*S6OD1 PCR genotyping reaction*

| Reaction components | | Temperature profile | | |
|---|---|---|---|---|
| **10x Buffer** | 2.5 µl | | | |
| **dNTPs (10mM)** | 0.5 µl | 95°C | | 3 min |
| **primer F (10mM)** | 1 µl | 35 cycles: | | |
| **primer R (10mM)** | 1 µl | | 94°C | 30 s |
| **DreamTaq Polymerase** | 0.25 µl | | 57°C | 30 s |
| **DNA** | 1 µl | | 72°C | 1 min |
| **$H_2O$** | 18.75 µl | 72°C | | 10 min |
| **total volume** | 25 µl | | | |

*S6OD2 PCR genotyping reaction*

| Reaction components | | Temperature profile | | |
|---|---|---|---|---|
| **10x Buffer** | 2.5 µl | | | |
| **dNTPs (10mM)** | 0.5 µl | 95°C | | 3 min |
| **primer F (10mM)** | 1 µl | 35 cycles: | | |
| **primer R (10mM)** | 1 µl | | 94°C | 30 s |
| **DreamTaq Polymerase** | 0.25 µl | | 55°C | 30 s |
| **DNA** | 1 µl | | 72°C | 1 min |
| **$H_2O$** | 18.75 µl | 72°C | | 7 min |
| **total volume** | 25 µl | | | |

*S6OD1 RT-PCR genotyping reaction*

| Reaction components | | Temperature profile | | |
|---|---|---|---|---|
| **10x Buffer** | 2.5 µl | | | |
| **dNTPs (10mM)** | 0.5 µl | 95°C | | 3 min |
| **primer F (10mM)** | 1 µl | 35 cycles: | | |
| **primer R (10mM)** | 1 µl | | 94°C | 30 s |
| **DreamTaq Polymerase** | 0.25 µl | | 48°C | 30 s |
| **cDNA** | 1 µl | | 72°C | 1 min |
| **$H_2O$** | 18.75 µl | 72°C | | 7 min |
| **total volume** | 25 µl | | | |

*S6OD2 RT-PCR genotyping reaction*

| Reaction components | | Temperature profile | | |
|---|---|---|---|---|
| **10x Buffer** | 2.5 µl | | | |
| **dNTPs (10mM)** | 0.5 µl | 95°C | | 3 min |
| **primer F (10mM)** | 1 µl | 34 cycles: | | |
| **primer R (10mM)** | 1 µl | | 94°C | 30 s |
| **DreamTaq Polymerase** | 0.25 µl | | 50°C | 30 s |
| **cDNA** | 1 µl | | 72°C | 1 min |
| **$H_2O$** | 18.75 µl | 72°C | | 7 min |
| **total volume** | 25 µl | | | |

**Table S5. Primers used for genotyping and transient expression in tobacco.**

| primer name | application | primer sequence (5'-3') |
|---|---|---|
| **ACT2 F** | RT-PCR | TCCCAGTGTTGTTGGTAGGC |
| **ACT2 R** | RT-PCR | CAAGACGGAGGATGGCATGA |
| **S6OD1 F1** | RT-CPR, PCR *s6od1-2* T-DNA | CCATGGAATGGACTTGGAC |
| **S6OD1 R1** | RT-PCR, cloning for transient expression | TTAGATTCTCATAACATCAAGG |
| **S6OD1 F2** | PCR *S6OD1* gene | CCATGGAATGGACTTGGAC |
| **S6OD1 R2** | PCR *S6OD1* gene, *s6od1-1* T-DNA | AGCTTTGAGCAACACTAACATCA |
| **SAIL T-DNA** | PCR *s6od1-1* T-DNA, *s6od2-2* T-DNA | GCCTTTTCAGAAATGGATAAATAGCCTTGCTTCC |
| **WISC T-DNA** | PCR *s6od1-2* T-DNA | AACGTCCGCAATGTGTTATTAAGTTGTC |
| **S6OD2 F** | PCR *S6OD2* gene, RT-PCR | GTTCCTTCTGTTCAGGAGAT |
| **S6OD2 R** | PCR *S6OD2* gene, PCR *S6OD2* T-DNA, RT-PCR | TCTTTATCCATTCCCGTGTTG |
| **SALK T-DNA** | PCR *s6od2-1* T-DNA | ATTTTGCCGATTTCGGAAC |
| **S6OD1 pCR8 F** | cloning for transient expression | ATGGAAGGTAAAGGAGTAACC |

A

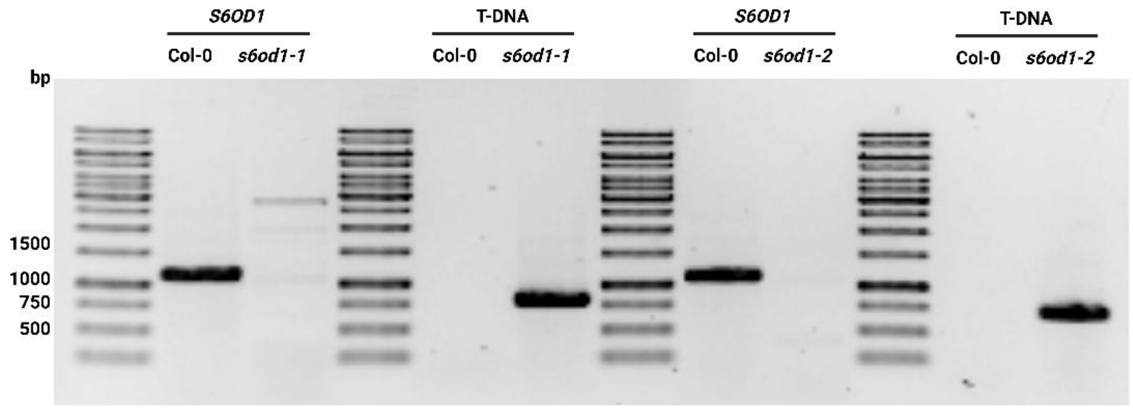


B

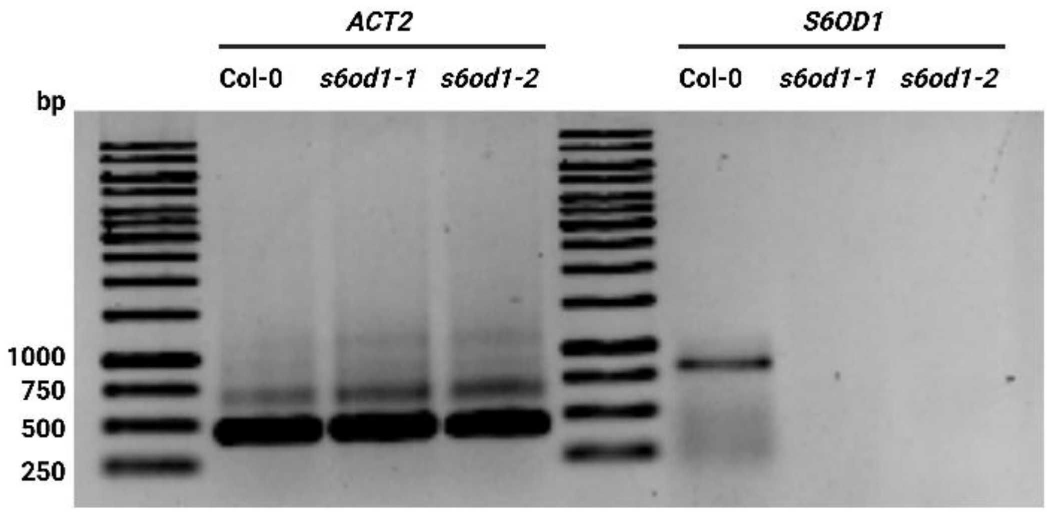


**Figure S26. Genomic (A) and transcript-level (B) confirmation of knockout in *s6od1* lines.** PCR with genomic DNA verified T-DNA insertion, and RT-PCR analysis confirmed loss of *S6OD1* transcripts. *ACTIN2* (*ACT2*) was used as an internal control.

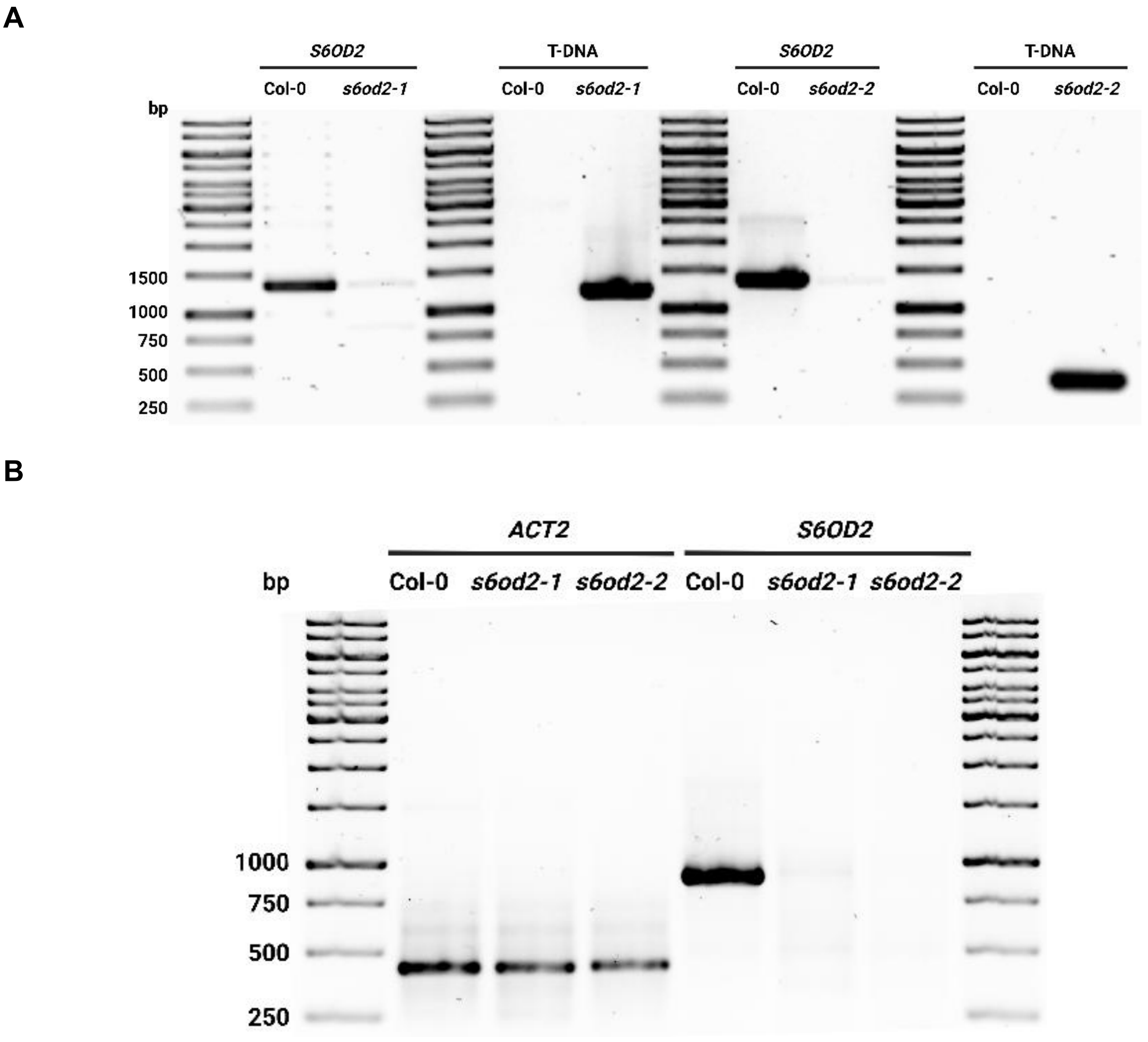


**Figure S27. Genomic (A) and transcript-level (B) confirmation of knockout in *s6od2* lines.** PCR with genomic DNA verified T-DNA insertion, and RT-PCR analysis confirmed loss of *S6OD2* transcripts. *ACTIN2* (*ACT2*) was used as an internal control.

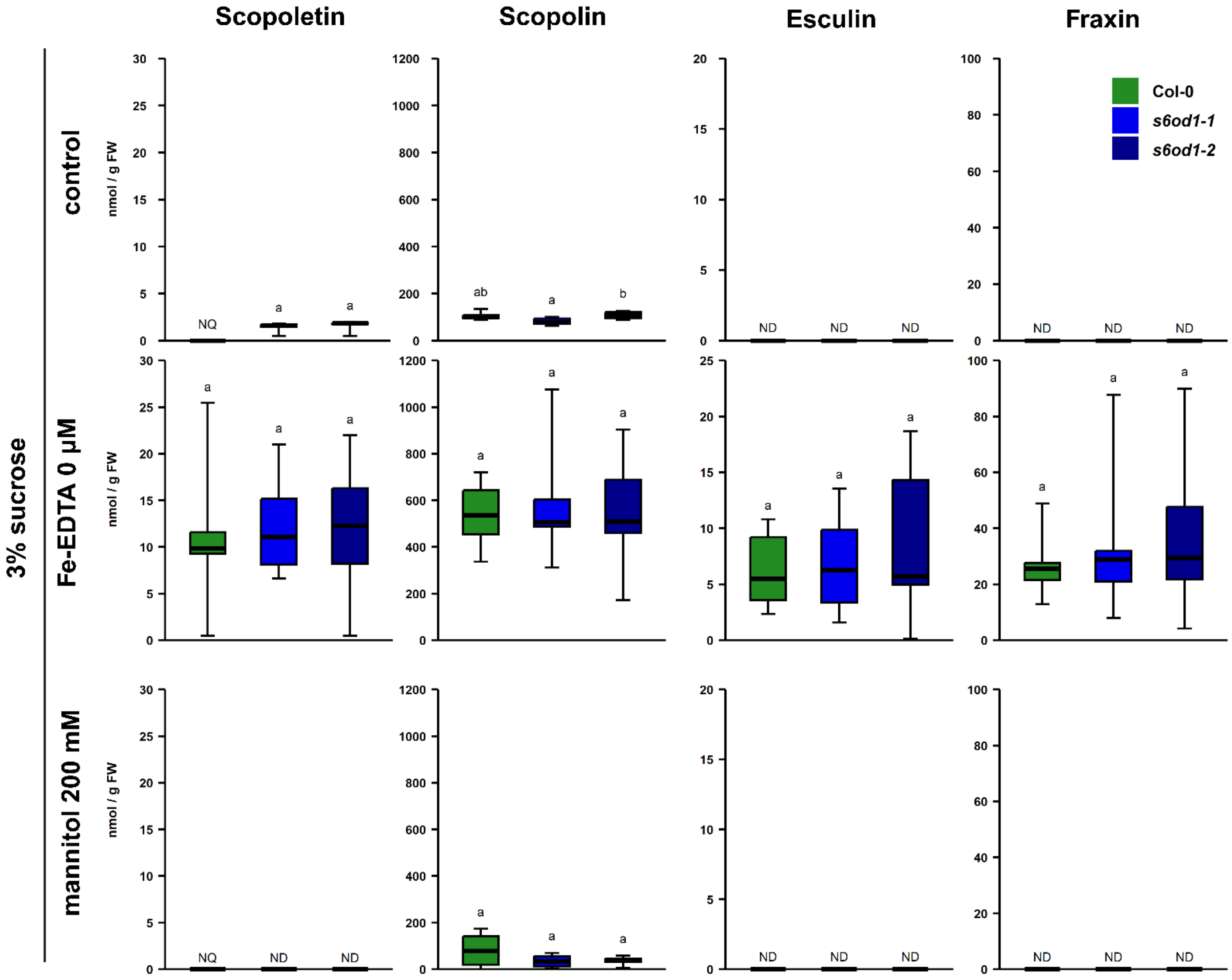


**Figure S28. Coumarin profiling of shoots of Col-0 and *s6od1*knockout mutant plants grown in liquid *in vitro* cultures in control, Fe deficiency and osmotic stress conditions.** Plants were grown on half-strength 1 % sucrose MS plates for 10 days, then transferred to half-strength 3% sucrose liquid media with 100 µM Fe-EDTA (control), 100 µM Fe-EDTA or 100 µM Fe-EDTA with 200 mM mannitol. ND – not detected, NQ – not quantifiable. n = 5 - 6. Statistical tests performed were ANOVA with Tukey's post hoc or Kruskal-Wallis test with Dunn's post hoc and Benjamini-Hochberg adjustment for multiple comparisons or two-tailed Student's *t*-test, depending on data.

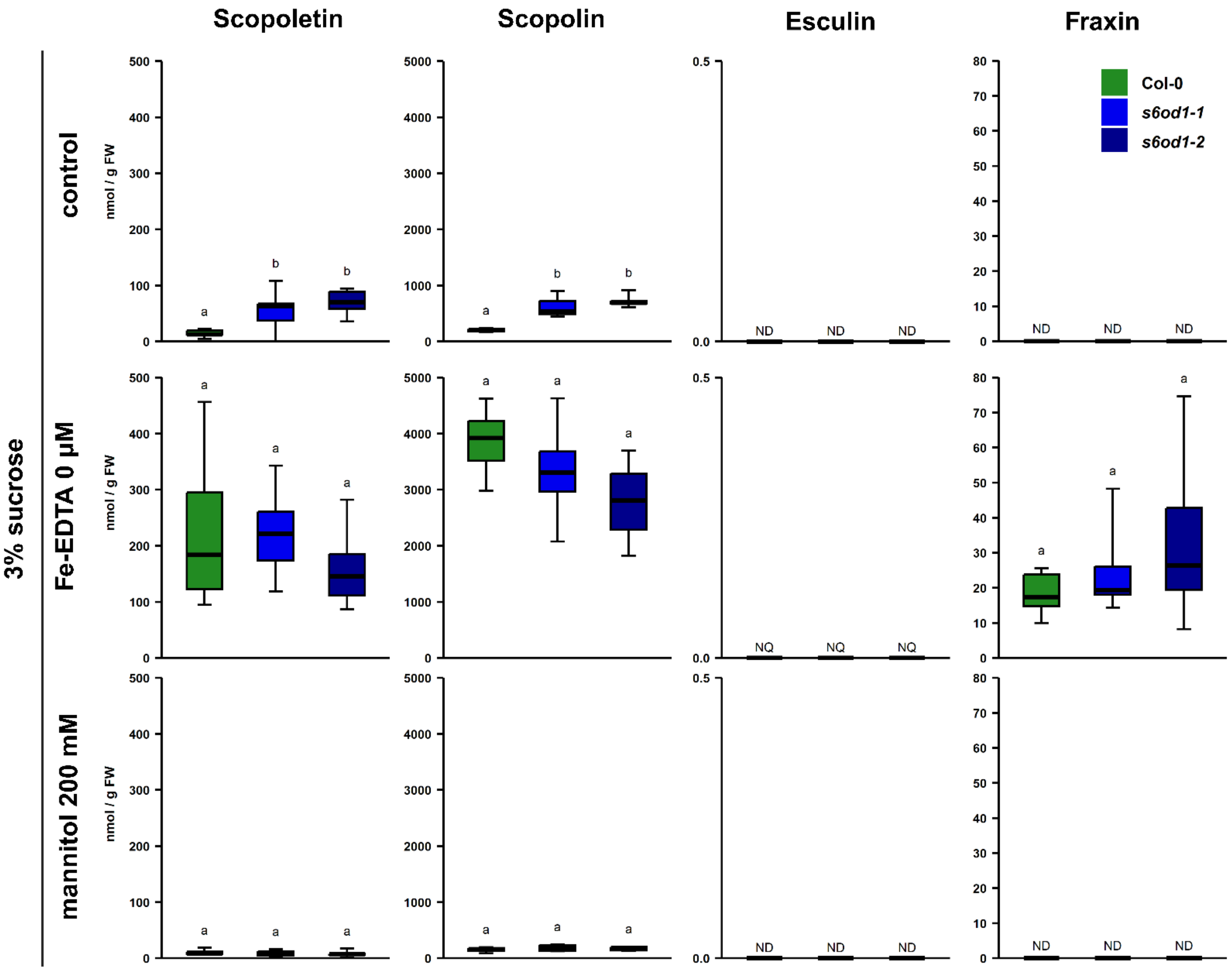


**Figure S29. Coumarin profiling of roots of Col-0 and *s6od1*knockout mutant plants grown in liquid *in vitro* cultures in control, Fe deficiency and osmotic stress conditions.** Plants were grown on half-strength 1 % sucrose MS plates for 10 days, then transferred to half-strength 3% sucrose liquid media with 100 µM Fe-EDTA (control), 100 µM Fe-EDTA or 100 µM Fe-EDTA with 200 mM mannitol. ND – not detected, NQ – not quantifiable. n = 5 - 6. Statistical tests performed were ANOVA with Tukey's post hoc or Kruskal-Wallis test with Dunn's post hoc and Benjamini-Hochberg adjustment for multiple comparisons, depending on data.

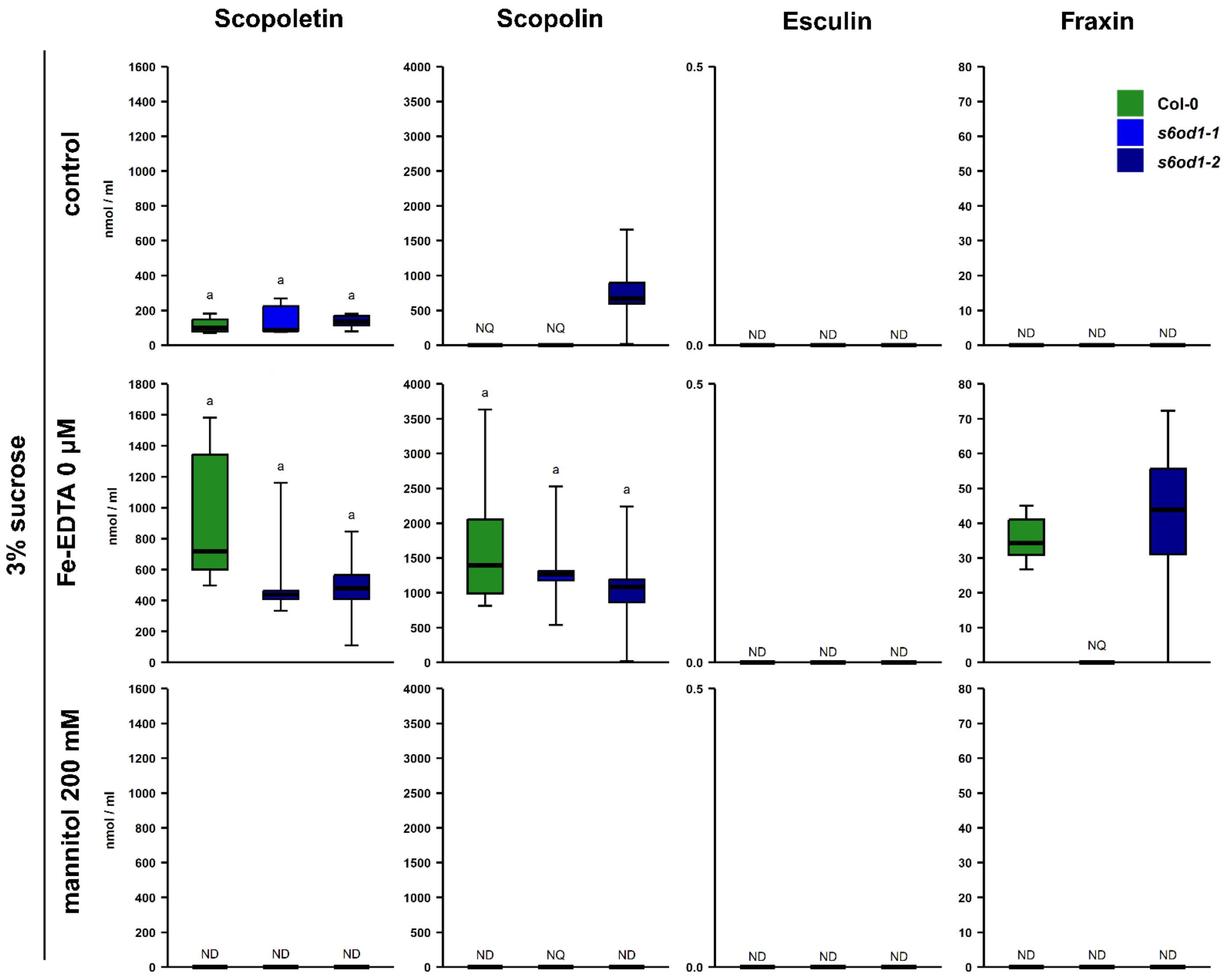


**Figure S30. Coumarin profiling of exudates of Col-0 and *s6od1*knockout mutant plants grown in liquid *in vitro* cultures in control, Fe deficiency and osmotic stress conditions.** Plants were grown on half-strength 1 % sucrose MS plates for 10 days, then transferred to half-strength 3% sucrose liquid media with 100 µM Fe-EDTA (control), 100 µM Fe-EDTA or 100 µM Fe-EDTA with 200 mM mannitol. ND – not detected, NQ – not quantifiable. n = 4 - 6. Statistical tests performed were ANOVA with Tukey's post hoc or Kruskal-Wallis test with Dunn's post hoc and Benjamini-Hochberg adjustment for multiple comparisons or two-tailed Student's *t*-test, depending on data.

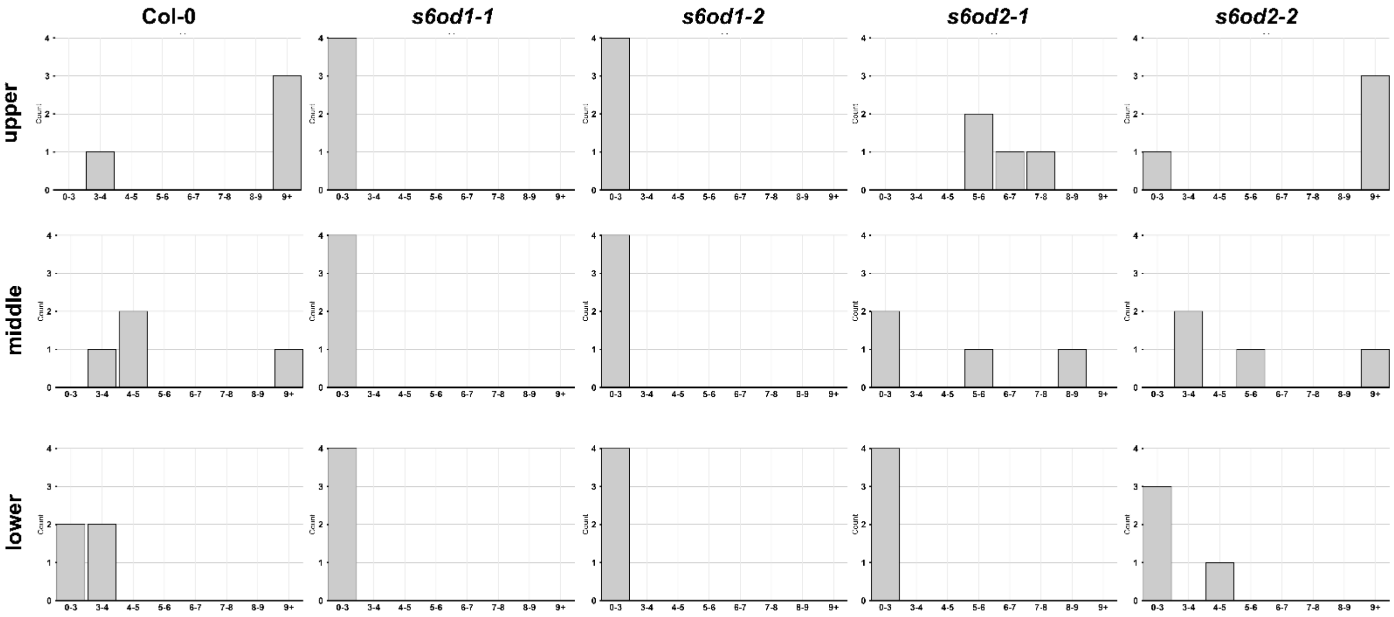


**Figure S31. Histograms showing distribution of signal to noise ratio (S/N) for esculetin peaks for Col-0, *s6od1* and *s6od2* mutant lines in different root parts.** Peaks with S/N < 3 were treated as non-detectable (ND), peaks with S/N ≥ 3 but < 9 were treated as non-quantifiable (NQ). Peaks with S/N ≥9 were treated as quantifiable. n = 4 (each consisting of root parts of 3 independently grown plants).

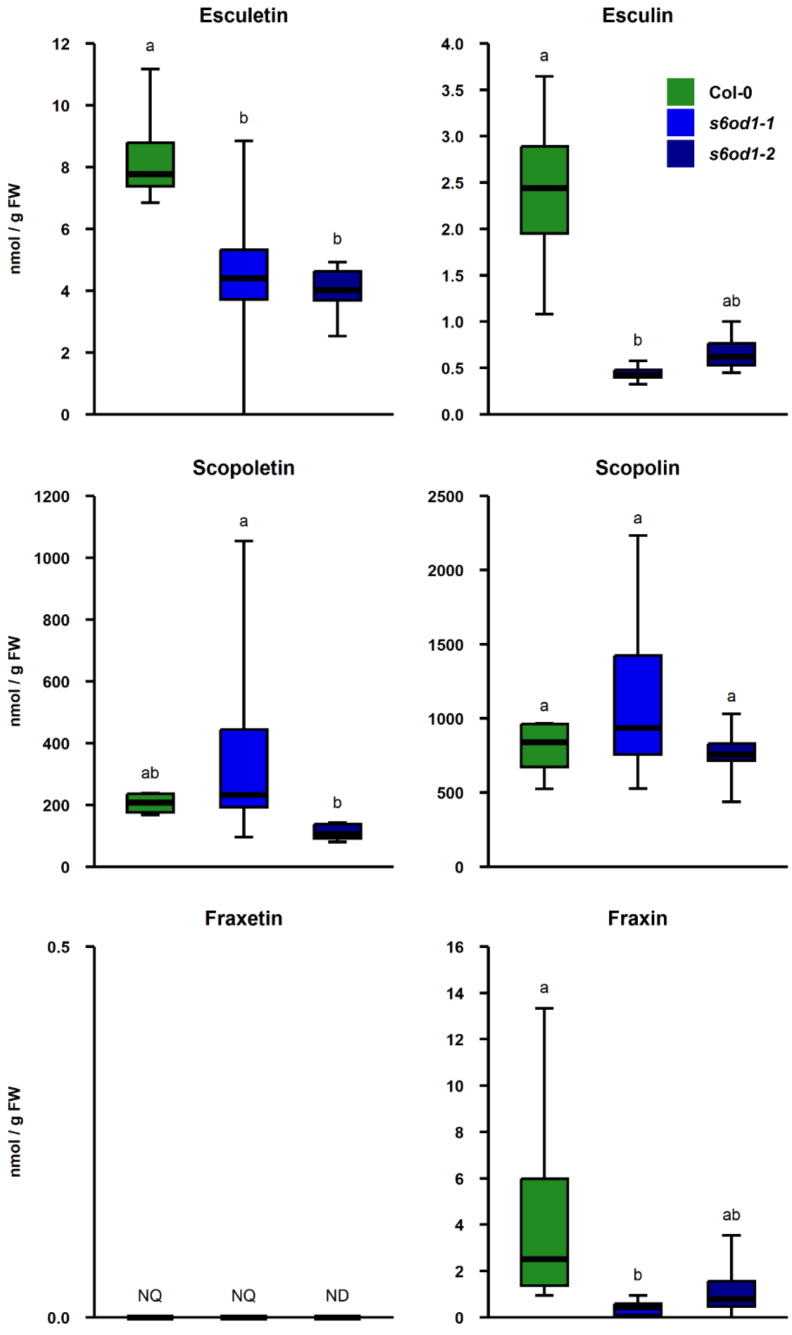


**Figure S32. Coumarin profiling of Col-0 and *s6od1*knockout mutant plants cultivated with 20 µM $FeCl_3$.** Plants were grown on half-strength MS plates for 4.5 weeks. ND – not detected, NQ – not quantifiable. n = 4 - 6, each consisting of roots of several plants grown on one or two plates Statistical tests performed were ANOVA with Tukey’s post hoc or Kruskal-Wallis test with Dunn’s post hoc and Benjamini-Hochberg adjustment for multiple comparisons, depending on data.

```
                     10        20        30        40        50        60        70        80
             ....|....|....|....|....|....|....|....|....|....|....|....|....|....|....|....|
Col-0 S6OD1  1   MEGKGVTFSSVIVPSVQEMVKEKVITTVLPPRYVRSDQEKGEAAIDSGENQIPIIDMSLLSSSTSMDSEIDKLDFACKEW
Stw-0 S6OD1  1   MEGKGVTFSSVIVPSVQEMVKEKVITTVLPPRYVRSDQEKGEAAIDSGLNQIPIIDMSLLSSSTSMDSEIDKLDFACKEW

                     90       100       110       120       130       140       150       160
             ....|....|....|....|....|....|....|....|....|....|....|....|....|....|....|....|
Col-0 S6OD1  81  GFFQLVNHGMDLDKFKSDIQDFFNLPMEEKKKLWQQPGDIEGFGQAFVFSEEQKLDWADVFFLTMQPVPLRKPHLFPKLP
Stw-0 S6OD1  81  GFFQLVNHGMDLDKFKSDIQDFFNLPMEEKKKLWQQPGDIEGFGQAFVFSEEQKLDWADVFFLTMQPVPLRKPHLFPKLP

                    170       180       190       200       210       220       230       240
             ....|....|....|....|....|....|....|....|....|....|....|....|....|....|....|....|
Col-0 S6OD1  161 LPFRDTLDTYSAELKSIAKVLFAKLASALKIKPEEMEKLFDDELGQRIRMNYYPPCPEPDKAIGLTPHSDATGLTILLQV
Stw-0 S6OD1  161 LPFRDTLDTYSAELKSIAKVLFAKLASALKIKPEEMEKLFDDELGQRIRMNYYPPCPEPDKAIGLTPHSDATGLTILLQV

                    250       260       270       280       290       300       310       320
             ....|....|....|....|....|....|....|....|....|....|....|....|....|....|....|....|
Col-0 S6OD1  241 NEVEGLQIKKDGKWVSVKPLPNALVVNVGDILEIITNGTYRSIEHRGVVNSEKERLSVASFHNTGFGKEIGPMRSLVERH
Stw-0 S6OD1  241 NEVEGLQIKKDGKWVSVKPLPNALVSMLETS

                    330       340       350
             ....|....|....|....|....|....|...
Col-0 S6OD1  321 KGALFKTLTTEEYFHGLFSRELDGKAYLDVMRI
Stw-0 S6OD1
```

**Figure S33. Amino acid sequence alignment of S6OD1 protein derived from Col-0 and Stw-0 natural accessions.** Stw-0 S6OD1 sequence ends prematurely, which is caused by a STOP codon resulting from a frameshift variant. Black background marks matching amino acids.

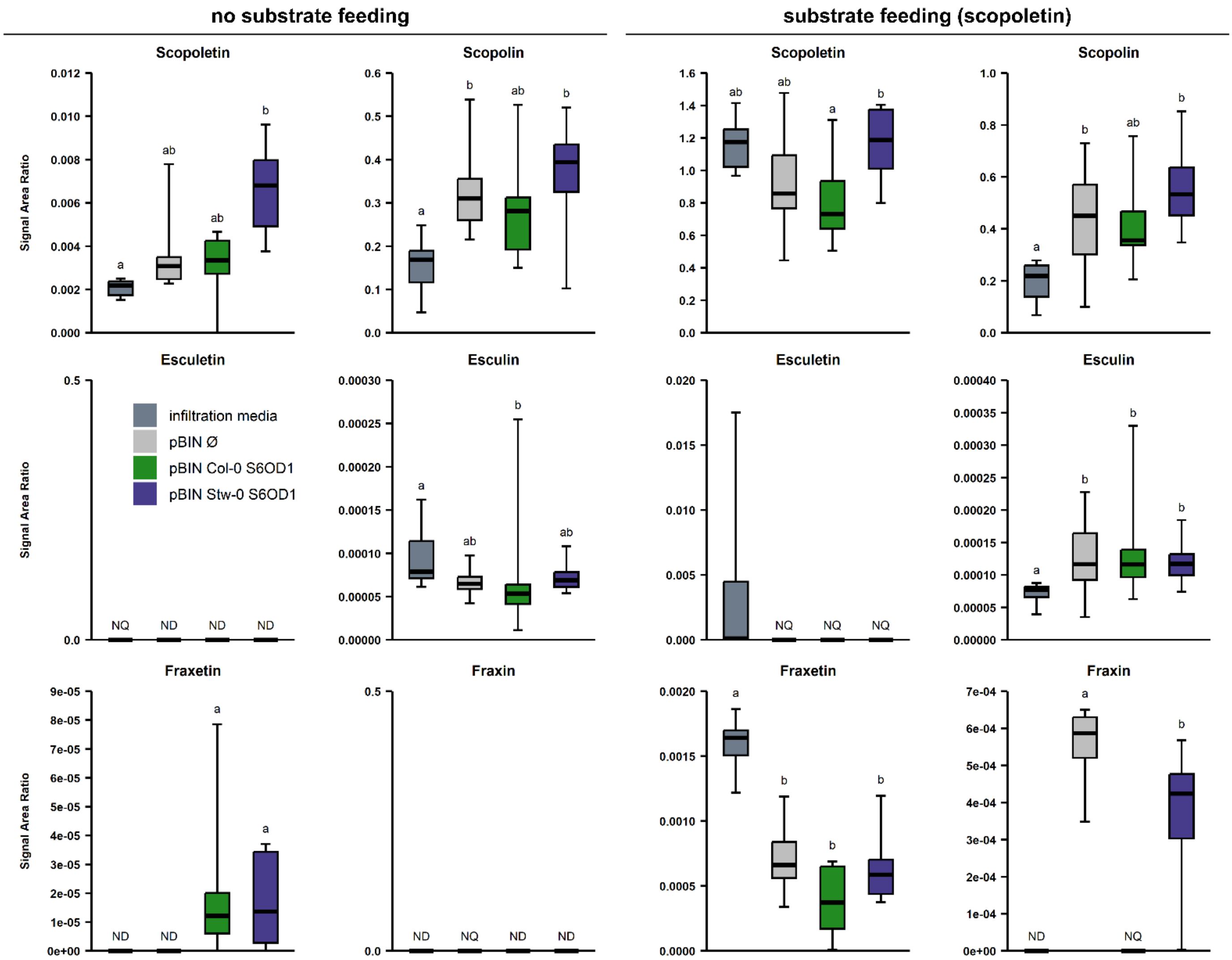


**Figure S34. Coumarin content in *Nicotiana benthamiana* leaves.** Transient expression was performed using *Agrobacterium tumefaciens* carrying pBIN empty vector as control, pBIN with *S6OD1* copy derived from Col-0 or Stw-0 accessions. Substrate feeding was carried out by infiltrating 500 µM scopoletin solution into the leaves two hours before harvesting samples. Results are shown as signal ratio of compound to external taxifolin standard. ND – not detected, NQ – not quantifiable. n = 8, each consisting of one or two leaf discs from a separate leaf. Statistical tests performed were ANOVA with Tukey's post hoc or Kruskal-Wallis test with Dunn's post hoc and Benjamini-Hochberg adjustment for multiple comparisons or two-tailed Student's *t*-test, depending on data

> P7ODM
MEKAKLMKLGNGLSIPSVQELAELTFAEVPSRYVCTNDENLLLMTMGASEIDDETVPVIDLQNLLSPEPAIGKSELDWLHYSCKEWGFF
QLVNHGVDALLVDHVKSEIHSFFNLPLNEKTKYGQRDGDVEGFGQAFLVSENQKLDWADMFFINTLPLHLRKPHLFPNLPLPLRETIES
YSSEMKKLSMVLFEMMGKAIEVIDIKEAITEMFEDGMQSMRMNYYPPCPQPERVIGITPHSDFDGLTILLQLNEVEGLQIRKEDKWISIK
PLPDAFIVNVGDIWEIMTNGVHRSVDHRGVINSTKERLSIATFHSPKLELEIGPISSLIRPETPAVFKSAGRFEDLLKEGLSRKLDGKSFLDC
MRM

> T6ODM
MEKAKLMKLGNGMEIPSVQELAKLTLAEIPSRYVCANENLLLPMGASVINDHETIPVIDIENLLSPEPIIGKLELDRLHFACKEWGFFQVV
NHGVDASLVDSVKSEIQGFFNLSMDEKTKYEQEDGDVEGFGQGFIESEDQTLDWADIFMMFTLPLHLRKPHLFSKLPVPLRETIESYSSE
MKKLSMVLFNKMEKALQVQAAEIKGMSEVFIDGTQAMRMNYYPPCPQPNLAIGLTSHSDFGGLTILLQINEVEGLQIKREGTWISVKPL
PNAFVVNVGDILEIMTNGIYHSVDHRAVVNSTNERLSIATFHDPSLESVIGPISSLITPETPALFKSGSTYGDLVEECKTRKLDGKSFLDSM
RI

> CODM
METPILIKLGNGLSIPSVQELAKLTLAEIPSRYTCTGESPLNNIGASVTDDETVPVIDLQNLLSPEPVVGKLELDKLHSACKEWGFFQLVN
HGVDALLMDNIKSEIKGFFNLPMNEKTKYGQQDGDFEGFGQPYIESEDQRLDWTEVFSMLSLPLHLRKPHLFPELPLPFRETLESYLSK
MKKLSTVVFEMLEKSLQLVEIKGMTDLFEDGLQTMRMNYYPPCPRPELVLGLTSHSDFSGLTILLQLNEVEGLQIRKEERWISIKPLPDA
FIVNVGDILEIMTNGIYRSVEHRAVVNSTKERLSIATFHDSKLESEIGPISSLVTPETPALFKRGRYEDILKENLSRKLDGKSFLDYMRM

> Basil O demethylase
MRITLQYIKLESKNTKERDMAESKAIGRSLEVPNVQELAKGKLASVPARYVRYSDRENTTLPPLTQIPVIDMQALLHPNSFEAELNSLHK
ACKQWGFFQLINHGVEAAVMEKMKLEMQEFFNLPLEEKQKFRQSADDMEGYGQSFVVSDEQKLDWADGFSVISLPTYLRKPHLIPKL
PAPFRDAIDAYGAQLKELAIKILGFMAEALGMDPHEMTALFEEGIQALRMNYYPPCPQPEMVSGLCPHSDAGGLTILMQVNEVEGLQV
RKDGGWVPVSPLPDAFIINLGDILEIVTNGEYFSVEHQATVNGDKERLSVAAFLNPKMEDNIGPAASFISGETPAKFKTITAAEYFKGLFS
KELDGKSYLDLMRIQN

> At1g17010 S6OD2
MEAKEETPWSSILVPSVQEMVKDKMITTVPPRYVRYDQDKTEVVVHDSGLSEIPIIDMNRLCSSTAVDSEVEKLDFACKEYGFFQLVNH
GIDPSFLDKIKSEIQDFFNLPMEEKKKLWQTPAVMEGFGQAFVVSEDQKLDWADLFFLIMQPVQLRKRHLFPKLPLPFRDTLDMYSTRV
KSIAKILLAKMAKALQIKPEEVEEIFGDDMMQSMRMNYYPPCPQPNLVTGLIPHSDAVGLTILLQVNEVDGLQIKKNGKWFFVKPLQNA
FIVNVGDVLEIITNGTYRSIEHRAMVNLEKERLSIATFHNTGMDKEIGPARSLVQRQEAAKFRSLKTKDYLNGLFSRELKGKAYLDAMRI
EGK

> At1g17020 AtOD3
MEAKGAAQWSSILVPSVQEMVKEKTITTVPPRYVRSDQDKTEVDDDFDVKEIPIIDMKRLCSSTTMDSEVEKLDFACKEWGFFQLVNH
GIDSSFLDKVKSEIQDFFNLPMEEKKKFWQRPDEIEGFGQAFVVSEDQKLDWADLFFHTVQPVELRKPHLFPKLPLPFRDTLEMYSSEV
QSVAKILIAKMARALEIKPEELEKLFDDVDSVQSMRMNYYPPCPQPDQVIGLTPHSDSVGLTVLMQVNDVEGLQIKKDGKWVPVKPLP
NAFIVNIGDVLEIITNGTYRSIEHRGVVNSEKERLSIATFHNVGMYKEVGPAKSLVERQKVARFKRLTMKEYNDGLFSRTLDGKAYLDAL
RI

> At4g25300 AtOD4
MEVKGATRSSIIVPSVQEMVKEKMITTVPPRYVRSDQDVAEIAVDSGLRNQIPIIDMSLLCSSTSMDSEIDKLDSACKEWGFFQLVNHGM
ESSFLNKVKSEVQDFFNLPMEEKKNLWQQPDEIEGFGQVFVVSEEQKLDWADMFFLTMQPVRLRKPHLFPKLPLPFRDTLDMYSAEVK
SIAKILLGKIAVALKIKPEEMDKLFDDELGQRIRLNYYPRCPEPDKVIGLTPHSDSTGLTILLQANEVEGLQIKKNAKWVSVKPLPNALVV
NVGDILEIITNGTYRSIEHRGVVNSEKERLSVAAFHNIGLGKEIGPMRSLVERHKAAFFKSVTTEEYFNGLFSRELDGKAYLDVMRL

> At4g25310 S6OD1
MEGKGVTFSSVIVPSVQEMVKEKVITTVLPPRYVRSDQEKGEAAIDSGENQIPIIDMSLLSSSTSMDSEIDKLDFACKEWGFFQLVNHGM
DLDKFKSDIQDFFNLPMEEKKKLWQQPGDIEGFGQAFVFSEEQKLDWADVFFLTMQPVPLRKPHLFPKLPLPFRDTLDTYSAELKSIAK
VLFAKLASALKIKPEEMEKLFDDELGQRIRMNYYPPCPEPDKAIGLTPHSDATGLTILLQVNEVEGLQIKKDGKWVSVKPLPNALVVNV
GDILEIITNGTYRSIEHRGVVNSEKERLSVASFHNTGFGKEIGPMRSLVERHKGALFKTLTTEEYFHGLFSRELDGKAYLDVMRI

> At1g78550 AtOD5
MEAEGEKQWSSLIVPFVLEIVKEKNFTTIPPRYVRVDQEKTEILNDSSLSSEIPVIDMTRLCSVSAMDSELKKLDFACQDWGFFQLVNHG
IDSSFLEKLETEVQEFFNLPMKEKQKLWQRSGEFEGFGQVNIVSENQKLDWGDMFILTTEPIRSRKSHLFSKLPPPFRETLETYSSEVKSI
AKILFAKMASVLEIKHEEMEDLFDDVWQSIKINYYPPCPQPDQVMGLTQHSDAAGLTILLQVNQVEGLQIKKDGKWVVVKPLRDALV
VNVGEILEIITNGRYRSIEHRAVVNSEKERLSVAMFHSPGKETIIRPAKSLVDRQKQCLFKSMSTQEYFDAFFTQKLNGKSHLDLMRI

**Figure S35.** Primary structures of enzymatically *Opium poppy* and Basil *O*-demethylases and DOXC clade of Arabidopsis.

**Table S6. Nucleotide sequence of the primers used to amplify S6OD1 to AtOD5.**

| Primer name | Sequence (5'–3') |
|---|---|
| **AT4G25310 (S6OD1) Fwd** | ATGGAAGGTAAAGGAGTAACCTTTAGTTCTG |
| **AT4G25310 (S6OD1) Rev** | TTAGATTCTCATAACATCAAGGTAAGCTTTTCCATC |
| **AT1G17010 (S6OD2) Fwd** | ATGGAAGCCAAAAGAAGAACCCCGTGG |
| **AT1G17010 (S6OD2) Rev** | TCACTTCCCTTCAATTCTCATAGCATCAAGATAAGC |
| **AT1G17020 (AtOD3) Fwd** | ATGGAAGCAAAAGGGGCAGCACAGTG |
| **AT1G17020 (AtOD3) Rev** | TTAGATTCTCAAAGCATCTAGATAAGCTTTTCCGTC |
| **AT4G25300 (AtOD4) Fwd** | ATGGAAGTCAAAGGAGCAACCCGGAG |
| **AT4G25300 (AtOD4) Rev** | TTAGAGTCTCATAACATCAAGGTAAGCTTTTCCATC |
| **AT1G78550 (AtOD5) Fwd** | ATGGAGGCGGAAGGAGAAAAACAGTG |
| **AT1G78550 (AtOD5) Rev** | TTAAATACGCATAAGATCAAGGTGAGATTTCCC |

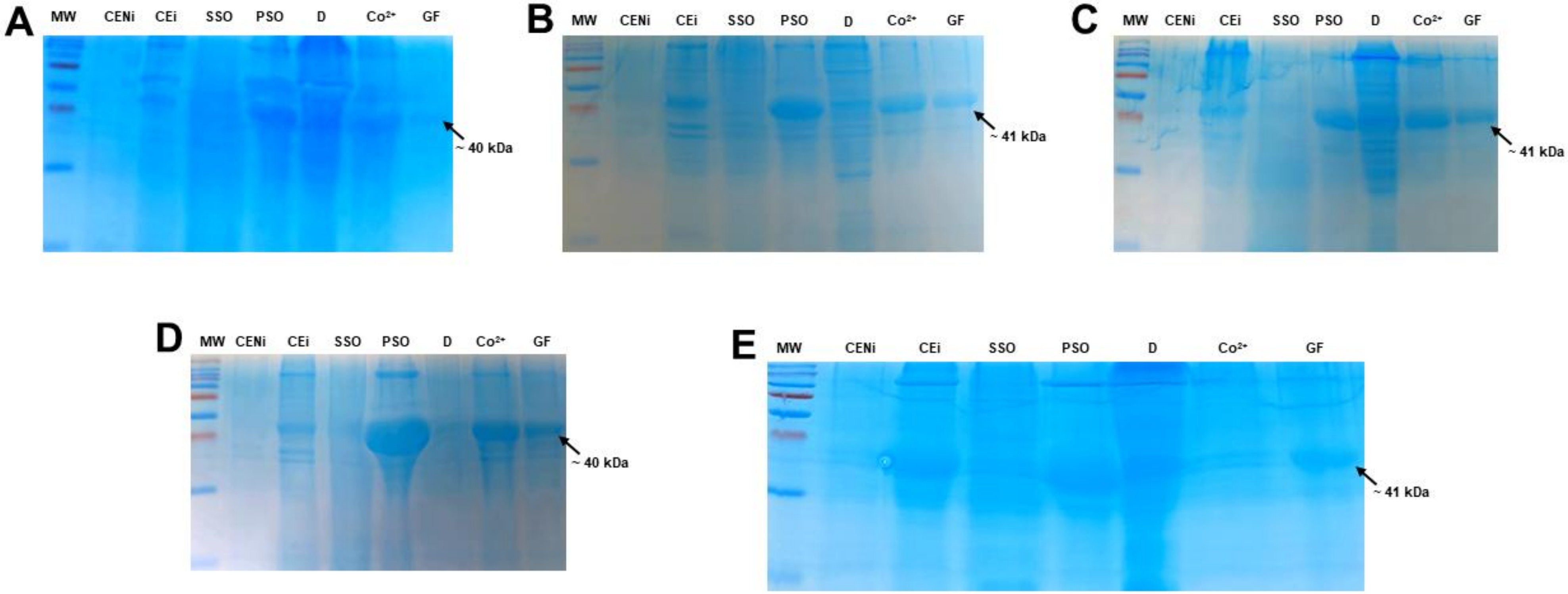


**Figure S36. SDS-PAGE of recombinants His-tagged *O*-demethylases proteins purification. A: S6OD1. B: S6OD2. C: AtOD3. D: AtOD4. E: AtOD5.** The purification of renaturated recombinant his-tagged proteins was done using the HIS-Select® Cobalt Affinity Gel (Sigma-Merck) and Sephadex® G-50 (Sigma-Merck) as described by the supplier. The predicted molecular weight of the his-tagged protein is ~ 41 kDa for all enzymes. (MW) molecular weight marker; (CENi) crude extracts, non-induced cultures; (CEi) crude extracts, induced cultures; (SSO) sonification supernatant; (PSO) sonification pellet; (D) Dialysate renaturated enzyme, ($Co^{2+}$) Eluted $Co^{2+}$ column; (GF) Gel filtration purified protein.
