## SUPPLEMENTARY FILE 2 for "First *O*-demethylation activity in Arabidopsis specialized metabolism resolves the missing step in esculetin biosynthesis"

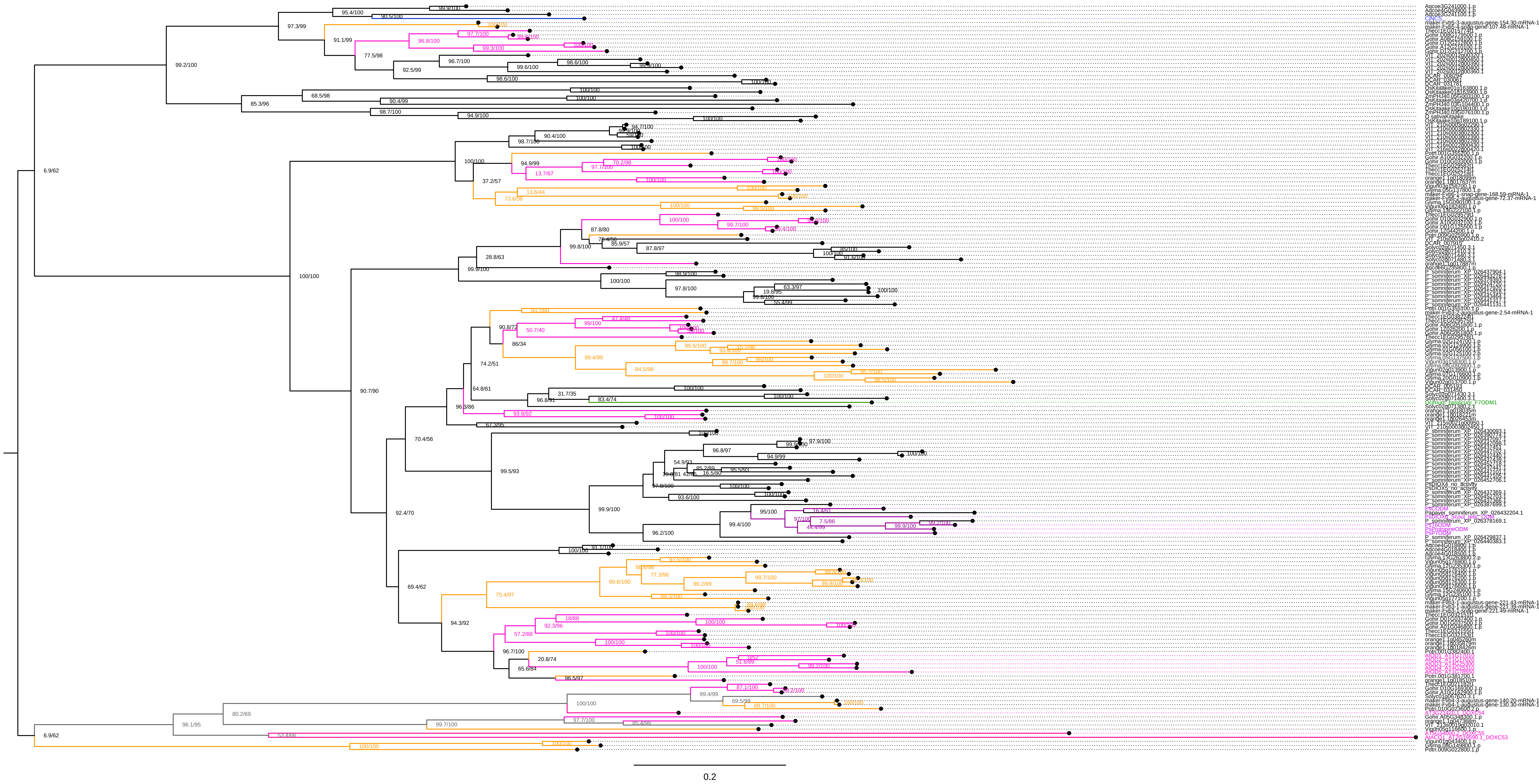
0.2

**Supplementary File 2. A detailed phylogenetic tree of the DOXC52 subfamily.** A maximum likelihood-based phylogenetic tree was constructed for the DOXC52 subfamily using protein sequences belonging to DOXC53, 54, and 55 as outgroups, based on MAFFT alignment and IQ-TREE inference (JTT+F+I+G4 model). The results of the Shimodaira-Hasegawa approximate likelihood-ratio test (1,000 replicates, left) and the ultrafast bootstrap test (1,000 replicates, right) are given for each node. The scale bar indicates an amino acid substitution rate per site of 0.20. The proteins from Fabids and Malvids are highlighted by orange and magenta, respectively. The previously-reported plant *O*-demethylases from *Papaver somniferum* and basil (*Ocimum basilicum*), as well as norcoclaurine synthase catalyzing the condensation of dopamine and 4- hydroxyphenylacetaldehyde to form norcoclaurin in *Coptis japonica*, are shown in purple, green, and blue, respectively. Except for these three species, the correspondence table between the protein name and the plant species are given in **Supplementary File 1.**
